# The NOD/RIPK2 signaling pathway contributes to osteoarthritis susceptibility

**DOI:** 10.1101/2022.02.07.479420

**Authors:** Michael J. Jurynec, Catherine M. Gavile, Matthew Honeggar, Ying Ma, Shivakumar R. Veerabhadraiah, Kendra A. Novak, Kazuyuki Hoshijima, Nikolas H. Kazmers, David J. Grunwald

## Abstract

Osteoarthritis (OA) is a debilitating disease characterized by loss of homeostasis of the joint with consequent remodeling of tissue architecture^1^. The molecular pathways that limit disease onset or progression are unknown^2-5^, and consequently no disease-modifying drugs are available^1,6-9^. We sought genes that contribute to dominant forms of hereditary OA with the aim of identifying pathways whose activity level contributes to OA susceptibility. We found seven independent alleles affecting the NOD/RIPK2 pathway. To determine if altered signaling is sufficient to confer heightened OA susceptibility, mice carrying the OA-associated hyperactive *Ripk2*^*104Asp*^ allele were generated. Knees of heterozygous *Ripk2*^*104Asp*^ mice exhibit no overt signs of joint remodeling. Nevertheless, the mice respond to injury with markedly advanced post-traumatic OA. Uninjured heterozygous *Ripk2*^*104Asp*^ mice appear primed to develop OA: their knees exhibit elevated NOD/RIPK2 pathway activity, localized inflammation, and altered expression of extracellular matrix genes linked to OA. In contrast to the joint, the mice display no evidence of systemic elevated inflammation. Elevated NOD/RIPK2 signaling confers vulnerability to OA.

## Main

The NOD/RIPK2 signaling pathway is a key arm of the innate immunity system, playing critical roles both in clearing bacterial infections and maintaining immune homeostasis^10^. The intracellular nucleotide-binding oligomerization domain (NOD) receptors are activated by bacterial cell wall breakdown products and additional damage-associated molecular patterns^10,11^. Activated NOD receptors signal through the Receptor Interacting Protein Kinase 2 (RIPK2), stimulating the MAPK and NF-κB pathways to elicit tissue-specific responses, most notably inflammatory responses^12-15^. NOD/RIPK2 signaling is tightly regulated, as mutations that either abrogate or elevate signaling are associated with chronic inflammatory diseases, including Crohn’s, Blau syndrome, early-onset sarcoidosis, and Behcet’s disease^10,16,17^. Although chronic inflammatory diseases are often associated with arthritis, no single inflammatory pathway has yet been linked to classic non-syndromic forms of OA^8,18^.

Identification of genes underlying familial forms of disease has uncovered pathways that confer elevated disease susceptibility^19-22^, but few studies have focused on the genetics of non-syndromic familial OA^23-26^. Previously we reported a rare dominant allele of the *RIPK2* gene (p.Asn104Asp) that co-segregated with OA of the first metatarsophalangeal (1^st^ MTP) joint in a family^23^. The OA-associated *RIPK2*^*104Asp*^ allele encodes a substitution for a conserved amino acid in the kinase domain^13^, producing a protein with heightened ability to activate the innate immune response and the NF-κB pathway^23^. We took an unbiased genetic approach to determine if the NOD/RIPK2 signaling pathway has a strong effect on OA susceptibility. We assembled a cohort of 150 independent OA families with a dominant inheritance pattern of OA. Each family is characterized by disease affecting distinct subsets of joints: distal and proximal interphalangeal OA^27^, glenohumeral OA^28^, or 1^st^ MTP joint OA^29,30^. Whole exome sequence (WES) analysis was performed on informative members of families, and coding variants that invariably segregated with OA and were predicted to alter gene function were identified^23^.

Six novel alleles affecting five NOD/RIPK2 pathway genes were found associated with OA in the new cohort of families (Table 1 and Supplementary Table 1, Extended Data Fig.1). Consistent with a dominant pattern of inheritance with strong penetrance, the variants are rare in human populations. Three variants affected the *NOD1* and *NOD2* genes, which encode intracellular receptors that function as upstream activators of RIPK2. The amino acid substitutions encoded by two of the variants reside within the autoinhibitory domain of the respective NOD protein^31^. Candidate variants in three additional families affected genes known to modify activity of the NOD/RIPK2 pathway, *CARD9, CHUK*, and *IKBKB*^32-35^. The family studies indicate a striking correlation between inheritance of variants that alter conserved sites within proteins of the NOD/RIPK2 signaling pathway and the occurrence of disease within families exhibiting OA of the hand, 1^st^ MTP joint, or shoulder.

**Table 1.** *NOD/RIPK2* Pathway Variants Identified in Independent Osteoarthritis Families.

| Gene | OA Phenotype (Family) | Variant | Minor Allele Frequency | Protein Domain Affected by Variant |
| --- | --- | --- | --- | --- |
| *NOD1* | Finger Interphalangeal Joint OA (FIJ744) | c.G2114A:p.R705Q | 0.0008 | Leucine Rich Repeat Domain |
| *NOD2* | 1st MTP Joint OA (UUHR2) | c.C2546T:p.A849V | 0.00007 | Leucine Rich Repeat Domain |
| *NOD2* | Finger Interphalangeal Joint OA (FIJ7) | c.G247A:p.A83T | 0.00008 | Caspase Activation and Recruitment Domain |
| *IKBKB* | Glenohumeral OA (SA735) | c.G1663A:p.G555R | 0.00008 | Scaffold Dimerization Domain |
| *CARD9* | Finger Interphalangeal Joint OA (FIJ9) | c.G722A:p.R241Q | 0.00005 | Structural Maintenance of Chromosomes |
| *CHUK* | 1st MTP Joint OA (MTP25) | c.A376T:p.S126C | 0.0008 | Kinase Domain |
| *RIPK2\** | 1st MTP Joint OA (UUHR1) | c.A310G:p.N104D | 0.0004 | Kinase Domain |

\* - Previously described in Juryneć, 2018<sup>22</sup>.

In the mouse, *Nod1/2* and *Ripk2* are expressed in uninjured joints, and the pathway is activated following injury (Extended Data Fig. 2). To determine whether altered pathway signaling is sufficient to confer increased susceptibility to OA, we used precise genome editing to introduce the human *RIPK2*^*104Asp*^ variant into the C57BL/6J inbred strain of mice, thereby creating an isogenic pair of mouse lines: the parental strain, which encodes the mammalian lineage-conserved Asn at position 104 (WT), and a derived line whose *Ripk2* allele encodes Asp at position 104 (*Ripk2*^*104Asp*^) (Fig. 1a). The modified allele is expressed at WT levels (Fig. 1b and Extended Data Fig. 3). Homozygous and heterozygous mice carrying the *Ripk2*^*104Asp*^ allele are viable and display no overt phenotypes. To recapitulate the dominant human phenotype^23^, heterozygous *Ripk2*^*104Asp*^ mice were used for all subsequent analyses.

**Figure 1.**
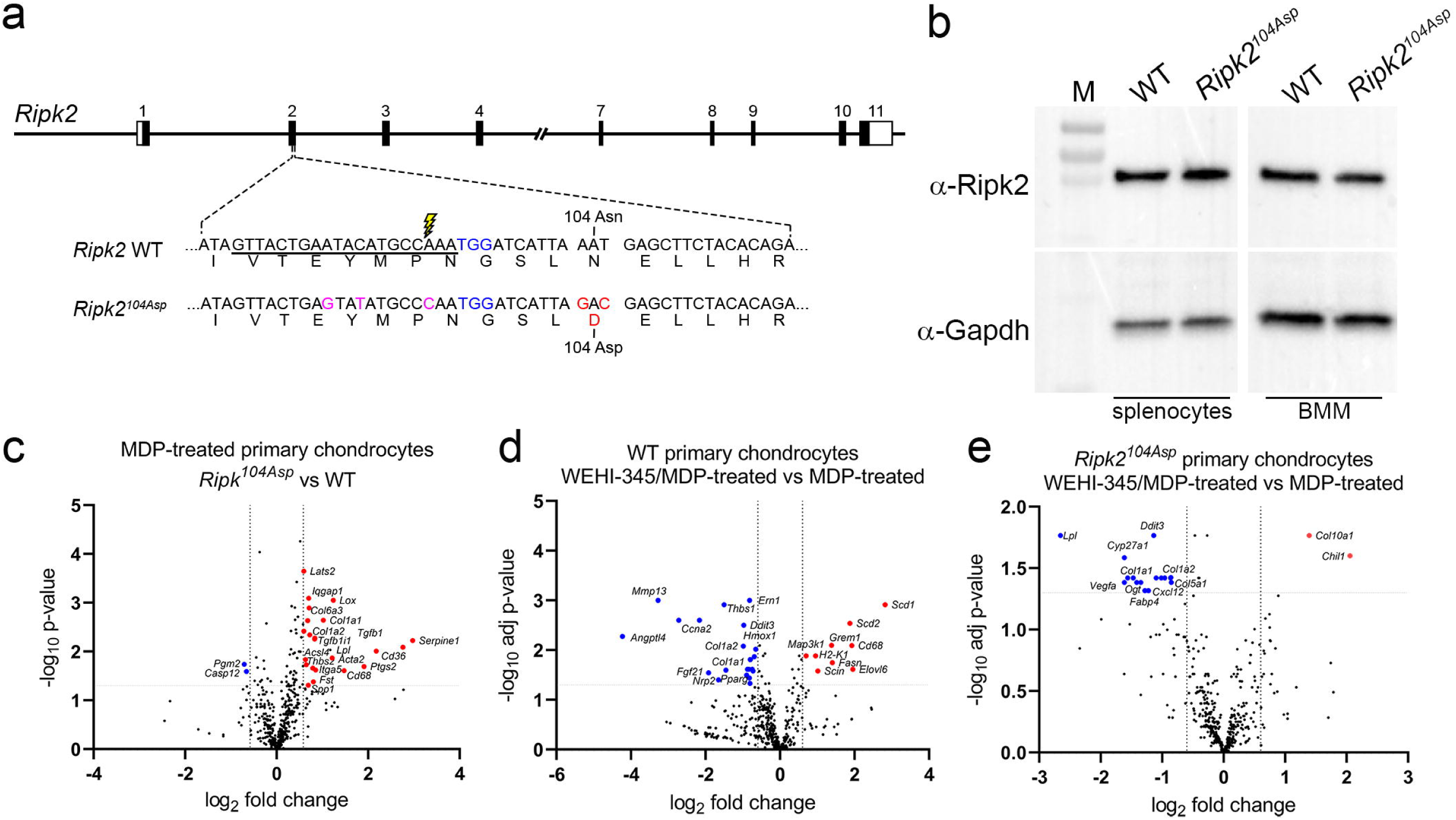
Generation and validation of the *Ripk2*^*104Asp*^ mouse. **a**, Schematic illustration of the mouse *Ripk2* locus and detailed view of exon 2. CRISPR/Cas9-stimulated homology directed repair was used to edit sequences of C57Bl/6J mice (WT) encoding the *Ripk2*^*104Asn*^ protein to generate an isogenic line that expressed the OA-associated *Ripk2*^*104Asp*^ protein from the native locus. The guide RNA target sequence is underlined, the PAM site is highlighted in blue, and the Cas9 cleavage site is denoted with lightning bolt. An oligonucleotide donor was used as a template to create the mutations to generate Asp 104 (red) as well as silent mutations (magenta) to prevent targeting of the modified locus. **b**, Immunoblot analysis indicates similar Ripk2 protein present in WT and *Ripk2*^*104Asp*^ splenocytes or bone marrow derived macrophages (BMM). Gapdh is used as a loading control. M = protein mass standards in kDa. **c**, A single copy of *Ripk2*^*104Asp*^ is sufficient to alter the gene expression response of primary articular chondrocytes to MDP treatment. Volcano plots indicate genes significantly upregulated (red) or downregulated (blue) in MDP-treated *Ripk2*^*104Asp*^ as compared to MDP-treated WT primary chondrocytes. **d, e**, Increased gene expression in response to MDP stimulation is dependent on Ripk2 activity. **d**, WT or **e**, *Ripk2*^*104Asp*^ primary chondrocytes were stimulated with MDP in the presence or absence of the Ripk2 inhibitor, WEHI-345. Volcano plots indicate genes significantly upregulated (red) or downregulated (blue) upon MDP-stimulation in the presence of the inhibitor.

To determine if the *Ripk2*^*104Asp*^ allele perturbed NOD/RIPK2 signaling, cultured primary articular chondrocytes^36^ were stimulated with the Nod2 agonist, muramyldipeptide (MDP). Both WT and *Ripk2*^*104Asp*^ chondrocytes responded to MDP by upregulating genes associated with proinflammatory signaling (Extended Data Fig. 4 and Supplementary Table 2). However, the transcriptional response of *Ripk2*^*104Asp*^ chondrocytes was significantly amplified as compared with that of WT controls, consistent with previous functional assays of the *Ripk2*^*104Asp*^ allele^23^ (Fig 1c). MDP-stimulated gene expression was indeed dependent on Ripk2 activity as co-incubation of chondrocytes with the Ripk2 inhibitor WEHI-345^37^ significantly reduced expression of many genes, including those whose expression is directly associated with OA (*Mmp13, Col1a1, Col1a2*, and *Ccna2*) (Fig. 1d and e).

The joints of mature *Ripk2*^*104Asp*^ animals appear structurally similar to those of WT mice with no histological evidence of joint degeneration (Fig. 2 a-d, i and j). Nevertheless, *Ripk2*^*104Asp*^ mice displayed increased sensitivity to experimentally induced OA, initiated by destabilization of the medial meniscus (DMM) of the stifle (knee) joint^38^. Eight weeks after DMM surgery *Ripk2*^*104Asp*^ mice exhibited a significant increase in the extent and severity of cartilage damage on the medial (both femoral condyle and tibial plateau) and lateral (femoral condyle) faces of the knee as compared to operated WT controls (Fig. 2a-j and Extended Data Fig. 5). Blinded histological scoring of the entire joint or individual quadrants revealed highly significant differences in the average and maximal OARSI scores^38^ of DMM-treated WT and *Ripk2*^*104Asp*^ mice (Fig. 2i and j). In contrast, no significant difference was evident in the degree of synovitis observed in the operated joints of the two groups of mice (Fig. 2k). As a non-invasive alternate method for inducing PTOA in 16-week-old mice, we employed mechanical rupture of the anterior cruciate ligament (ACL)^39^. Again, *Ripk2*^*104Asp*^ mice showed a significant increase in cartilage damage compared to WT controls when examined 5 weeks post-rupture (Extended Data Fig. 6).

**Figure 2.**
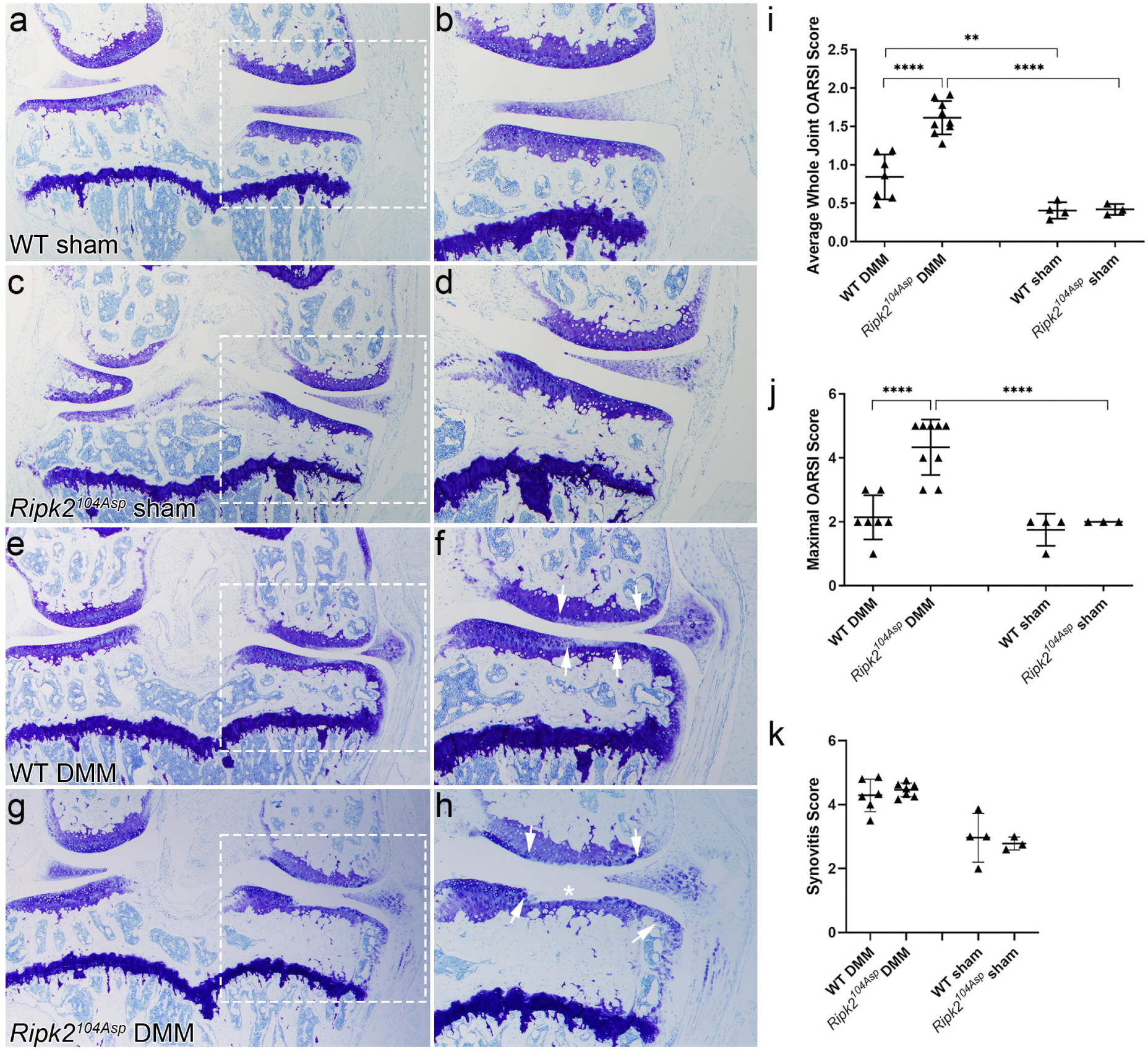
The *Ripk2*^*104Asp*^ allele acts dominantly and is sufficient to confer increased susceptibility to post-traumatic osteoarthritis. **a-d**, Knee joints of WT and *Ripk2*^*104Asp*^ mice that underwent sham surgery are similar histologically, with no indication of an OA phenotype. **e, f**, Following DMM surgery, WT knees displayed mild/moderate loss of proteoglycan content in the articular cartilage on the medial side of the knee (indicated by loss of toluidine blue staining). The extent of the damage is indicated by the arrows in **f. g, h**, Following DMM surgery, joints of *Ripk2*^*104Asp*^ mice displayed moderate/severe loss of proteoglycan content (arrows in **h**), cartilage fibrillation, and complete loss of articular cartilage in the medial tibial plateau (asterisk in **h**). **i**, Average whole joint and **j**, maximal OARSI scores of sham-operated and DMM-operated knee joints. **k**, There was no difference in the degree of synovitis between different genotypes. **a, c, e, g** are images of the entire knee joint; dashed boxes were magnified in **b, d, f, h** to focus on degradation on the medial side of the joint. Femur is up and medial is to the right in all images. WT sham (n=4), *Ripk2*^*104Asp*^ sham (n=3), WT DMM (n=7), *Ripk2*^*104Asp*^ DMM (n=9). All animals were analyzed 8 weeks post-surgery. Error bars represent ±SD and statistically significant differences of P ≤ 0.01 (**), P ≤ 0.001 (***), and P ≤ 0.0001 (****) were determined by two-way ANOVA with Tukey’s multiple comparisons test.

Injured knee joints of *Ripk2*^*104Asp*^ mice exhibited gene expression signatures associated with classical OA, but indicative of a more advanced disease state as compared with WT. Genes upregulated in the injured *Ripk2*^*104Asp*^ knee joints soon after ACL rupture are involved in both the innate and adaptive immune response (*Prg2, Trpc6, Bank1, Siglecf, Clca3a1, Isg15, Ighg1, Ighg2b*, and *Rsad2*) and include genes linked to OA (*B2m, A2m, Il17r, Ctsk, Angpt4, Aldh1a2*, and *Alox15*) (Fig. 3a and Supplementary Table 2). Conversely, many ECM genes whose depletion is associated with OA pathogenesis were significantly downregulated in *Ripk2*^*104Asp*^ joints, including *Acan, Col2a1, Col3a1, Comp, Prg4, Fn1, Hspg2, Matn2*, and *Cspg4* (Fig. 3a and Supplementary Table 2). Thus the transcriptional response of *Ripk2*^*104Asp*^ mice to injury closely parallels that of WT mice. However, it is exaggerated, with increased expression of inflammatory and catabolic factors and increased down-regulation of many ECM genes in the knee joints.

**Figure 3.**
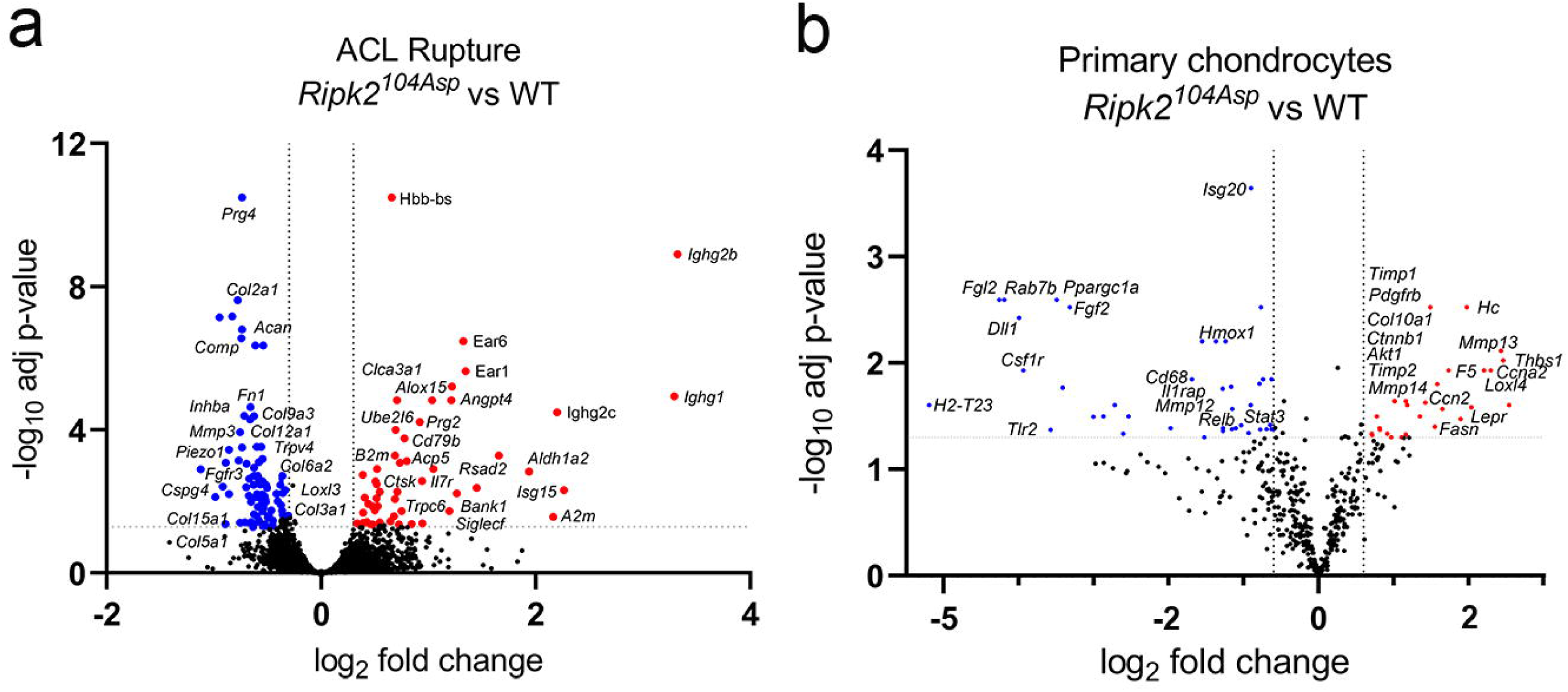
*Ripk2*^*104Asp*^ enhances expression of OA-associated markers in the surgically injured joint as well as primary articular chondrocytes. **a**, Comparative analysis of RNA-seq performed on whole joints isolated from WT or *Ripk2*^*104Asp*^ mice 10 days post ACL rupture. The volcano plot indicates genes significantly upregulated (red) or downregulated (blue) in *Ripk2*^*104Asp*^ as compared to WT joints. **b**, The nCounter Fibrosis panel was used to measure gene expression in AC cultured from WT or *Ripk2*^*104Asp*^ mice. Volcano plot indicates genes significantly upregulated (red) or downregulated (blue) in *Ripk2*^*104Asp*^ primary chondrocytes compared to WT primary chondrocytes.

In the absence of evident tissue remodeling that might indicate emergent OA in the knee joints of *Ripk2*^*104Asp*^ mice (Fig. 2a-d, i and j), we asked whether their joints exhibited altered signs of gene expression and/or inflammatory state. Analysis of primary chondrocytes revealed that gene expression was significantly altered in *Ripk2*^*104Asp*^ cells as compared with those isolated from WT mice (Fig 3b). Genes upregulated included well-known markers of OA, including hypertrophic chondrocytes and ECM remodeling (*Mmp13, Mmp14, Timp1, Timp2, Loxl4*, and *Col10a1*), growth factor signaling (*Ctnnb1, Ckap4*, and *Pdgfrb*), leptin signaling (*Lepr*), PI3K/Akt/mTOR signaling (*Akt1*), as well as genes involved in inflammatory signaling (*Lpcat1, Rbx1, Ccn2, Fasn, Cfhr2, F5, Elovl6*, Hc, and *Thbs1*) (Fig. 3b and Supplementary Table 2). The striking linkage between the altered gene expression profile of cultured *Ripk2*^*104Asp*^ chondrocytes and markers of mature OA led us to investigate if altered marker expression presaged the response to injury in the whole joint. The joints of both sham-operated and DMM mice revealed substantive effects of the *Ripk2*^*104Asp*^ allele on Nod/Ripk2 activity, matrix components, and markers of inflammation. Although Ripk2 is present at low levels in tibial chondrocytes of WT and *Ripk2*^*104Asp*^ animals subjected to sham surgery (Fig. 4a), pathway activity appears elevated in the *Ripk2*^*104Asp*^ joint, as reflected in higher levels of activated phospho-NF-κB (pNF-κB) compared with that of WT controls (Fig 4b). Following DMM surgery, pathway activity differences are enhanced, as expression of both Ripk2 and pNF-κB are considerably elevated in the operated joints of *Ripk2*^*104Asp*^ mice relative to WT (Fig. 4a and b). Similarly, expression of matrix markers is altered in sham-operated *Ripk2*^*104Asp*^ joints in a manner that appears to portend overt OA. As compared with WT controls, knee joints of sham surgery *Ripk2*^*104Asp*^ mice express elevated levels of Mmp13, a metalloproteinase that targets collagen for degradation (Fig. 4c). Mmp13 expression in the *Ripk2*^*104Asp*^ mice extends beyond the superficial layer of cartilage into deeper layers relative to WT controls (brackets in Fig. 4c). Consistent with this finding, collagen deposition scored by the presence of ColII appears relatively deficient in the joints of sham-operated *Ripk2*^*104Asp*^ mice (Fig. 4d). The differences in the levels of expression of Mmp13 and ColII between WT and *Ripk2*^*104Asp*^ mice become even more exaggerated following DMM surgery (Fig. 4c and d). These data indicate the *Ripk2*^*104Asp*^ allele alters the basal physiological status of the joint in ways that preview a definitive OA state.

**Figure 4.**
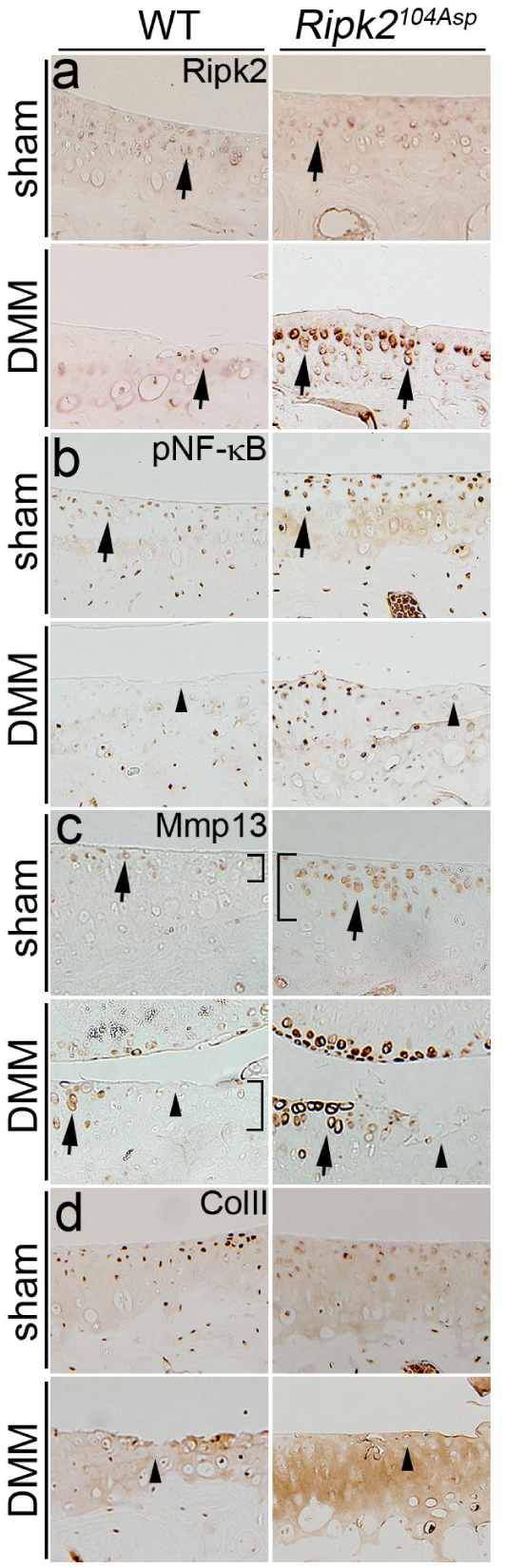
*Ripk2*^*104Asp*^ enhances NOD/RIPK2 signaling as well as OA-associated markers of matrix remodeling in uninjured joints and joints with PTOA. Immunohistochemical detection of **a**, Ripk2, **b**, pNF-κB, **c**, Mmp13, or **d**, ColII in WT and *Ripk2*^*104Asp*^ mice 8 weeks following sham or DMM surgery. **a**, Ripk2 is expressed at low levels in chondrocytes (arrows) of WT sham-, WT DMM-, and *Ripk2*^*104Asp*^ sham-operated joints. In contrast, Ripk2 expression is highly elevated in chondrocytes in *Ripk2*^*104Asp*^ knees following DMM surgery (arrows). **b**, Activated NF-κB (pNF-κB) expression levels are higher in *Ripk2*^*104Asp*^ sham-operated joints as compared to WT controls (arrows). The relatively increased Ripk2 expression is maintained in *Ripk2*^*104Asp*^ joints following DMM surgery. **c**, Mmp13 expression is elevated in chondrocytes of sham-operated *Ripk2*^*104Asp*^ mice as compared to WT controls (arrows); the relatively increased expression Ripk2 is maintained in *Ripk2*^*104Asp*^ mice following DMM surgery (arrows). Furthermore, in the unoperated *Ripk2*^*104Asp*^ joint, Mmp13 expression extends into deeper layers of cartilage, an expression domain normally only seen in WT joints following DMM surgery (brackets in **c**). **d**, ColII expression is reduced in *Ripk2*^*104Asp*^ sham surgery mice relative to WT controls, and this loss is further exacerbated by DMM surgery (arrows). In regions with severe cartilage damage (arrowheads), pNF-κB, Mmp13, and ColII expression is low in both WT and *Ripk2*^*104Asp*^ DMM-operated joints.

As *Ripk2*^*104Asp*^ chondrocytes have elevated expression of proinflammatory genes, and as local inflammation can sensitize joints to PTOA^40^, knees were examined in situ for the expression of inflammatory markers. iNos is a major inflammatory mediator of OA pathogenesis and is expressed in many tissues of the joint, including chondrocytes and macrophages^41^, both of which contribute directly to homeostatic maintenance of the joint^42,43^. Whereas control WT mice have uniformly low levels of iNos expression throughout the joint, *Ripk2*^*104Asp*^ mice exhibit a striking elevation of proinflammatory iNos signal in cartilage, meniscus, and synovial tissue (Fig. 5a). Following DMM surgery, iNos is induced in WT mice and its expression remains elevated in the cartilage, meniscus, and osteophytes of *Ripk2*^*104Asp*^ mice (Fig. 5a).

**Figure 5.**
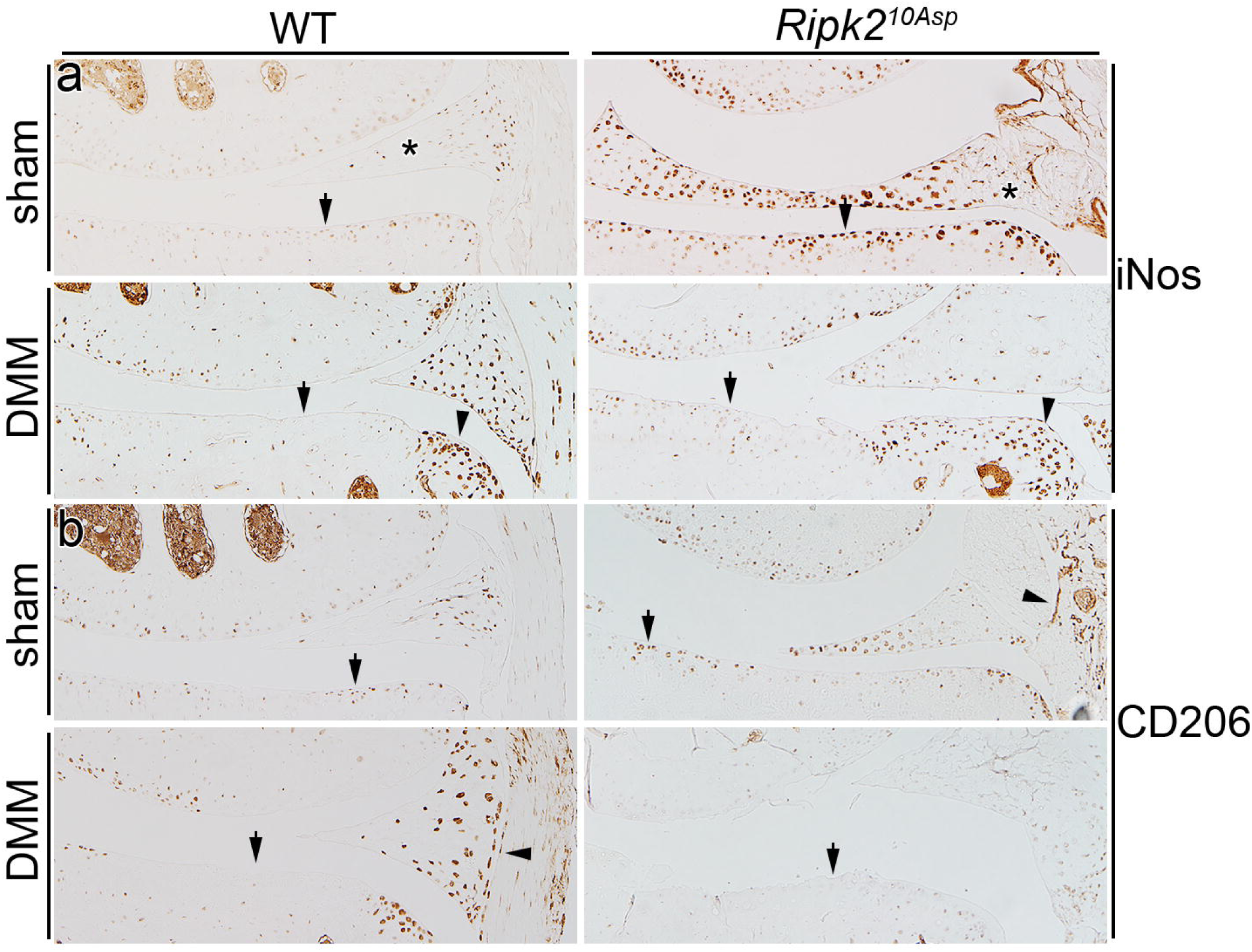
*Ripk2*^*104Asp*^ joints have elevated expression of proinflammatory markers. **a**, In WT mice, the proinflammatory marker iNos is normally expressed at low levels in the joint, and is markedly elevated following DMM surgery. In contrast, knee joints of *Ripk2*^*104Asp*^ have chronically high levels of iNos expression, independent of injury. In **a**, arrows indicate chondrocytes, asterisks mark the meniscus, and arrowheads indicate osteophytes in DMM-operated joints. **b**, There is no difference in expression of the anti-inflammatory marker, CD206, between sham-operated surgery WT and *Ripk2*^*104Asp*^ joints. Following DMM surgery, CD206 is prominently expressed in WT joints whereas it is almost absent in the joints of *Ripk2*^*104Asp*^ mice. In **b**, arrows indicate chondrocytes and arrowheads indicate synovium. All joints are 8 weeks post-surgery.

In the normal response to inflammation, CD206+ macrophages accrue in joints and are involved in resolving inflammation and tissue repair. Despite elevated iNos expression in the knees of sham-operated *Ripk2*^*104Asp*^ mice, there was no increase in the presence of CD206+ macrophages (Fig. 5b). Moreover, *Ripk2*^*104Asp*^ mice had a clear deficit in the recruitment of anti-inflammatory CD206+ cells into the cartilage, meniscus, and synovium of operated joints as compared with joints of WT mice (Fig. 5b). In sum, prior to overt injury, knee joints of *Ripk2*^*104Asp*^ mice exhibit higher than normal levels of inflammatory gene expression; following injury, *Ripk2*^*104Asp*^ mice are relatively poor at recruiting factors to resolve acute inflammation.

The NOD/RIPK2 signaling pathway operates broadly and the heightened expression of OA-associated markers in the knee joints of *Ripk2*^*104Asp*^ mice may reflect widespread elevation of inflammation. We examined the ability of cultured bone marrow-derived macrophages (BMM) from WT and *Ripk2*^*104Asp*^ mice to respond to MDP. In contrast, to the effects of the *Ripk2*^*104Asp*^ allele on the expression profile of chondrocytes, relatively few genes were differentially expressed upon stimulation of the BMM cells (Extended Data Fig. 7). To assess systemic differences in the inflammatory states of the mice, serum cytokine levels pre- or post-DMM surgery were measured. There was no difference in the serum concentration of any of 13 inflammatory cytokines sampled from 16-week old pre-surgery WT and *Ripk2*^*104Asp*^ mice (Fig. 6), indicating unoperated *Ripk2*^*104Asp*^ mice do not have a measurably elevated systemic phenotype. In contrast, in response to localized joint injury, *Ripk2*^*104Asp*^ mice mount a highly augmented systemic response. At 4 weeks following DMM surgery both WT and *Ripk2*^*104Asp*^ responded with raised serum levels of the proinflammatory cytokines IL1β, INFβ, and the anti-inflammatory IL10, but the degree of increase and the levels of these cytokines were significantly elevated in the *Ripk2*^*104Asp*^ mice (Fig. 6). The differences in serum cytokine response are transient, as they are resolved by 8 weeks post-surgery (Fig. 6).

**Figure 6.**
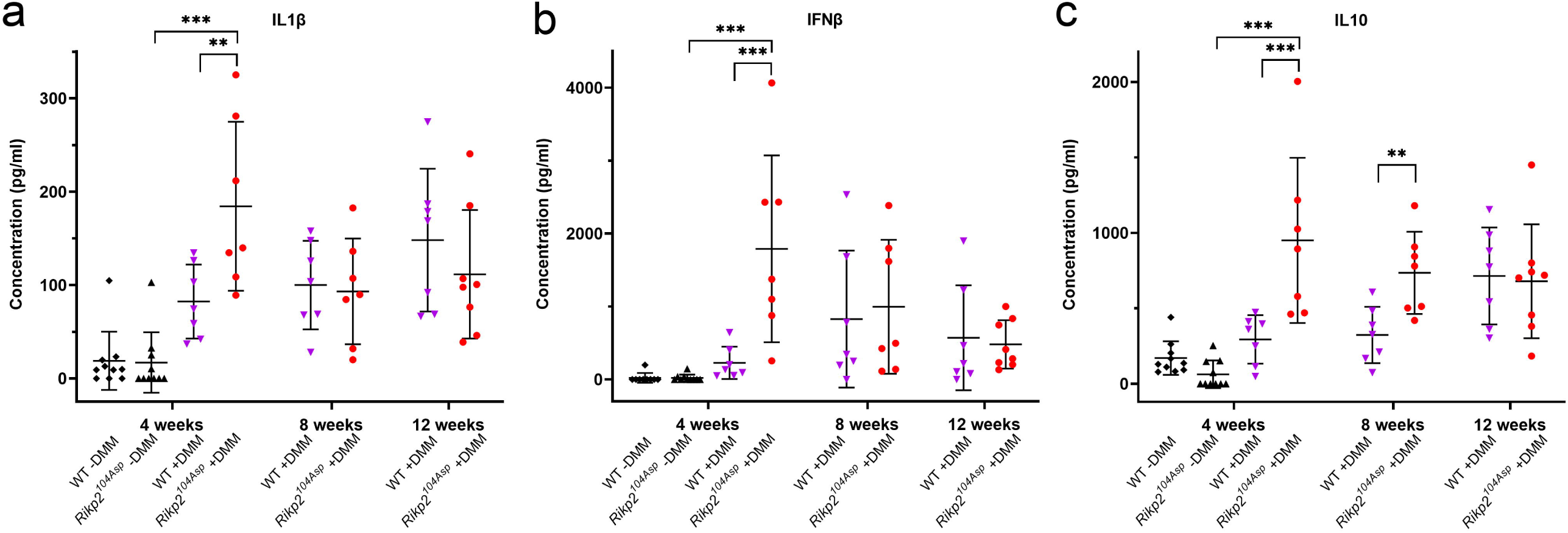
DMM surgery induces an acute systemic inflammatory response in *Ripk2*^*104Asp*^ mice. Quantification of serum **a**, IL1β, **b**, IFNβ, and **c**, IL10 levels from 16-week-old WT and *Ripk2*^*104Asp*^ mice just prior to DMM surgery (black diamonds and triangles) and at 4, 8 and 12 weeks post-surgery (magenta triangles and red circles). Error bars represent ±SD and statistically significant differences of P ≤ 0.01 (**) and P ≤ 0.001 (***) were determined by one-way ANOVA with Tukey’s multiple comparisons test (4-week post-DMM group) and a two-tailed unpaired t-test (8 and 12 week groups).

In sum, animals carrying the single amino acid change encoded by the *Ripk2*^*104Asp*^ variant have a magnified response to joint injury that leads to a predisposition to develop OA. The allele creates a chronically hyperactive inflammatory state in the joint with early signs of defective joint maintenance, as evidenced by gene expression in chondrocytes isolated from young mice and altered expression of pNF-κB, iNos, Mmp13, and ColII in mature animals. Nevertheless, the elevated activity of the NOD/RIPK2/NF-κB pathway caused by the variant allele has a very modest effect on tissue remodeling under normal laboratory conditions.

*Ripk2*^*104Asp*^ joints appear catabolically poised to progress to a severely damaged condition. Indeed *Ripk2*^*104Asp*^ mice mount an accelerated response to acute injury such as DMM surgery, characterized locally in the joint by amplified Ripk2 and pNF-κB expression, exaggerated changes in the expression of genes indicative of OA progression, including Mmp13 and ColII, and histologically recognizable deterioration. Both local as well as systemic inflammatory responses are amplified, seen by the deficit of CD206+ macrophages in the joint, altered gene expression in BMM, and the transient rise in serum cytokines.

We propose modulation of the NOD/RIPK2 signaling pathway is a general vulnerability factor for OA. We have found rare variants altering conserved positions in NOD/RIPK2 pathway proteins in 7 of the 151 (5%) families we have examined with non-syndromic OA. Supporting our hypothesis, hyperactivity of the NOD/RIPK2 pathway has been found associated with several human inflammatory syndromes that include arthritis is a comorbidity^16,17,44^. Our data indicate that modification of the NOD/RIPK2 pathway can render multiple joints (both weight and non-weight bearing) susceptible to OA. While the initiating factor (injury, repetitive use, obesity, etc.) may be different between joints and individuals, our work has shown that altered NOD/RIPK2 signaling is a predictive indicator of susceptibility to OA. Further pursuit of this signaling pathway and the spatiotemporal requirement for its activity may lead to assays for detection of early stages of disease and have therapeutic potential.

## Methods

### Study Approval

Written informed consent was obtained under the guidance of the Institutional Review Board at the University of Utah. Mice were maintained in accordance with approved institutional protocols at the University of Utah.

### Identification of families with a dominant pattern of OA inheritance

Our study utilizes data drawn from the Utah Population Database (UPDB) (https://uofuhealth.utah.edu/huntsman/utah-population-database/). The UPDB provides person-based interlinked records documenting genealogy, medical records, and vital statistics for over 11 million individuals from the late 18th century to the present. Medical records derive from the two largest healthcare providers in Utah (Intermountain Healthcare and University of Utah Health), Medicare claims, and the Utah and Idaho Cancer Registries. Vital records include statewide birth, death, and marriage certificates, as well as drivers’ licenses. UPDB data are available for approved research projects. Privacy of individuals whose data is available through UPDB is strictly protected through the Utah Resource for Genetic and Epidemiological Research (https://rge.utah.edu), established by executive order of the Governor of Utah. We identified individuals with OA between 1996 and 2021 in the UPDB using the following diagnosis and related procedure codes: 1^st^ MTP joint OA – ICD-9 735.2 or CPT 28289 and 28750; distal and proximal interphalangeal OA – ICD-9 715.14, ICD-10 M19.04x and CPT 26862, 26863, 26860, 26861, 26535, or 26536; and glenohumeral osteoarthritis OA – ICD-9 715.11, ICD-10 M19.011, M19.012, M19.019, Z96.611, Z96.612, or M19.0x and CPT 23472. Individuals with any of the following codes were excluded: ICD-9 714.0, 714.2, or 714.3 and ICD-10 M05.xxx, M06.xx, or M08.xxx. Manual chart review was performed on affected individuals to verify our coding strategy to identify OA cases. To determine if there was excess familial clustering of OA in each pedigree, we utilized the Familial Standardized Incidence Ratio (FSIR), with a threshold of ≥ 2.0. FSIR allows for the quantification of familial risk of a disease by comparing the incidence of a disease in a family to its expected incidence in the general population. See Kazmers, 2021 and Kazmers, 2020 for detailed methods ^45,46^. Pedigrees segregating a dominant pattern of OA inheritance were selected for genomic analysis.

### Whole exome sequencing and analysis

Whole exome sequencing (WES) and analysis was performed using genomic DNA isolated from whole blood or saliva as previously described^23^. Libraries were prepared using the Agilent SureSelect XT Human All Exon + UTR (v8) kit followed by Illumina NovaSeq 6000 150 cycle paired end sequencing. We followed best practices established by the Broad Institute GATK for variant discovery (https://gatk.broadinstitute.org/hc/en-us). Analysis of variants was performed with ANNOVAR (http://annovar.openbioinformatics.org/en/latest/)^47^ and pVAAST (http://www.hufflab.org/software/pvaast/)^48^ in concert with PHEVOR2 (http://weatherby.genetics.utah.edu/phevor2/index.html)^49^. pVAAST is a probabilistic search tool that classifies variants with respect to the likely effect they have on gene function. It incorporates information including position of a variant within a gene, cross-species phylogenetic conservation, biological function, and pedigree structure. PHEVOR2 works in concert with the output of pVAAST to integrate phenotype, gene function, and disease information for improved power to identify disease-causing alleles.

### Generation of the *Ripk2*^*104Asp*^ allele

CRISPR/Cas9-stimulated Homology Directed Repair (HDR) was used to edit^50,51^ the WT *Ripk2* (p.104Asn) gene of C57BL/6J mice to generate an isogenic line that expressed the variant Ripk2^104Asp^ protein from the native locus. The pair of alleles is analogous to the human WT *RIPK2*^*104Asn*^ and disease-associated *RIPK2*^*104Asp*^ alleles^23^. Briefly, fertilized C57BL/6J eggs were injected with Cas9 ribonucleoprotein complex targeting a site in exon 2 of Ripk2 within 10 base pairs of the Asn codon, and a DNA oligonucleotide to direct the single amino acid coding change^50,51^. *Ripk2*^*Asp104*^ mice were generated by the University of Utah Transgenic and Gene Targeting Mouse Core. The template for homologous recombination (synthesized by Integrated DNA Technologies) was a 200nt single-stranded oligodeoxynucleotide (ssODN) with ∼100bp homology arms on each side of the target site and containing several synonymous substitutions to prevent the Cas9 ribonucleoprotein complex from cutting the locus once it had been modified. An EcoRI restriction endonuclease site was also introduced to allow easy identification of the edited allele. The guide RNA sequence is GTTACTGAATACATGCCAAA(TGG), with the PAM site in parentheses. Genotyping primers are as follows – *Ripk2* gF1 5’-ATTTGCAATGAGCCTGAATTC-3’ and *Ripk2* gR1 5’-CTAAAGAGCCATTGGGCATATAC-3’.

### Destabilization of the medial meniscus surgery and anterior cruciate ligament rupture

DMM surgery, OARSI scoring of OA severity, and scoring of synovitis severity was performed as previously described^38,52^. Sham surgeries consisted of opening the joint capsule and exposing, but not transecting, the medial meniscotibial ligament. Non-invasive ACL rupture was induced by a single overload cycle of tibial compression as previously described^39^. Sixteen-week old male mice were used in all DMM and ACL rupture experiments.

### Histology and Immunohistochemistry

Animals were euthanized and knees were dissected and fixed in 10% neutral buffered formalin at 4°C for 48 hours, processed through an ethanol series, demineralized using FormiCal, equilibrated in xylenes, and embedded in paraffin. 5-6 µm thick tissue sections were cut and mounted onto slides. Tissue sections were deparaffinized and rehydrated as follows: Xylene 3X 5 minutes, 95% EtOH 2X 3 minutes, 70% EtOH 2X 3 minutes, ddH_2_O 1X 2 minutes, and 1X PBS for 5 minutes. For histological analysis, slides were stained with Toluidine blue. For immunohistochemistry, antigen retrieval was according to the antibody manufacturer’s protocol. Primary antibodies used were Mmp13 (Abcam, ab51072), ColII (Fitzgerald – 10R-C135B), Ripk2 (Abnova – H00008767-M02), pNF-κB (Thermofisher, 44-711G), CD206 (Abcam, ab64693), and iNos (Abcam, ab3523). Tissue sections were incubated with primary antibody at 4°C, for 18-24 hours in a humidified chamber, rinsed 3X with PBS, and then blocked with 2.5% horse serum (Vector Laboratories, MP-7401) for 20 minutes at 25°C. Block solution was replaced with ImmPress-HRP Horse anti-rabbit/mouse secondary antibody (Vector Laboratories, MP-7401/ MP-7402-15) and incubated for 30 minutes at room temperature followed by 2X 5 minute rinses with PBS. Slides were stained with DAB (Vector Laboratories, SK-4100) for 10-20 minutes, rinsed for 5 minutes with PBS, dried, and mounted.

### Cytokine measurements

Blood from puncture of the submandibular vein was collected into microfuge tubes and incubated at 4°C for a minimum of 2 hours. Serum was isolated by centrifuging whole blood at 1500g for 10 minutes. Cytokine concentrations were determined using the LEGENDplex Inflammation Panel (BioLegend) according to the manufacturer’s protocol.

### Primary cell cultures

Primary articular chondrocytes were cultured as previously described^36^. Briefly, mice were euthanized at postnatal day 21-28. The femur was dislocated from the acetabulum, and the articular cartilage cap was collected from the femoral head using blunt-ended forceps. Cartilage pieces were collected into 1X PBS in a 50 mL conical vial on ice, washed 3 times in 1X PBS, resuspended in collagenase D solution (5 mg/mL in DMEM), and incubated in tissue culture plates overnight. The next day, the suspensions were further dissociated by pipetting, filtered through a 70uM filter, and washed once in complete media. Cells were cultured at a density of 200,000 per mL in 24 well plates for 5-7 days, until adhered to the plate and approximately 80% confluent. Media was changed every 3 days. Bone marrow derived macrophages (BMM) were cultured as previously described^53^.

### MDP and WEHI-345 treatment

MDP was reconstituted in endotoxin-free water, as per manufacturer’s instructions (Invivogen). Stock solution was diluted into serum-free media, and cells were treated with a concentration of 10 µg/mL of MDP for 6 hours. WEHI-345 was reconstituted in DMSO as per manufacturer’s instructions (MedChemExpress). The working solution was prepared by diluting the stock in 1x PBS, and cells were treated at a final concentration of 1µM for 6 hours. As a control for MDP and WEHI-345 treatment, cells were treated with an equal volume of endotoxin-free water or 1µM DMSO for 6 hours, respectively.

### RNA isolation

Total RNA from primary articular chondrocytes or BMM was isolated using Direct-zol RNA Miniprep Kit (Zymo Research).

### Nanostring gene expression analysis

Primary articular chondrocyte or BMM RNA was used for targeted gene expression analysis using the nCounter Fibrosis Panel (Nanostring), which allows quantification of 770 genes, including many genes that have been associated with the OA phenotype. Samples were processed and analyzed using the Nanostring nCounter platform by the Molecular Diagnostics Core at the University of Utah. Three biological replicates were used for each condition. Data was analyzed by ROSALIND (https://rosalind.bio/), with a HyperScale architecture developed by ROSALIND, Inc. (San Diego, CA). Read Distribution percentages, violin plots, identity heatmaps, and sample MDS plots were generated as part of the QC step. Normalization, fold changes and p-values were calculated using criteria provided by Nanostring. ROSALIND follows the nCounter Advanced Analysis protocol of dividing counts within a lane by the geometric mean of the normalizer probes from the same lane. Housekeeping probes to be used for normalization are selected based on the geNorm algorithm as implemented in the NormqPCR R library^54^. Fold changes and pValues are calculated using the fast method as described in the nCounter Advanced Analysis 2.0 User Manual. P-value adjustment is performed using the Benjamini-Hochberg method of estimating false discovery rates (FDR).

### RNA-sequencing and gene expression analysis

ACL rupture was performed on 16-week-old male mice. Whole knee joints were dissected 10 days post-rupture in RNAlater (ThermoFisher) and stored at −80°C. Joints were cut into small fragments, transferred to a tube containing Trizol and 2.8mm stainless steel beads, and homogenized using the BeadBug microtube homogenizer. Total RNA was isolated as described above. Total RNA from 4 WT and 4 *Ripk2*^*104Asp*^ knee joints was used for RNA-sequencing (RNA-seq). RNA-seq, quality control, and alignment was performed by Novogene. Aligned reads were counted using HTSeq v0.11.3^55^, and the counts were then analyzed for differential gene expression using median-ratio-normalization^56^ with Deseq2 v1.30.0^57^. Fold-changes were calculated by comparing read counts in *Ripk2*^*104Asp*^ knee joints relative to WT knee joints. Genes with an adjusted P-value < 0.05 were considered differentially expressed.

### Quantitative PCR (qPCR)

DMM surgery was performed unilaterally on 16-week-old male mice. Whole knee joints were dissected from uninjured (control) or operated (DMM) knees for RNA isolation 10 days post-surgery as described above. 1µg total RNA was reverse transcribed into cDNA, and quantitative assessment of *Nod1, Nod2, Ripk2*, and β*-actin* sequences was performed as described previously^23^. The copy number of *Nod1, Nod2, Ripk2* was normalized to 1,000 copies of β-actin as previously described^58^. Primers used are as follows: β*-actin* F 5’-GTAACAATGCCATGTTCAAT-3’, β*-actin* R 5’-CTCCATCGTGGGCCGCTCTAG, *Nod1* F 5’-CCTGAAGCAGAACACCACACTG-3’, *Nod1* R 5’-CTTGGCTGTGATGCGATTCTGG-3’, *Nod2* F 5’-CCTAGCACTGATGCTGGAGAAG-3’, *Nod2* R 5’-CGGTAGGTGATGCCATTGTTGG-3’*Ripk2* F 5’-TCGTGTGGATCCTCTCTGCTCT-3’, *Ripk2* R 5’-TTCCAGGACAGTGGTGTGCCTT-3’.

### Statistical Analyses

Statistical analysis was performed using GraphPad Prism software. Tests performed and statistical significance are indicated in the figure legends. An unpaired t-test was used to determine statistical significance for qPCR analyses. P-values <0.05 were considered statistically significant.

## Acknowledgments

We would like to thank Charles L. Saltzman, Jerry Kaplan, and the members of the Department of Orthopaedics Osteoarthritis Discovery and Treatment Initiative advisory board for their support and critical feedback. We thank the families that participated in this study. This work was funded by the Skaggs Foundation for Research (MJJ and NHK), the Utah Genome Project (MJJ, NHK, DJG) the Arthritis National Research Foundation (MJJ – 707634), and the National Institute on Aging (MJJ and DJG – R21AG063534-01A1).

## Figure Legends

**Extended Data Figure 1.**
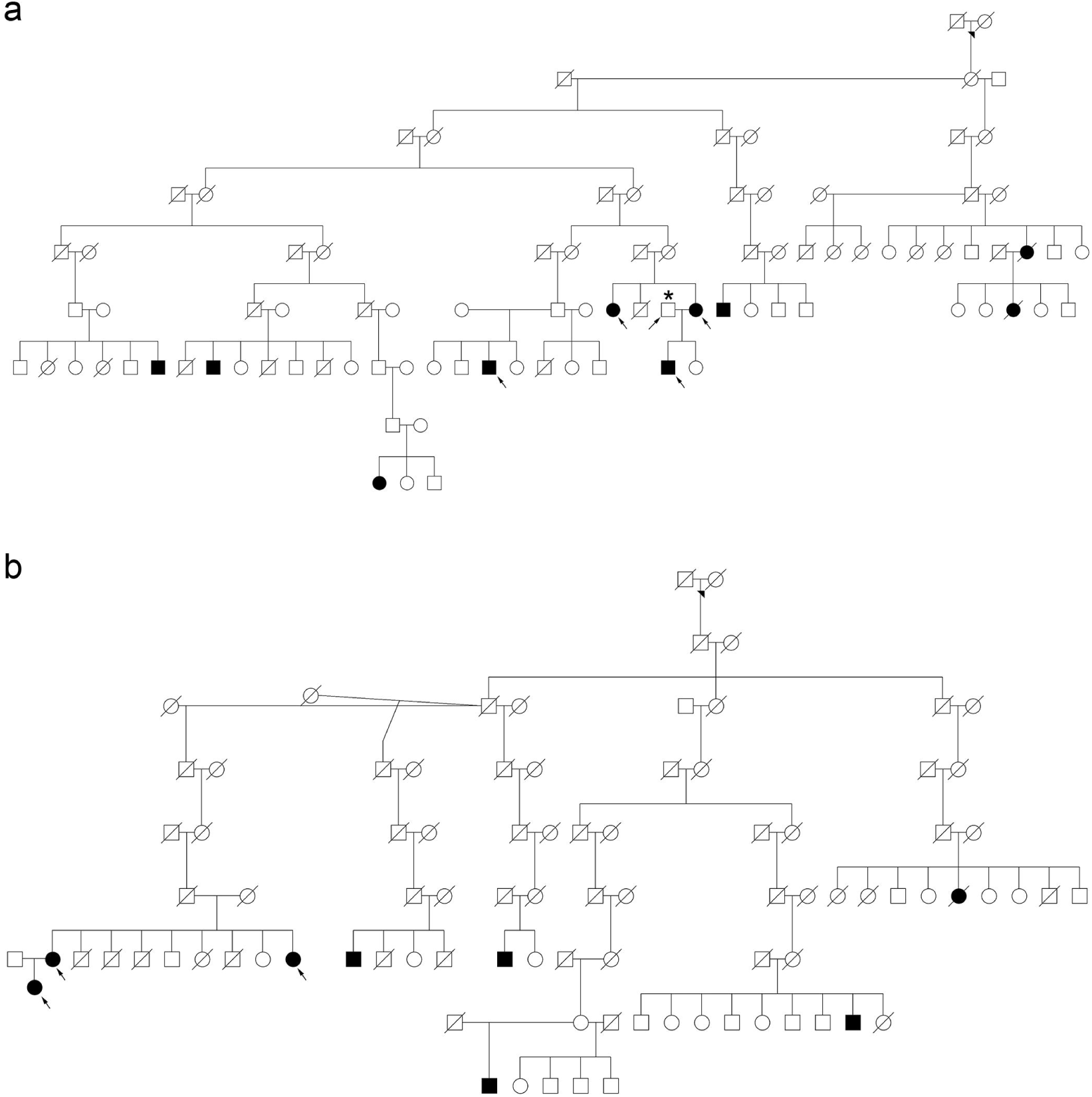
Examples of high-risk osteoarthritis pedigrees. Two multigeneration high-risk pedigrees segregating **a**, finger interphalangeal joint OA (FIJ744 family, *NOD1*) and **b**, glenohumeral OA (SA7355 family, *IKBKB*) identified from the Utah Population Database. The OA phenotype segregates as an apparent autosomal dominant trait. Circles = females, squares = males, slash = deceased. Filled circles/squares = affected individuals; open circles/squares = individuals with unknown affection status. Arrowhead indicates the founder of each family. Arrows indicate individuals who were sequenced and used for genomic analyses. Asterisk in **a** indicates an unaffected individual who was sequenced and used for genomic analyses.

**Extended Data Figure 2.**
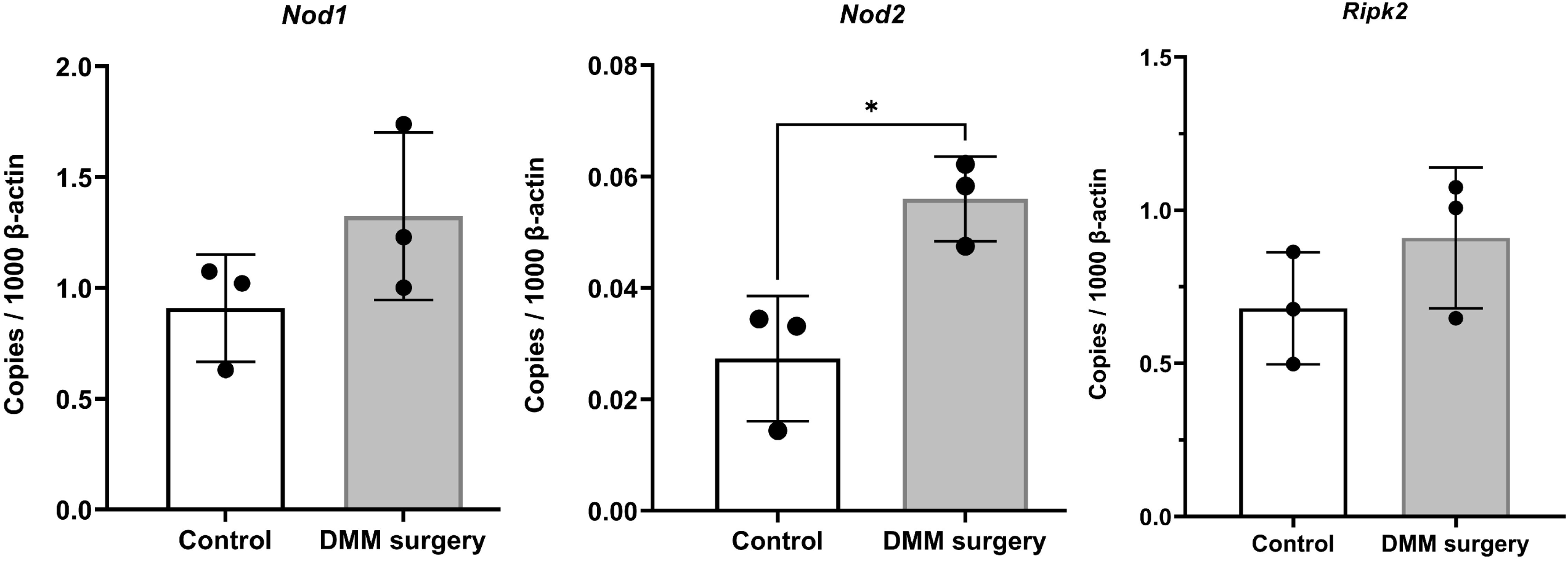
*Nod1, Nod2*, and *Ripk2* are expressed in the uninjured joint and the pathway is activated in response to joint injury. qPCR analysis of *Nod1, Nod2*, and *RIpk2* expression using RNA isolated from whole joints 10 days after DMM surgery. Error bars represent ±SD and a statistically significant difference of P ≤ 0.05 (*) was determined by a two-tailed unpaired t-test.

**Extended Data Figure 3.**
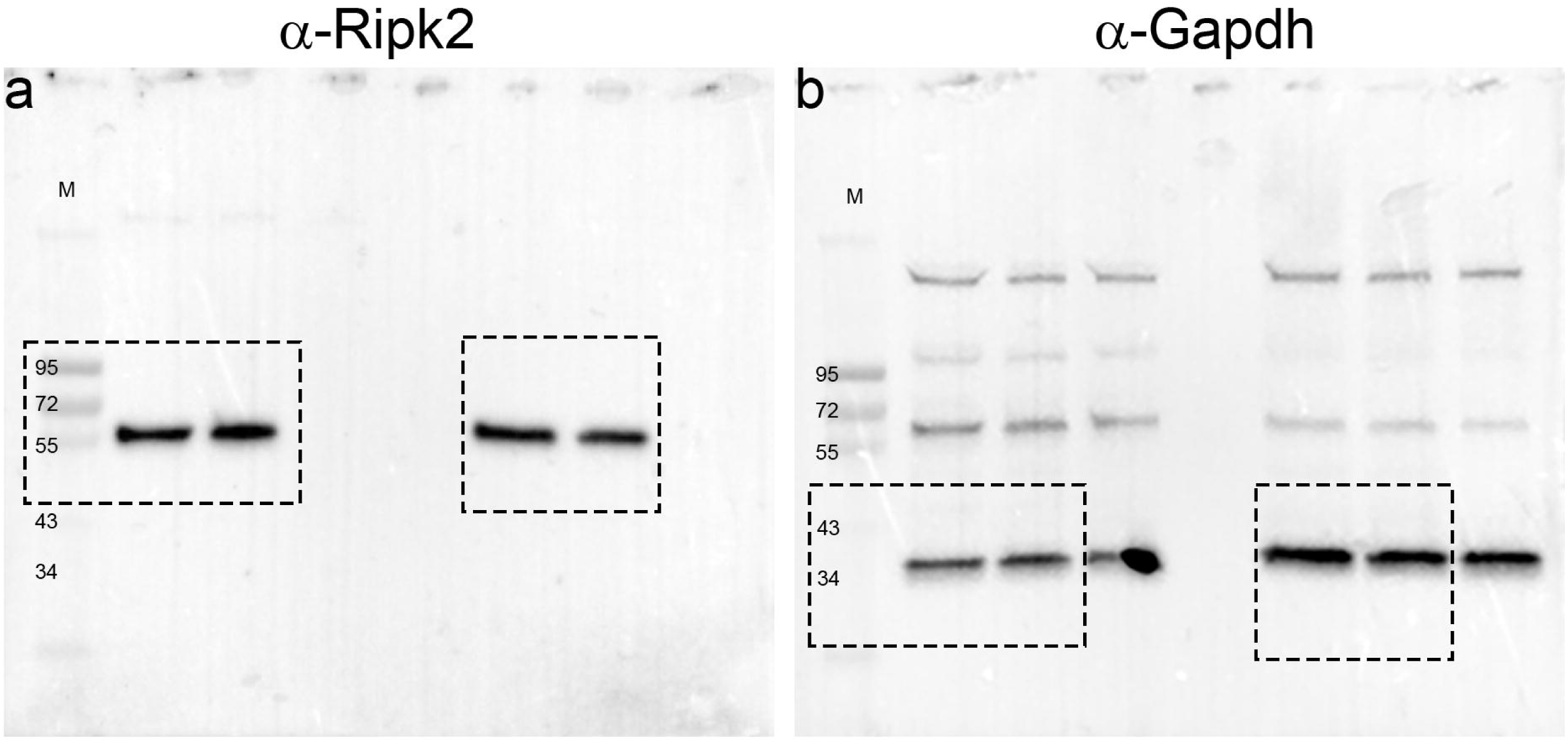
Unmodified Ripk2 and Gapdh immunoblots used to generate data in Fig. 1b. **a**, The D10B11 antibody (Cell Signaling Technology) was used to detect Ripk2. **b**, the sc-365062 antibody (Santa Cruz Biotechnology) was used to detect Gapdh. Dashed boxes indicate the portion of immunoblots that were used in Fig. 1b. M = protein mass standards in kDa.

**Extended Data Figure 4.**
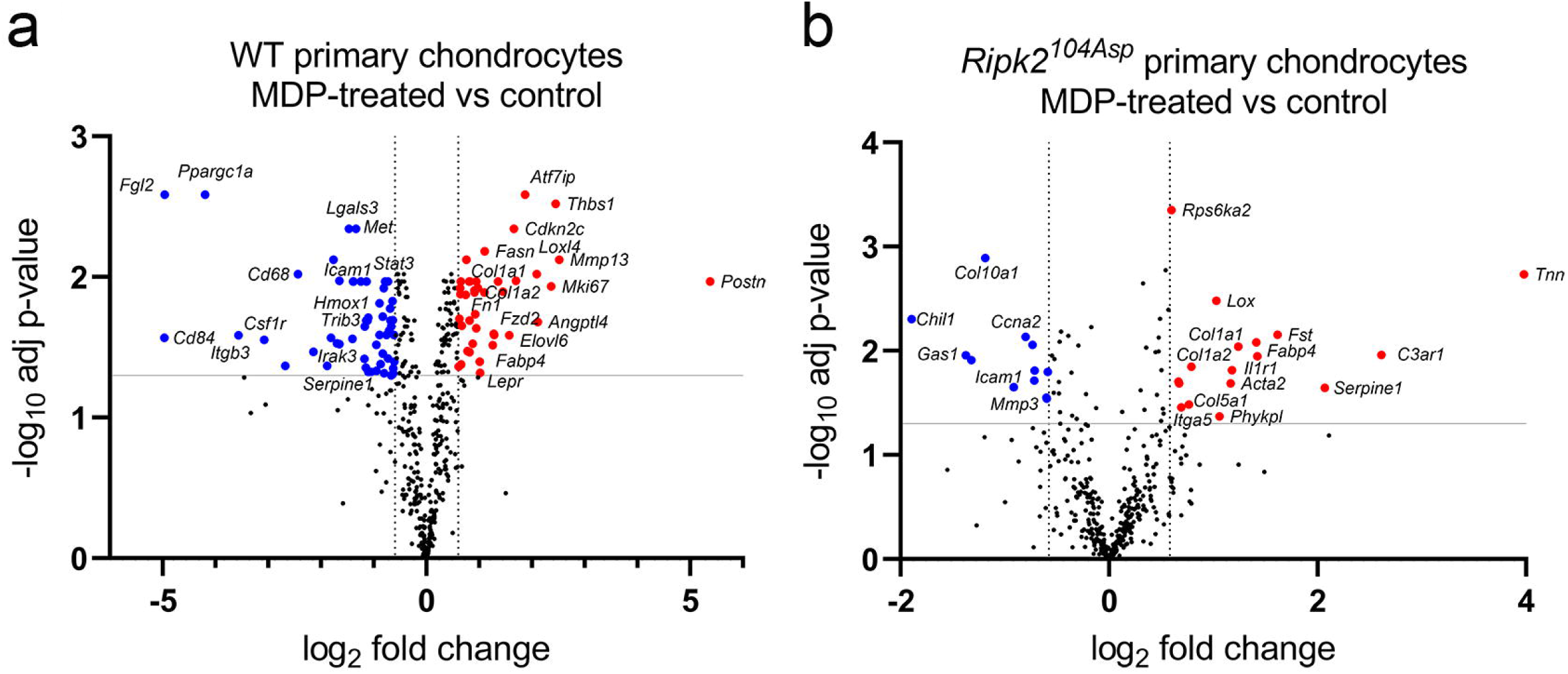
WT and *Ripk2*^*104Asp*^ primary chondrocytes respond to NOD2-stimulation by MDP. **a**, WT and **b**, *Ripk2*^*104Asp*^ primary chondrocytes were treated for 6 hours with 10 µg/mL MDP to activate the NOD2/RIPK2 pathway. **a**, Volcano plot illustrating genes significantly upregulated (red) or downregulated (blue) following stimulation of WT primary chondrocytes with MDP. **b**, Volcano plot illustrating genes significantly upregulated (red) or downregulated (blue) following stimulation of *Ripk2*^*104Asp*^ primary chondrocytes with MDP. The nCounter Fibrosis panel was used for all experiments.

**Extended Data Figure 5.**
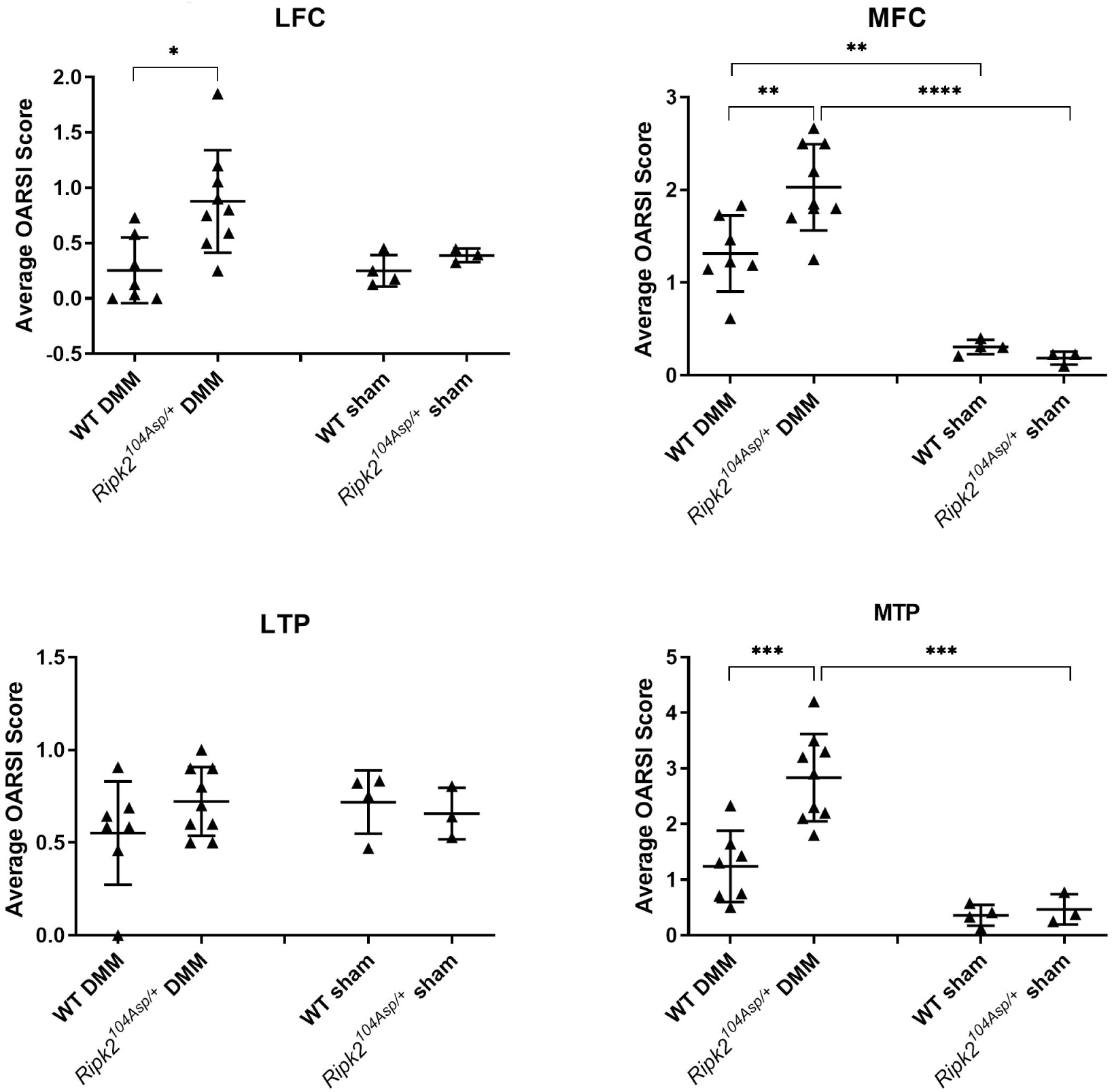
Regional analysis of the severity of OA in WT and *Ripk2*^*104Asp*^ joints subjected to sham or DMM surgery. Quantification of average OARSI scores of joints in 8-week post DMM WT sham (n=3), *Ripk2*^*104Asp*^ sham (n=4), WT DMM (n=7), and *Ripk2*^*104Asp*^ DMM (n=9) mice. LFC = lateral femoral condyle, MFC = medial femoral condyle, LTP = lateral tibial plateau, MTP = medial tibial plateau. Error bars represent ±SD and statistically significant differences of P ≤ 0.05 (*), P ≤ 0.01 (**), P ≤ 0.001 (***), and P ≤ 0.0001 (****) were determined by two-way ANOVA with Tukey’s multiple comparisons test.

**Extended Data Figure 6.**
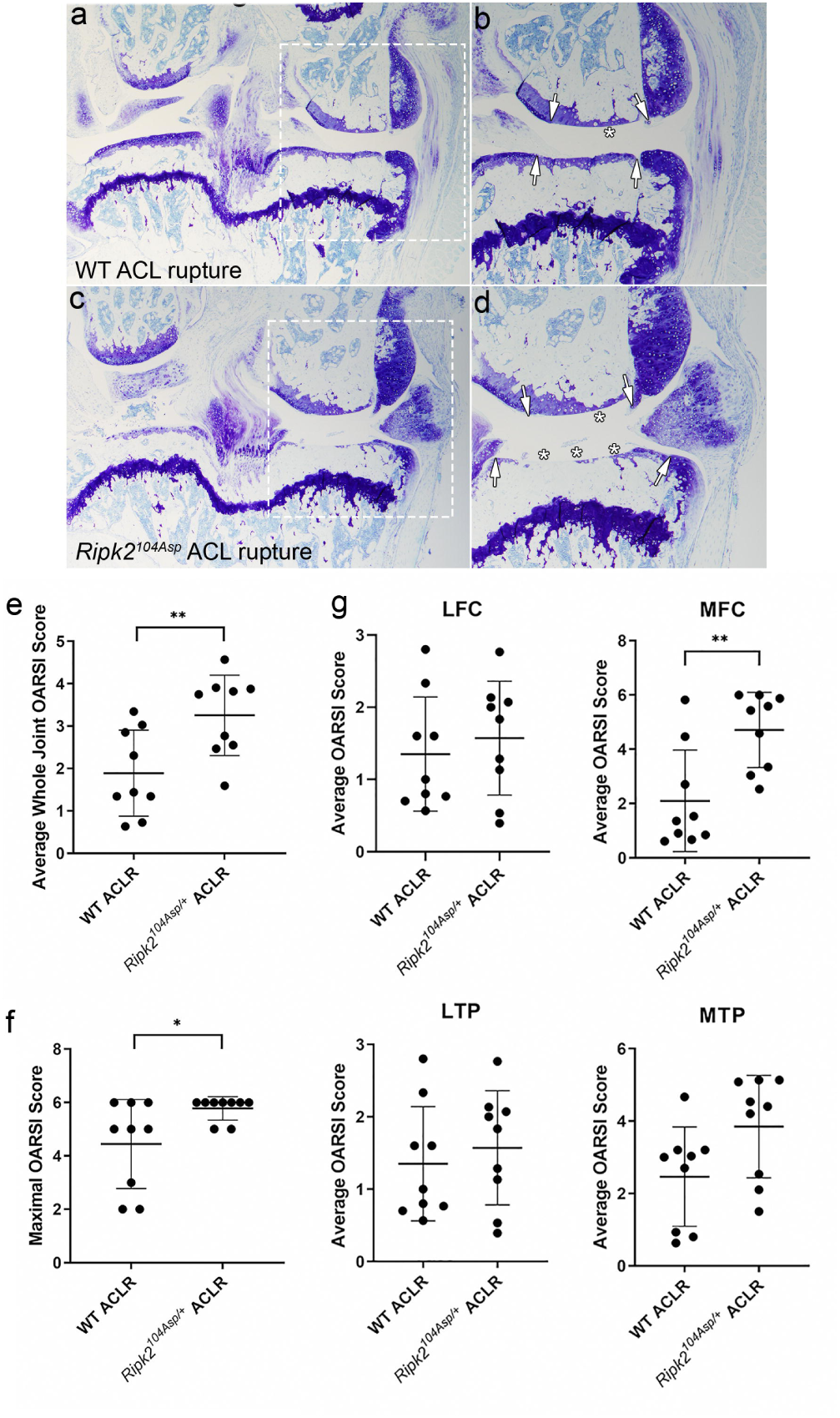
The *Ripk2*^*104Asp*^ allele acts dominantly and is sufficient to confer increased susceptibility to post-traumatic osteoarthritis induced by non-invasive ACL rupture. **a-d**, WT and *Ripk2*^*104Asp*^ mice underwent non-invasive ACL rupture (ACLR) at 16-weeks of age and were analyzed at 5 weeks post-rupture. **a, b**, Knee joints of WT mice subjected to ACL rupture displayed moderate/severe loss of proteoglycan content and cartilage (asterisk in **b**) on the medial side of the knee. The extent of damage is indicated by the arrows in **b. c, d**, Knee joints of *Ripk2*^*104Asp*^ mice subjected to ACL rupture displayed severe loss of proteoglycan content (arrows in **d**) and loss of cartilage that progressed down to the subchondral bone in the medial tibial plateau (asterisk in **d**). **e**,**f**, Knee joints of *Ripk2*^*104Asp*^ mice subjected to ACL rupture have an increased average whole joint and maximal OARSI scores compared to WT controls. **g**, regional analysis of the severity of OA in WT and *Ripk2*^*104Asp*^ joints subjected to ACL rupture. Tissue sections were stained with toluidine blue. **a** and **c** are images of the entire knee joint. Dashed boxes are presented at higher magnification in **b** and **d** to visualize the medial side of the joint. Femur is up and medial is to the right in all images. LFC = lateral femoral condyle, MFC = medial femoral condyle, LTP = lateral tibial plateau, MTP = medial tibial plateau. WT ACL rupture (n=9) and *Ripk2*^*104Asp*^ ACL rupture (n=9). Error bars represent ±SD and statistically significant differences of P ≤ 0.05 (*) and P ≤ 0.01 (**) were determined by a two-tailed unpaired t-test.

**Extended Data Figure 7.**
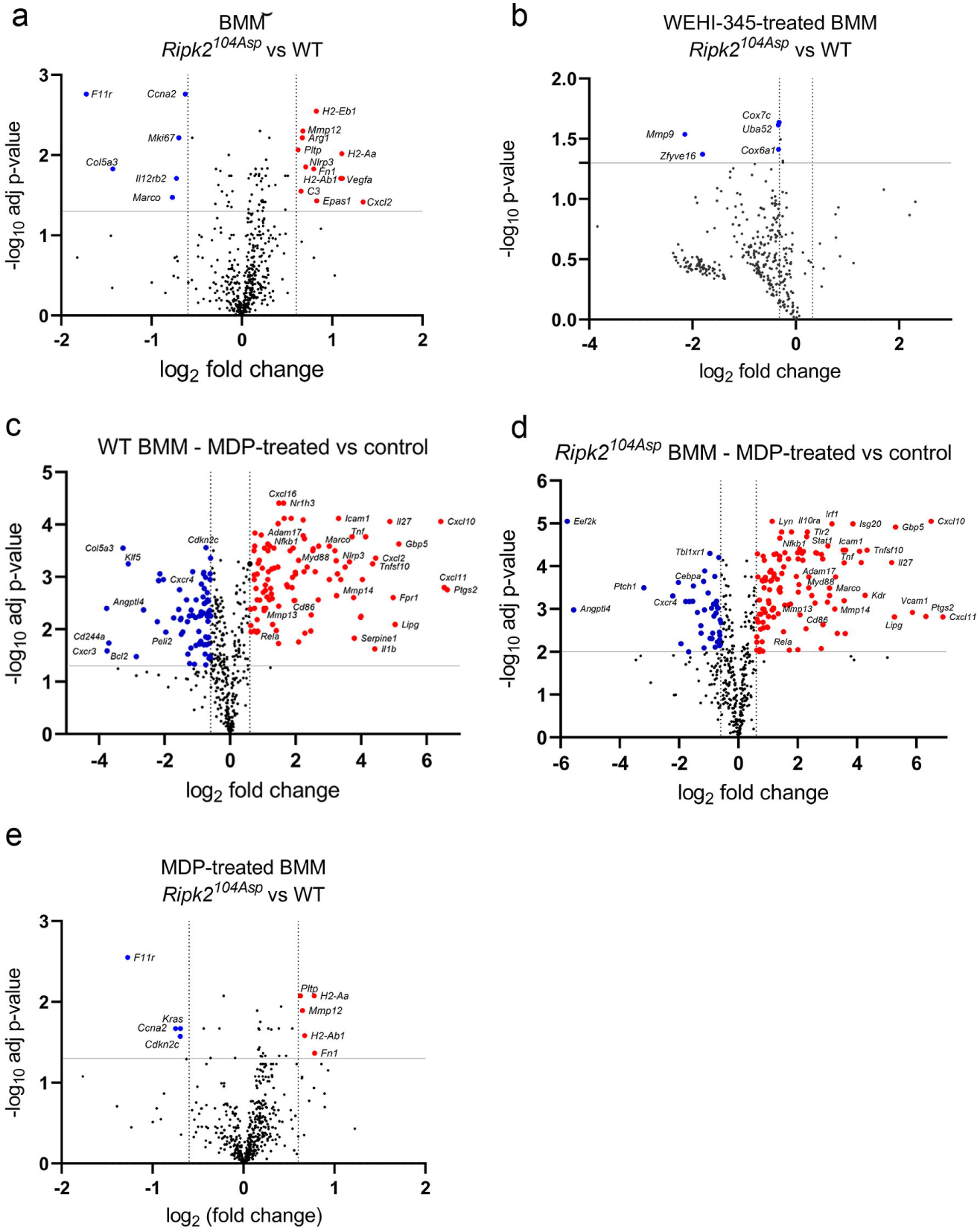
*Ripk2*^*104Asp*^ acts dominantly and is sufficient to alter gene expression in cultured BMM. Gene expression in BMM was measured using the nCounter Fibrosis panel. **a**, Volcano plot indicating genes with significantly altered expression in *Ripk2*^*104Asp*^ as compared WT cultured BMM. **b**, Volcano plot indicating that very few genes are differentially expressed between *Ripk2*^*104Asp*^ and WT cultured BMM following treatment with the RIPK2 inhibitor, WEHI-345. **c**, Volcano plot indicating genes with significantly altered expression following stimulation of WT BMM with MDP. **d**, Volcano plot indicating genes with significantly altered expression following stimulation of *Ripk2*^*104Asp*^ BMM with MDP. **e**, Volcano plot indicating genes with significantly altered expression comparing MDP-stimulated *Ripk2*^*104Asp*^ BMM with MDP-stimulated WT BMM.

## Table Legends

Supplemental Table 1. Family and Phenotype Details.

Supplemental Table 2. nCounter Fibrosis Panel and RNA-seq Differential Gene Expression Data.

