## Supplementary Table 2 for "The NOD/RIPK2 signaling pathway contributes to osteoarthritis susceptibility"

Primary articular chondrocytes - *Ripk2* $^{104Asp}$ vs WT
Nanostring Fibrosis Panel

**Downregulated**

| upperSymb | Symbol | GeneID | Alias | Descriptior | 20210507_ | 20210507_ | 20210507_ | 20210302_ | 20210302_ | 20210302_ | baseMean | log2FoldCh | pvalue | padj | FC |
|---|---|---|---|---|---|---|---|---|---|---|---|---|---|---|---|
| H2-T23 | H2-T23 | 15040 | 37b\|37c\|H | histocompa | 6.069182 | 6.170469 | 5.857816 | 4.355141 | 4.96217 | 4.241966 | 38.74999 | -5.19555 | 0.00195 | 0.02481 | 36.64504 |
| FGL2 | FGL2 | 14190 | AI385601\| | fibrinogen- | 4.571682 | 4.836568 | 4.463537 | 4.355141 | 3.96217 | 4.367497 | 21.49753 | -4.2595 | 1.42E-05 | 0.002539 | 19.15302 |
| RAB7B | RAB7B | 226421 | 5430435G | RAB7B_ m | 4.256809 | 4.711037 | 4.726571 | 4.470618 | 4.547133 | 4.482974 | 23.14332 | -4.19277 | 1.63E-05 | 0.002539 | 18.28736 |
| DLL1 | DLL1 | 13388 | Delta1 | delta-like 1 | 4.680617 | 4.42153 | 4.192235 | 3.940104 | 4.769525 | 3.782535 | 19.66772 | -3.99672 | 7.22E-05 | 0.003744 | 15.96367 |
| CSF1R | CSF1R | 12978 | AI323359\| | colony stin | 4.10871 | 4.806194 | 4.793686 | 4.577534 | 4.132095 | 2.782535 | 18.38077 | -3.93802 | 0.000459 | 0.011674 | 15.32721 |
| TLR2 | TLR2 | 24088 | Ly105 | toll-like rec | 4.680617 | 4.643923 | 4.463537 | 4.470618 | 3.547133 | 4.689425 | 21.34573 | -3.57394 | 0.005743 | 0.04257 | 11.90869 |
| PPARGC1A | PPARGC1A | 19017 | A830037N | peroxisom | 3.899257 | 4.251605 | 4.034694 | 3.940104 | 3.769525 | 3.782535 | 15.41525 | -3.49338 | 2.17E-05 | 0.002539 | 11.26188 |
| ADH1 | ADH1 | 11522 | ADH-AA\|A | alcohol del | 4.256809 | 4.05896 | 4.463537 | 4.470618 | 4.421602 | 4.58989 | 20.77682 | -3.41587 | 0.000949 | 0.017038 | 10.6728 |
| FGF2 | FGF2 | 14173 | Fgf-2\|Fgfb | fibroblast g | 3.706612 | 4.158496 | 3.97811 | 4.355141 | 4.66261 | 3.782535 | 17.23478 | -3.32257 | 5.01E-05 | 0.002986 | 10.00445 |
| ARRB1 | ARRB1 | 109689 | 1200006I1 | arrestin_ b | 3.543113 | 4.05896 | 4.334254 | 4.470618 | 4.284099 | 4.58989 | 18.55182 | -3.00496 | 0.003086 | 0.032028 | 8.027551 |
| ITGA9 | ITGA9 | 104099 | (alpha)9\|2 | integrin alp | 3.599697 | 4.205802 | 4.378648 | 4.577534 | 4.66261 | 4.58989 | 20.19179 | -2.87 | 0.002939 | 0.031675 | 7.310635 |
| FADD | FADD | 14082 | Mort1/FAD | Fas (TNFRS | 4.571682 | 4.836568 | 5.216897 | 4.770179 | 4.869061 | 4.689425 | 28.35705 | -2.7204 | 0.001833 | 0.02481 | 6.590558 |
| FAS | FAS | 14102 | AI196731\| | Fas (TNF re | 5.706612 | 4.979526 | 5.777197 | 5.018106 | 3.96217 | 4.95246 | 33.49821 | -2.60881 | 0.006674 | 0.046071 | 6.1 |
| H2-AB1 | H2-AB1 | 14961 | AI845868\| | histocompa | 7.079166 | 6.958964 | 6.81807 | 5.814573 | 5.354488 | 5.306097 | 74.64082 | -2.53614 | 0.002923 | 0.031675 | 5.800357 |
| PRKDC | PRKDC | 19090 | AI326420\| | protein kin | 5.39111 | 5.591455 | 5.356622 | 5.018106 | 5.049633 | 5.174852 | 38.41585 | -1.97262 | 0.00491 | 0.040947 | 3.924803 |
| CD68 | CD68 | 12514 | Lamp4\|Sca | CD68 antig | 7.959953 | 8.197049 | 7.665171 | 6.700916 | 6.485732 | 6.689425 | 155.7449 | -1.6896 | 0.000669 | 0.014192 | 3.225664 |
| LGALS3 | LGALS3 | 16854 | GBP\|L-34\| | lectin_ gala | 7.476686 | 7.290524 | 7.416403 | 6.355141 | 6.421602 | 6.140087 | 115.366 | -1.55303 | 0.000155 | 0.006248 | 2.93433 |
| FNIP2 | FNIP2 | 329679 | D630023B | folliculin in | 8.298001 | 8.223149 | 7.992465 | 6.836268 | 7.437904 | 6.306097 | 182.9933 | -1.52526 | 0.008108 | 0.049821 | 2.878378 |
| HMOX1 | HMOX1 | 15368 | D8Wsu38e | heme oxyg | 11.553 | 11.41588 | 11.27163 | 9.960003 | 10.12456 | 10.13126 | 1713.491 | -1.37067 | 0.000159 | 0.006248 | 2.585906 |
| IL1RAP | IL1RAP | 16180 | 6430709H | interleukin | 5.291574 | 5.251605 | 5.442778 | 5.293741 | 5.284099 | 5.174852 | 39.11838 | -1.28008 | 0.001011 | 0.017481 | 2.428532 |
| NR1H3 | NR1H3 | 22259 | AU018371 | nuclear rec | 5.585757 | 5.880962 | 5.963611 | 5.293741 | 5.769525 | 5.367497 | 49.98819 | -1.27809 | 0.005967 | 0.043543 | 2.425181 |
| MMP12 | MMP12 | 17381 | AV378681 | matrix met | 7.599697 | 8.42153 | 8.038158 | 7.127731 | 7.132095 | 6.826929 | 184.1014 | -1.27748 | 0.004852 | 0.040947 | 2.424149 |
| ERO1L | ERO1L | 50527 | ERO1-L | ERO1-like ( | 10.95216 | 11.13959 | 10.93231 | 9.878703 | 9.844814 | 9.683402 | 1356.02 | -1.24059 | 0.000161 | 0.006248 | 2.362954 |
| CDKN1A | CDKN1A | 12575 | CAP20\|CDI | cyclin-depe | 9.64745 | 9.909817 | 10.0225 | 8.873466 | 8.886983 | 8.530728 | 635.5336 | -1.16657 | 0.000894 | 0.016695 | 2.244767 |
| RELB | RELB | 19698 | shep | avian retic | 6.959953 | 6.735421 | 6.793686 | 6.414035 | 6.171624 | 5.95246 | 90.79429 | -1.1582 | 0.005259 | 0.0417 | 2.231792 |
| NCEH1 | NCEH1 | 320024 | Aadacl1\|B | neutral cho | 6.10871 | 6.273973 | 6.572659 | 5.899462 | 5.916367 | 5.689425 | 67.49768 | -1.15463 | 0.002319 | 0.02707 | 2.226265 |
| TRAF2 | TRAF2 | 22030 | AI325259 | TNF recept | 7.430638 | 7.182343 | 7.179744 | 6.724375 | 6.606027 | 6.274388 | 119.3939 | -1.1052 | 0.004651 | 0.040947 | 2.151285 |
| MET | MET | 17295 | AI838057\| | met proto- | 8.436475 | 8.25442 | 8.276768 | 7.442604 | 7.79506 | 7.086315 | 235.8851 | -1.03707 | 0.004118 | 0.038462 | 2.052061 |
| S100A4 | S100A4 | 20198 | 18A2\|42a\| | S100 calciu | 9.573449 | 10.21378 | 10.07403 | 9.083062 | 8.93945 | 9.262315 | 736.4012 | -0.93398 | 0.006328 | 0.045368 | 1.910533 |
| SPOPL | SPOPL | 76857 | 4921517N | speckle-typ | 7.147184 | 6.972704 | 7.095805 | 6.414035 | 6.74353 | 6.426391 | 111.426 | -0.90949 | 0.001906 | 0.02481 | 1.878379 |
| ISG20 | ISG20 | 57444 | 1600023I0 | interferon- | 8.603161 | 8.600333 | 8.577428 | 7.814573 | 7.869061 | 7.804903 | 296.4359 | -0.90184 | 4.86E-07 | 0.000227 | 1.868454 |
| BST2 | BST2 | 69550 | 2310015I1 | bone marr | 5.910484 | 5.821461 | 5.777197 | 5.770179 | 5.769525 | 5.689425 | 55.31934 | -0.7909 | 0.000771 | 0.015662 | 1.730151 |
| ATOX1 | ATOX1 | 11927 | AI256639\| | ATX1 (antio | 9.200753 | 9.21739 | 9.327192 | 8.544966 | 8.33721 | 8.765528 | 477.3289 | -0.78229 | 0.005499 | 0.042096 | 1.719862 |
| DDIT3 | DDIT3 | 13198 | CHOP-10\|( | DNA-dama | 12.00517 | 11.99728 | 12.00425 | 11.21416 | 11.19938 | 11.31781 | 3154.087 | -0.77072 | 3.60E-05 | 0.002986 | 1.706119 |
| COX4I2 | COX4I2 | 84682 | Cox4b\|Cox | cytochrom | 9.220058 | 9.071784 | 9.213837 | 8.525066 | 8.606027 | 8.397245 | 457.9362 | -0.74198 | 0.000648 | 0.014192 | 1.672473 |
| STAT3 | STAT3 | 20848 | 1110034C | signal tran | 9.23353 | 9.195585 | 9.043338 | 8.640543 | 8.66261 | 8.329429 | 461.7088 | -0.69381 | 0.005358 | 0.0417 | 1.617544 |
| FABP5 | FABP5 | 16592 | E-FABP\|Fa | fatty acid b | 10.27538 | 10.40642 | 10.58337 | 9.849663 | 9.690091 | 9.877932 | 1108.052 | -0.64908 | 0.003989 | 0.038019 | 1.56817 |
| BAX | BAX | 12028 | - | BCL2-assoc | 8.733728 | 8.786881 | 8.85389 | 8.30156 | 8.171624 | 8.306097 | 368.5279 | -0.63061 | 0.000649 | 0.014192 | 1.548214 |
| SEC61B | SEC61B | 66212 | 1190006C1 | Sec61 beta | 10.53316 | 10.46106 | 10.6885 | 10.0625 | 9.785538 | 10.0352 | 1227.063 | -0.62975 | 0.005319 | 0.0417 | 1.547298 |
| MAPK8 | MAPK8 | 26419 | AI849689\| | mitogen-ac | 7.446151 | 7.268414 | 7.328607 | 7.127731 | 6.984538 | 6.869998 | 144.098 | -0.60199 | 0.0052 | 0.0417 | 1.517808 |

**Upregulated**

| | | | | | | | | | | | | | | | |
|---|---|---|---|---|---|---|---|---|---|---|---|---|---|---|---|
| AKT1 | AKT1 | 11651 | Akt\|PKB\|P | thymoma v | 8.338049 | 8.604752 | 8.836091 | 9.347607 | 9.497446 | 9.179139 | 500.4842 | 0.70639 | 0.006739 | 0.046071 | 1.631716 |
| TIMP2 | TIMP2 | 21858 | D11Bwg11 | tissue inhib | 10.53225 | 10.89771 | 10.84946 | 11.38305 | 11.7287 | 11.31099 | 2221.051 | 0.709894 | 0.007354 | 0.047698 | 1.635684 |
| CTNNB1 | CTNNB1 | 12387 | Bfc\|Catnb\| | catenin (ca | 10.33649 | 10.49299 | 10.58692 | 11.25198 | 11.41955 | 11.08974 | 1862.396 | 0.773093 | 0.002984 | 0.031675 | 1.70893 |
| CKAP4 | CKAP4 | 216197 | 5630400A | cytoskeleto | 8.76808 | 9.008162 | 9.247143 | 9.833574 | 10.04166 | 9.659052 | 688.0061 | 0.810292 | 0.004889 | 0.040947 | 1.753567 |
| YWHAB | YWHAB | 54401 | 1300003C1 | tyrosine 3- | 7.043915 | 7.115791 | 7.229072 | 8.008583 | 8.284099 | 7.748319 | 190.2338 | 0.81598 | 0.005634 | 0.042436 | 1.760494 |
| AMOTL2 | AMOTL2 | 56332 | AW549739 | angiomotin | 6.350468 | 6.813848 | 6.826107 | 7.677069 | 7.820151 | 7.537422 | 144.0918 | 0.91711 | 0.006906 | 0.046071 | 1.888329 |
| MMP14 | MMP14 | 17387 | AI325305\| | matrix met | 6.899257 | 7.052505 | 7.141609 | 8.008583 | 8.421602 | 7.713272 | 186.0403 | 0.967188 | 0.00799 | 0.049821 | 1.955026 |
| TIMP1 | TIMP1 | 21857 | Clgi\|EPA\|T | tissue inhib | 10.10383 | 10.20289 | 10.52716 | 11.33337 | 11.24526 | 11.30903 | 1766.802 | 1.010493 | 0.001508 | 0.022878 | 2.0146 |
| VIM | VIM | 22352 | - | vimentin | 11.35818 | 11.69221 | 11.57594 | 12.56419 | 12.69984 | 12.40824 | 4239.761 | 1.012274 | 0.001433 | 0.022878 | 2.017088 |
| LPCAT1 | LPCAT1 | 210992 | 2900035H | lysophosph | 6.550309 | 6.52773 | 7.270892 | 7.93005 | 8.219558 | 7.815958 | 167.2369 | 1.103093 | 0.008007 | 0.049821 | 2.148148 |
| RBX1 | RBX1 | 56438 | 1500002P1 | ring-box 1 | 5.274296 | 5.790764 | 5.992465 | 6.940104 | 6.984538 | 7.030462 | 80.75269 | 1.160168 | 0.007139 | 0.046955 | 2.234835 |
| PDGFRB | PDGFRB | 18596 | AI528809\| | platelet de | 5.291574 | 5.44143 | 5.378648 | 6.603069 | 6.96217 | 6.537422 | 65.60432 | 1.16152 | 0.001519 | 0.022878 | 2.23693 |
| COL10A1 | COL10A1 | 12813 | Col10\|Col1 | collagen_ t | 10.14422 | 10.05896 | 10.2169 | 11.25601 | 11.6574 | 10.99928 | 1689.203 | 1.181616 | 0.001966 | 0.02481 | 2.268307 |
| FLNB | FLNB | 286940 | AL024016\| | filamin_ be | 7.256809 | 7.451278 | 7.651667 | 8.984498 | 9.075913 | 8.382448 | 280.8719 | 1.351434 | 0.002922 | 0.031675 | 2.551657 |
| CCN2 | CCN2 | 14219 | Ccn2\|Fisp1 | connective | 11.89502 | 12.22512 | 12.4946 | 13.56256 | 13.91509 | 13.40984 | 7734.221 | 1.418313 | 0.001615 | 0.023564 | 2.672729 |
| CDKN2C | CDKN2C | 12580 | C77269\|IN | cyclin-depe | 6.203038 | 6.349637 | 6.204619 | 7.735963 | 7.916367 | 7.725049 | 130.007 | 1.484313 | 5.11E-05 | 0.002986 | 2.797838 |
| FASN | FASN | 14104 | A630082H | fatty acid s | 5.453846 | 5.339068 | 5.601041 | 7.110029 | 7.437904 | 6.615425 | 76.61484 | 1.54716 | 0.004332 | 0.039668 | 2.922414 |
| ATF7IP | ATF7IP | 54343 | 2610204M | activating t | 5.469113 | 5.295999 | 5.442778 | 6.998997 | 7.354488 | 6.848624 | 75.32203 | 1.582872 | 0.000812 | 0.015802 | 2.995657 |
| CFHR2 | CFHR2 | 545366 | Cfhrb_4/2 | compleme | 5.39111 | 5.880962 | 5.826107 | 7.213122 | 7.782349 | 7.174852 | 93.36115 | 1.648351 | 0.002294 | 0.02707 | 3.134752 |
| F5 | F5 | 14067 | AI173222\| | coagulatio | 6.156645 | 6.228885 | 6.524079 | 8.046303 | 8.284099 | 7.869998 | 145.5127 | 1.732087 | 0.000475 | 0.011674 | 3.322081 |
| ELOVL6 | ELOVL6 | 170439 | C77826\|FA | ELOVL fam | 3.291574 | 3.836568 | 3.919216 | 5.577534 | 5.916367 | 5.537422 | 25.63033 | 1.893904 | 0.003372 | 0.033508 | 3.716394 |
| HC | HC | 15139 | C5\|C5a\|He | hemolytic c | 6.291574 | 6.555386 | 6.494125 | 8.538363 | 8.405114 | 8.344777 | 173.4316 | 1.971469 | 4.17E-05 | 0.002986 | 3.921671 |
| LEPR | LEPR | 16847 | LEPROT\|Le | leptin rece | 3.98672 | 5.109586 | 4.888843 | 6.770179 | 7.006565 | 6.510455 | 52.42045 | 2.033947 | 0.002109 | 0.025915 | 4.095238 |
| LOXL4 | LOXL4 | 67573 | 4833426I2 | lysyl oxidas | 9.654979 | 10.18308 | 10.34128 | 12.16303 | 12.58147 | 12.08803 | 2301.958 | 2.206259 | 0.000446 | 0.011674 | 4.614769 |
| CCNA2 | CCNA2 | 12428 | AA408589 | cyclin A2 | 4.7819 | 5.295999 | 5.28845 | 7.414035 | 7.547133 | 7.208799 | 76.42924 | 2.291795 | 0.000403 | 0.011674 | 4.896649 |
| MMP13 | MMP13 | 17386 | Clg\|MMP-1 | matrix met | 8.93268 | 9.349637 | 9.639176 | 11.74028 | 11.96709 | 11.55815 | 1479.782 | 2.431597 | 0.000214 | 0.007674 | 5.394901 |
| THBS1 | THBS1 | 21825 | TSP-1\|TSP | thrombosp | 8.200753 | 8.438957 | 8.504179 | 10.71416 | 11.2348 | 10.47602 | 773.261 | 2.46036 | 0.000284 | 0.009464 | 5.503539 |
| MKI67 | MKI67 | 17345 | D630048A | antigen ide | 4.325522 | 5.317695 | 5.216897 | 7.22961 | 7.927955 | 7.208799 | 73.74191 | 2.539733 | 0.001918 | 0.02481 | 5.814815 |

Bone Marrow Derived Macrophages- *Ripk2* $^{104Asp}$ vs WT

Nanostring Fibrosis Panel

<table>
<tr><td></td><td><u>Downregulated</u></td><td></td><td></td><td></td><td></td><td></td><td></td><td></td><td></td><td></td><td></td><td></td><td></td><td></td><td></td></tr>
<tr><td>upperSymb</td><td>Symbol</td><td>GeneID</td><td>Alias</td><td>Descriptior</td><td>20210614_</td><td>20210614_</td><td>20210614_</td><td>20210614_</td><td>20210614_</td><td>20210614_</td><td>baseMean</td><td>log2FoldCh</td><td>pvalue</td><td>padj</td><td>FC</td></tr>
<tr><td>F11R</td><td>F11R</td><td>16456</td><td>9130004G</td><td>F11 recept</td><td>7.190494</td><td>7.121934</td><td>7.064945</td><td>5.585564</td><td>5.604461</td><td>5.531684</td><td>81.56323</td><td>-1.72233</td><td>4.89E-06</td><td>0.001735</td><td>3.299684</td></tr>
<tr><td>COL5A3</td><td>COL5A3</td><td>53867</td><td>-</td><td>collagen_ t</td><td>4.495349</td><td>4.38858</td><td>4.238094</td><td>3.820029</td><td>3.73939</td><td>3.309292</td><td>15.98288</td><td>-1.42973</td><td>0.000559</td><td>0.01482</td><td>2.693969</td></tr>
<tr><td>MARCO</td><td>MARCO</td><td>17167</td><td>AI323439|</td><td>macrophag</td><td>6.009922</td><td>6.111046</td><td>6.28872</td><td>5.498101</td><td>5.632475</td><td>5.26865</td><td>55.77264</td><td>-0.77111</td><td>0.003299</td><td>0.033594</td><td>1.706587</td></tr>
<tr><td>IL12RB2</td><td>IL12RB2</td><td>16162</td><td>A930027I1</td><td>interleukin</td><td>7.021895</td><td>6.973542</td><td>6.799973</td><td>6.339403</td><td>6.179963</td><td>6.26865</td><td>96.82029</td><td>-0.72648</td><td>0.001357</td><td>0.019422</td><td>1.654602</td></tr>
<tr><td>MKI67</td><td>MKI67</td><td>17345</td><td>D630048A</td><td>antigen ide</td><td>10.73148</td><td>10.8521</td><td>10.83375</td><td>10.14915</td><td>10.11691</td><td>10.06269</td><td>1406.293</td><td>-0.69991</td><td>9.29E-05</td><td>0.006073</td><td>1.624405</td></tr>
<tr><td>CCNA2</td><td>CCNA2</td><td>12428</td><td>AA408589</td><td>cyclin A2</td><td>10.02784</td><td>10.01063</td><td>9.966015</td><td>9.384814</td><td>9.39556</td><td>9.356112</td><td>826.0943</td><td>-0.62837</td><td>6.95E-06</td><td>0.001735</td><td>1.545821</td></tr>
<tr><td></td><td></td><td></td><td></td><td></td><td></td><td></td><td></td><td></td><td></td><td></td><td></td><td></td><td></td><td></td><td></td></tr>
<tr><td></td><td><u>Upregulated</u></td><td></td><td></td><td></td><td></td><td></td><td></td><td></td><td></td><td></td><td></td><td></td><td></td><td></td><td></td></tr>
<tr><td>PLTP</td><td>PLTP</td><td>18830</td><td>Bpife|OD1</td><td>phospholip</td><td>11.57724</td><td>11.6174</td><td>11.69323</td><td>12.21949</td><td>12.22032</td><td>12.31086</td><td>3928.48</td><td>0.62217</td><td>0.000172</td><td>0.008577</td><td>1.539188</td></tr>
<tr><td>C3</td><td>C3</td><td>12266</td><td>AI255234|</td><td>compleme</td><td>9.334253</td><td>9.177787</td><td>9.484213</td><td>9.965707</td><td>9.932035</td><td>10.04475</td><td>807.0185</td><td>0.655222</td><td>0.002494</td><td>0.028113</td><td>1.574858</td></tr>
<tr><td>ARG1</td><td>ARG1</td><td>11846</td><td>AI|AI2565</td><td>arginase_ l</td><td>6.292856</td><td>6.441691</td><td>6.400366</td><td>7.011171</td><td>6.998124</td><td>6.972257</td><td>102.9698</td><td>0.666719</td><td>0.00011</td><td>0.006073</td><td>1.587458</td></tr>
<tr><td>MMP12</td><td>MMP12</td><td>17381</td><td>AV378681</td><td>matrix met</td><td>13.31856</td><td>13.25277</td><td>13.22678</td><td>13.89292</td><td>13.96486</td><td>13.96522</td><td>12447.06</td><td>0.67539</td><td>5.01E-05</td><td>0.005004</td><td>1.597029</td></tr>
<tr><td>NLRP3</td><td>NLRP3</td><td>216799</td><td>AGTAVPRL</td><td>NLR family</td><td>6.478071</td><td>6.441691</td><td>6.518202</td><td>7.179925</td><td>7.04056</td><td>7.183761</td><td>111.9752</td><td>0.704866</td><td>0.000392</td><td>0.013982</td><td>1.629994</td></tr>
<tr><td>FN1</td><td>FN1</td><td>14268</td><td>E330027I0</td><td>fibronectin</td><td>7.103032</td><td>7.236576</td><td>7.005648</td><td>7.941045</td><td>7.8024</td><td>7.880834</td><td>180.3833</td><td>0.794463</td><td>0.000572</td><td>0.01482</td><td>1.734431</td></tr>
<tr><td>H2-EB1</td><td>H2-EB1</td><td>14969</td><td>Eb|H-2Eb|</td><td>histocomp</td><td>6.529296</td><td>6.509595</td><td>6.489633</td><td>7.288178</td><td>7.341426</td><td>7.205456</td><td>118.9269</td><td>0.824296</td><td>1.70E-05</td><td>0.002821</td><td>1.770671</td></tr>
<tr><td>EPAS1</td><td>EPAS1</td><td>13819</td><td>HIF-2alpha</td><td>endothelia</td><td>6.64219</td><td>6.767091</td><td>6.587244</td><td>7.261867</td><td>7.43983</td><td>7.639212</td><td>133.0882</td><td>0.828336</td><td>0.003779</td><td>0.03698</td><td>1.775636</td></tr>
<tr><td>H2-AB1</td><td>H2-AB1</td><td>14961</td><td>AI845868|</td><td>histocomp</td><td>7.091716</td><td>7.441691</td><td>7.408019</td><td>8.279461</td><td>8.320053</td><td>8.505689</td><td>229.3019</td><td>1.091901</td><td>0.001406</td><td>0.019422</td><td>2.131547</td></tr>
<tr><td>H2-AA</td><td>H2-AA</td><td>14960</td><td>Aalpha|H-</td><td>histocomp</td><td>8.015921</td><td>7.760138</td><td>7.84013</td><td>8.927159</td><td>8.923557</td><td>9.006645</td><td>340.6765</td><td>1.10646</td><td>0.000211</td><td>0.009578</td><td>2.153166</td></tr>
<tr><td>VEGFA</td><td>VEGFA</td><td>22339</td><td>Vegf|Vpf</td><td>vascular er</td><td>7.967218</td><td>7.64408</td><td>7.600664</td><td>8.768007</td><td>8.965458</td><td>8.724329</td><td>310.4663</td><td>1.111225</td><td>0.001422</td><td>0.019422</td><td>2.160291</td></tr>
<tr><td>CXCL2</td><td>CXCL2</td><td>20310</td><td>CINC-2a|G</td><td>chemokine</td><td>4.961013</td><td>4.526083</td><td>4.703758</td><td>5.668026</td><td>5.976429</td><td>5.693955</td><td>38.1835</td><td>1.340541</td><td>0.004602</td><td>0.038408</td><td>2.532463</td></tr>
</table>

Primary articular chondrocytes , MDP treated - *Ripk2* [104Asp] vs WT
Nanostring Fibrosis Panel

**Downregulated**

| upperSymb | Symbol | GeneID | Alias | Descriptior | 20201124_ | 20201124_ | 20201124_ | 20201124_ | 20201124_ | 20201124_ | baseMean | log2FoldCh | pvalue | padj | FC |
|---|---|---|---|---|---|---|---|---|---|---|---|---|---|---|---|
| PGM2 | PGM2 | 72157 | 2610020G | phosphogl | 6.818757 | 6.971055 | 6.683723 | 6.168504 | 6.401179 | 6.735479 | 99.02926 | -0.70861 | 0.018239 | 0.202295 | 1.634228 |
| CASP12 | CASP12 | 12364 | - | caspase 12 | 8.842604 | 8.94174 | 9.020536 | 8.113761 | 7.956039 | 8.835015 | 392.9719 | -0.6597 | 0.025515 | 0.222067 | 1.579758 |

**Upregulated**

| | | | | | | | | | | | | | | | |
|---|---|---|---|---|---|---|---|---|---|---|---|---|---|---|---|
| LATS2 | LATS2 | 50523 | 4932411G | large tumo | 7.850466 | 7.794898 | 7.901528 | 8.424487 | 8.345871 | 8.64237 | 286.013 | 0.594855 | 0.000225 | 0.032704 | 1.510321 |
| PHYKPL | PHYKPL | 72947 | 2900006B | 5-phospho | 5.97076 | 6.067271 | 5.96446 | 6.408331 | 6.401179 | 7.176052 | 80.52372 | 0.595213 | 0.003802 | 0.092292 | 1.510696 |
| ACSL4 | ACSL4 | 50790 | 9430020A | acyl-CoA sy | 6.063869 | 5.920429 | 6.388267 | 6.669198 | 6.636282 | 7.176052 | 88.99689 | 0.630743 | 0.014657 | 0.173109 | 1.548362 |
| THBS2 | THBS2 | 21826 | TSP2\|Thbs | thrombosp | 7.95626 | 8.379861 | 8.449258 | 8.74062 | 9.148956 | 8.916809 | 387.6544 | 0.645783 | 0.018517 | 0.202295 | 1.564588 |
| MEF2A | MEF2A | 17258 | A430079H | myocyte e | 7.159794 | 7.221599 | 7.02476 | 7.727657 | 7.88478 | 7.950492 | 180.3739 | 0.676778 | 0.002308 | 0.077579 | 1.598566 |
| SPP1 | SPP1 | 20750 | 2AR\|Apl-1 | secreted pl | 12.92232 | 13.01042 | 13.00351 | 13.9076 | 13.93463 | 12.98544 | 10043.58 | 0.691176 | 0.048712 | 0.284235 | 1.614599 |
| IQGAP1 | IQGAP1 | 29875 | AA682088 | IQ motif cc | 8.965943 | 8.890056 | 8.986286 | 9.609682 | 9.762152 | 9.607604 | 631.9298 | 0.699142 | 0.000807 | 0.061792 | 1.623539 |
| COL6A3 | COL6A3 | 12835 | AI507288\| | collagen_ t | 10.74265 | 10.74243 | 10.90382 | 11.43609 | 11.63933 | 11.45169 | 2276.604 | 0.709218 | 0.001273 | 0.061792 | 1.634917 |
| COL1A2 | COL1A2 | 12843 | AA960264 | collagen_ t | 10.69541 | 10.86406 | 10.97977 | 11.33373 | 11.5615 | 11.80003 | 2361.929 | 0.723442 | 0.004534 | 0.099077 | 1.651116 |
| ITGA5 | ITGA5 | 16402 | Cd49e\|Fnr | integrin alp | 7.116981 | 6.766517 | 6.70494 | 7.778822 | 7.770824 | 7.542834 | 155.4334 | 0.788496 | 0.021882 | 0.212498 | 1.727273 |
| GNB4 | GNB4 | 14696 | 6720453A | guanine nu | 4.648832 | 4.737565 | 5.066339 | 5.168504 | 5.603368 | 6.320442 | 38.25319 | 0.801546 | 0.041423 | 0.270264 | 1.742968 |
| TGFB1 | TGFB1 | 21803 | TGF-beta1 | transformi | 8.004036 | 8.142822 | 8.217933 | 8.951537 | 9.135589 | 8.822942 | 373.7189 | 0.827868 | 0.005547 | 0.105402 | 1.775061 |
| TGFB1I1 | TGFB1I1 | 21804 | ARA55\|Hic | transformi | 5.741941 | 5.528112 | 5.606908 | 6.496558 | 6.239211 | 6.835015 | 67.39755 | 0.835341 | 0.005269 | 0.104667 | 1.784279 |
| FST | FST | 14313 | AL033346\| | follistatin | 5.523301 | 5.256032 | 5.307347 | 6.213828 | 6.120159 | 6.57198 | 56.96911 | 0.848958 | 0.023893 | 0.222067 | 1.8012 |
| COL1A1 | COL1A1 | 12842 | Col1a-1\|Cc | collagen_ t | 10.30132 | 10.44623 | 10.53394 | 11.21661 | 11.31232 | 11.76417 | 1949.786 | 1.020058 | 0.002277 | 0.077579 | 2.028001 |
| LPL | LPL | 16956 | - | lipoprotein | 5.469862 | 6.42293 | 5.928836 | 6.997619 | 6.981879 | 7.451686 | 93.19207 | 1.171953 | 0.009221 | 0.146623 | 2.253165 |
| ACTA2 | ACTA2 | 11475 | 0610041G | actin_ alph | 6.168206 | 6.373602 | 6.221618 | 7.596909 | 7.581 | 7.176052 | 115.592 | 1.210838 | 0.013567 | 0.16469 | 2.314721 |
| LOX | LOX | 16948 | AI893619\| | lysyl oxidas | 10.53473 | 10.99368 | 10.68239 | 12.08178 | 12.07185 | 11.78611 | 2625.582 | 1.237295 | 0.000888 | 0.061792 | 2.35756 |
| CD68 | CD68 | 12514 | Lamp4\|Sca | CD68 antig | 5.696138 | 6.113074 | 6.066339 | 7.153074 | 6.322467 | 8.137578 | 95.76622 | 1.47542 | 0.024477 | 0.222067 | 2.780645 |
| PTGS2 | PTGS2 | 19225 | COX2\|Cox- | prostaglan | 7.008728 | 6.647053 | 7.138289 | 9.228625 | 9.086811 | 7.60055 | 220.5572 | 1.907176 | 0.020209 | 0.20538 | 3.750742 |
| CD36 | CD36 | 12491 | FAT\|GPIV\| | CD36 antig | 4.599922 | 5.615575 | 4.96446 | 6.520806 | 6.41389 | 7.761014 | 63.0873 | 2.176682 | 0.009776 | 0.146623 | 4.521127 |
| C3AR1 | C3AR1 | 12267 | AZ3B\|C3AI | compleme | 4.063869 | 4.435003 | 4.606908 | 6.198878 | 5.19571 | 7.320442 | 39.49144 | 2.756729 | 0.008099 | 0.141578 | 6.758621 |
| SERPINE1 | SERPINE1 | 18787 | PAI-1\|PAI1 | serine (or c | 7.116981 | 7.493896 | 6.89231 | 10.58474 | 10.39958 | 8.513087 | 362.0636 | 2.972579 | 0.005935 | 0.10807 | 7.849383 |

Bone Marrow Derived Macrophages, MDP treated - *Ripk2* $^{104Asp}$ vs WT
Nanostring Fibrosis Panel

**Downregulated**

| upperSyml | Symbol | GeneID | Alias | Descriptior | 20210302_ | 20210302_ | 20210302_ | 20210302_ | 20210302_ | 20210302_ | baseMean | log2FoldCh | pvalue | padj | FC |
|---|---|---|---|---|---|---|---|---|---|---|---|---|---|---|---|
| F11R | F11R | 16456 | 9130004G: | F11 recept | 7.019617 | 6.90166 | 7.042877 | 5.951443 | 5.811373 | 5.886984 | 86.56182 | -1.27422 | 5.89E-06 | 0.002806 | 2.418688 |
| CCNA2 | CCNA2 | 12428 | AA408589 | cyclin A2 | 8.313902 | 8.496471 | 8.482162 | 7.776159 | 7.705504 | 7.661424 | 269.2129 | -0.74817 | 0.000419 | 0.021315 | 1.679655 |
| CDKN2C | CDKN2C | 12580 | C77269\|IN | cyclin-depe | 8.081456 | 8.069388 | 8.030599 | 7.36342 | 7.534663 | 7.311481 | 212.576 | -0.69569 | 0.001123 | 0.026723 | 1.619657 |
| KRAS | KRAS | 16653 | AI929937\| | v-Ki-ras2 K | 5.779566 | 5.861583 | 5.888155 | 5.354202 | 5.170915 | 5.411646 | 47.75823 | -0.69487 | 0.000658 | 0.021315 | 1.618743 |

**Upregulated**

| | | | | | | | | | | | | | | | |
|---|---|---|---|---|---|---|---|---|---|---|---|---|---|---|---|
| PLTP | PLTP | 18830 | Bpife\|OD1 | phospholip | 10.36038 | 10.39571 | 10.43053 | 11.05108 | 11.042 | 10.95873 | 1670.894 | 0.625677 | 6.19E-05 | 0.008373 | 1.542935 |
| MMP12 | MMP12 | 17381 | AV378681 | matrix met | 11.87941 | 12.0139 | 11.90212 | 12.53641 | 12.59125 | 12.6094 | 4889.31 | 0.648633 | 0.000166 | 0.012736 | 1.567682 |
| H2-AB1 | H2-AB1 | 14961 | AI845868\| | histocomp | 7.676472 | 7.466723 | 7.66008 | 8.161557 | 8.283124 | 8.287713 | 242.6296 | 0.671776 | 0.001038 | 0.026012 | 1.593033 |
| H2-AA | H2-AA | 14960 | Aalpha\|H-: | histocomp | 8.081456 | 8.020175 | 8.018215 | 8.74828 | 8.861529 | 8.768339 | 341.6398 | 0.777058 | 7.04E-05 | 0.008373 | 1.713632 |
| FN1 | FN1 | 14268 | E330027I0 | fibronectin | 5.943953 | 6.086875 | 6.018215 | 6.850159 | 6.526396 | 6.69795 | 81.79408 | 0.780345 | 0.00226 | 0.043022 | 1.717542 |

WT primary articular chondrocytes, WEHI-345 and MDP treated vs MDP treated
Nanostring Fibrosis Panel

**Downregulated**

| upperSymb | Symbol | GeneID | Alias | Descriptior | 20201124_ | 20201124_ | 20201124_ | 20210507_ | 20210507_ | 20210507_ | baseMean | log2FoldCh | pvalue | padj | log_10_padj |
|---|---|---|---|---|---|---|---|---|---|---|---|---|---|---|---|
| ANGPTL4 | ANGPTL4 | 57875 | Arp4\|Bk89 | angiopoiet | 6.854204 | 6.722088 | 7.243832 | 3.877416 | 4.034616 | 4.365114 | 45.76623 | -4.23095 | 0.000104 | 0.005274 | -2.2779 |
| MMP13 | MMP13 | 17386 | Clg\|MMP- | matrix met | 11.99399 | 11.91061 | 12.33195 | 8.386651 | 9.006909 | 9.223095 | 1423.803 | -3.27741 | 2.61E-06 | 0.000995 | -3.00239 |
| MKI67 | MKI67 | 17345 | D630048A | antigen ide | 7.14765 | 7.524923 | 7.664407 | 4.877416 | 5.002907 | 6.824545 | 90.94829 | -2.71955 | 3.17E-05 | 0.002496 | -2.6028 |
| CCNA2 | CCNA2 | 12428 | AA408589 | cyclin A2 | 7.250133 | 7.310556 | 7.443856 | 5.324875 | 5.598517 | 6.824545 | 98.7298 | -2.16414 | 3.28E-05 | 0.002496 | -2.6028 |
| FGF21 | FGF21 | 56636 | - | fibroblast g | 6.442778 | 6.58984 | 6.294713 | 4.941546 | 5.211493 | 6.535039 | 64.11403 | -1.91966 | 0.0022 | 0.028668 | -1.5426 |
| NRP2 | NRP2 | 18187 | 1110048P0 | neuropilin | 8.182626 | 8.249765 | 8.516085 | 7.08332 | 7.080909 | 6.950076 | 204.6664 | -1.6552 | 0.003477 | 0.039638 | -1.40189 |
| THBS1 | THBS1 | 21825 | TSP-1\|TSP | thrombosp | 10.93463 | 11.08758 | 11.19301 | 9.423608 | 9.552713 | 9.872908 | 1299.801 | -1.50324 | 8.44E-06 | 0.001218 | -2.91434 |
| PPARG | PPARG | 19016 | Nr1c3\|PPA | peroxisom | 4.928205 | 4.909458 | 5.443856 | 4.06184 | 3.970485 | 6.535039 | 31.4462 | -1.45184 | 0.001713 | 0.025191 | -1.59875 |
| HMOX1 | HMOX1 | 15368 | D8Wsu38e | heme oxyg | 10.51317 | 10.68069 | 10.4195 | 9.406669 | 9.776669 | 9.613041 | 1073.636 | -0.98673 | 0.000237 | 0.008308 | -2.08052 |
| DDIT3 | DDIT3 | 13198 | CHOP-10\|( | DNA-dama | 11.91169 | 12.0083 | 11.75868 | 10.81137 | 10.86585 | 11.1333 | 2730.342 | -0.9741 | 5.55E-05 | 0.003162 | -2.5001 |
| CDKN2C | CDKN2C | 12580 | C77269\|IN | cyclin-depe | 8.116334 | 8.235081 | 8.01588 | 7.366351 | 7.46525 | 7.613041 | 223.1685 | -0.89864 | 0.002505 | 0.031729 | -1.49854 |
| F5 | F5 | 14067 | AI173222\| | coagulatior | 7.442778 | 7.324496 | 7.384694 | 6.547267 | 6.787178 | 7.272004 | 139.7208 | -0.87236 | 0.001395 | 0.024186 | -1.61643 |
| GPC4 | GPC4 | 14735 | 9530073D2 | glypican 4 | 8.641272 | 8.695518 | 8.576081 | 8.058229 | 7.941506 | 7.888676 | 315.2197 | -0.83686 | 0.001591 | 0.024186 | -1.61643 |
| BRAF | BRAF | 109880 | 9930012E1 | Braf transf | 7.723734 | 7.890842 | 7.543392 | 7.166177 | 6.945682 | 7.452577 | 175.3063 | -0.82688 | 0.003122 | 0.036516 | -1.43752 |
| ERN1 | ERN1 | 78943 | 9030414B1 | endoplasm | 8.550466 | 8.560694 | 8.561314 | 7.739912 | 7.850904 | 8.120001 | 300.3599 | -0.81783 | 4.36E-06 | 0.000995 | -3.00239 |
| CHUK | CHUK | 12675 | AI256658\| | conserved | 8.471347 | 8.524923 | 8.519144 | 7.984038 | 7.819393 | 7.888676 | 294.3224 | -0.80714 | 0.00438 | 0.046452 | -1.33299 |
| COL1A1 | COL1A1 | 12842 | Col1a-1\|Co | collagen_ t | 10.3583 | 10.54217 | 10.56354 | 9.4948 | 9.773149 | 9.919703 | 1104.066 | -0.80038 | 0.000713 | 0.015456 | -1.81089 |
| LOXL4 | LOXL4 | 67573 | 4833426I2 | lysyl oxidas | 11.90712 | 12.01073 | 12.15982 | 11.12448 | 11.18613 | 11.41496 | 3177.932 | -0.7983 | 0.000777 | 0.015456 | -1.81089 |
| DDR2 | DDR2 | 18214 | AW495251 | discoidin d | 9.85149 | 9.947111 | 9.85567 | 9.288349 | 9.295689 | 9.065554 | 749.9463 | -0.74556 | 0.001581 | 0.024186 | -1.61643 |
| OGT | OGT | 108155 | 1110038P2 | O-linked N- | 9.406252 | 9.43302 | 9.455084 | 8.893717 | 8.857567 | 8.613041 | 552.4807 | -0.72662 | 0.001859 | 0.02635 | -1.57922 |
| SERPINF1 | SERPINF1 | 20317 | AI195227\| | serine (or c | 10.16524 | 10.29825 | 10.15614 | 9.650405 | 9.583856 | 9.473638 | 947.4602 | -0.69376 | 0.000592 | 0.013502 | -1.86962 |
| COL1A2 | COL1A2 | 12843 | AA960264 | collagen_ t | 10.75239 | 10.96 | 11.00937 | 10.25087 | 10.32884 | 10.28398 | 1549.487 | -0.65044 | 0.000294 | 0.009575 | -2.01884 |

**Upregulated**

| | | | | | | | | | | | | | | | |
|---|---|---|---|---|---|---|---|---|---|---|---|---|---|---|---|
| MAP3K1 | MAP3K1 | 26401 | MAPKKK1\| | mitogen-ac | 6.741436 | 6.73258 | 6.519144 | 7.205886 | 7.259247 | 7.888676 | 133.2349 | 0.699249 | 0.000464 | 0.012991 | -1.88635 |
| H2-K1 | H2-K1 | 14972 | H-2K\|H-2K | histocompa | 8.708815 | 8.73258 | 8.625186 | 9.796279 | 9.773149 | 9.387482 | 576.2623 | 0.954969 | 0.000541 | 0.012991 | -1.88635 |
| SCIN | SCIN | 20259 | AW545522 | scinderin | 7.723734 | 7.737798 | 7.315982 | 8.779928 | 8.713178 | 8.452577 | 278.3069 | 1.015682 | 0.001907 | 0.02635 | -1.57922 |
| GREM1 | GREM1 | 23892 | Cktsf1b1\|D | gremlin 1 | 7.650671 | 7.629681 | 7.437401 | 8.885589 | 8.947765 | 9.065554 | 308.5678 | 1.382837 | 0.000175 | 0.007971 | -2.09849 |
| FASN | FASN | 14104 | A630082H( | fatty acid s | 6.843316 | 6.73258 | 6.806426 | 8.357563 | 8.295689 | 7.950076 | 180.7195 | 1.404713 | 0.000939 | 0.017849 | -1.74839 |
| SCD2 | SCD2 | 20250 | Mir5114\|S | stearoyl-Co | 10.78476 | 10.9942 | 10.63509 | 12.97844 | 12.71578 | 12.28398 | 3401.701 | 1.876792 | 4.42E-05 | 0.002878 | -2.54093 |
| CD68 | CD68 | 12514 | Lamp4\|Sca | CD68 antig | 5.753118 | 6.209018 | 6.095933 | 7.577855 | 8.118384 | 7.950076 | 123.7025 | 1.926937 | 0.000212 | 0.008067 | -2.09329 |
| ELOVL6 | ELOVL6 | 170439 | C77826\|FA | ELOVL fam | 5.225187 | 5.456946 | 5.062766 | 7.300627 | 6.758981 | 7.172469 | 71.64675 | 1.954196 | 0.001568 | 0.024186 | -1.61643 |
| SCD1 | SCD1 | 20249 | AA589638 | stearoyl-Co | 8.94358 | 9.253413 | 8.658869 | 12.12534 | 11.74407 | 11.33666 | 1299.424 | 2.818513 | 1.07E-05 | 0.001218 | -2.91434 |

*Ripk2* $^{104Asp}$ primary articular chondrocytes, WEHI-345 and MDP treated vs MDP treated
Nanostring Fibrosis Panel

**Downregulated**

| upperSymb | Symbol | GeneID | Alias | Descriptior | 20201124_ | 20201124_ | 20201124_ | 20210507_ | 20210507_ | 20210507_ | baseMean | log2FoldCh | pvalue | padj | log 10 |
|---|---|---|---|---|---|---|---|---|---|---|---|---|---|---|---|
| LPL | LPL | 16956 | - | lipoprotein | 6.61791 | 7.068903 | 6.573375 | 3.54908 | 4.386761 | 4.676842 | 44.59505 | -2.65588 | 0.000145 | 0.017118 | -1.76655 |
| VEGFA | VEGFA | 22339 | Vegf\|Vpf | vascular er | 7.796448 | 8.988284 | 7.71939 | 6.66882 | 6.741294 | 6.571659 | 170.5813 | -1.61461 | 0.001779 | 0.041356 | -1.38346 |
| CYP27A1 | CYP27A1 | 104086 | 1300013A( | cytochrom | 5.726059 | 6.754794 | 5.457897 | 4.54908 | 4.211674 | 4.320698 | 36.00271 | -1.61382 | 0.000412 | 0.025912 | -1.5865 |
| ACTA2 | ACTA2 | 11475 | 0610041G( | actin_ alph | 7.217199 | 6.793268 | 7.172495 | 5.63 | 5.323567 | 5.431729 | 76.7118 | -1.5603 | 0.001273 | 0.037897 | -1.4214 |
| COL1A1 | COL1A1 | 12842 | Col1a-1\|Cc | collagen_ t | 10.8369 | 11.38139 | 10.90381 | 9.414695 | 9.794696 | 9.526247 | 1269.13 | -1.47048 | 0.000919 | 0.037897 | -1.4214 |
| TNN | TNN | 329278 | Tnw\|tenas | tenascin N | 5.643597 | 6.754794 | 6.352404 | 4.936103 | 4.716909 | 4.721236 | 45.91331 | -1.41504 | 0.001697 | 0.041356 | -1.38346 |
| OGT | OGT | 108155 | 1110038P2 | O-linked N | 9.022108 | 9.632804 | 8.729001 | 7.712827 | 7.757324 | 7.95993 | 354.3421 | -1.34746 | 0.00188 | 0.041356 | -1.38346 |
| CXCL12 | CXCL12 | 20315 | Pbsf\|Scyb1 | chemokine | 9.545954 | 8.600623 | 9.55079 | 7.842994 | 8.152296 | 7.980622 | 391.3222 | -1.28225 | 0.002304 | 0.048154 | -1.31737 |
| FABP4 | FABP4 | 11770 | 422/aP2\|A | fatty acid b | 6.626523 | 7.300228 | 7.44862 | 5.706622 | 5.998945 | 5.876514 | 90.06587 | -1.22367 | 0.002408 | 0.048154 | -1.31737 |
| DDIT3 | DDIT3 | 13198 | CHOP-10\|( | DNA-dama | 11.27757 | 10.95473 | 11.16649 | 9.997117 | 9.863883 | 10.10834 | 1511.072 | -1.13692 | 4.89E-05 | 0.017118 | -1.76655 |
| THBS2 | THBS2 | 21826 | TSP2\|Thbs | thrombosp | 8.36091 | 8.534025 | 8.740451 | 7.455971 | 7.514866 | 7.298893 | 253.2092 | -1.09742 | 0.001148 | 0.037897 | -1.4214 |
| LOX | LOX | 16948 | AI893619\| | lysyl oxidas | 11.70207 | 11.40332 | 11.66335 | 10.50774 | 10.77859 | 10.41662 | 2162.695 | -1.01556 | 0.000869 | 0.037897 | -1.4214 |
| COL1A2 | COL1A2 | 12843 | AA960264 | collagen_ t | 10.95402 | 11.41725 | 11.15299 | 10.1818 | 10.28607 | 10.19893 | 1661.776 | -0.95861 | 0.00115 | 0.037897 | -1.4214 |
| ERN1 | ERN1 | 78943 | 9030414B1 | endoplasm | 8.292197 | 8.643373 | 7.917329 | 7.562886 | 7.274286 | 7.370329 | 229.667 | -0.86381 | 0.001378 | 0.037897 | -1.4214 |
| ARHGEF2 | ARHGEF2 | 16800 | AA408978 | rho/rac gu | 9.142598 | 9.390831 | 8.930718 | 8.228019 | 8.263102 | 8.351034 | 421.0119 | -0.85998 | 0.000685 | 0.037676 | -1.42393 |
| COL5A1 | COL5A1 | 12831 | AI413331 | collagen_ t | 10.65942 | 11.16373 | 10.66234 | 10.04831 | 9.99557 | 9.920016 | 1358.908 | -0.8519 | 0.001863 | 0.041356 | -1.38346 |

**Upregulated**

| | | | | | | | | | | | | | | | |
|---|---|---|---|---|---|---|---|---|---|---|---|---|---|---|---|
| COL10A1 | COL10A1 | 12813 | Col10\|Col1 | collagen_ t | 10.0057 | 10.00477 | 10.17812 | 11.28795 | 11.61475 | 11.37533 | 1715.528 | 1.390271 | 9.05E-05 | 0.017118 | -1.76655 |
| CHIL1 | CHIL1 | 12654 | AW20876( | chitinase-li | 9.052788 | 9.83536 | 8.403716 | 11.16322 | 11.15267 | 11.35522 | 1144.496 | 2.054077 | 0.000342 | 0.025115 | -1.60008 |

WT primary articular chondrocytes - MDP treated vs control
Nanostring Fibrosis Panel

**<u>Downregulated</u>**

| upperSyml | Symbol | GeneID | Alias | Descriptior | 20210507_ | 20210507_ | 20210507_ | 20201124_ | 20201124_ | 20201124_ | baseMean | log2FoldCh | pvalue | padj | FC |
|---|---|---|---|---|---|---|---|---|---|---|---|---|---|---|---|
| CD84 | CD84 | 12523 | A130013D: | CD84 antig | 4.39874 | 5.774494 | 5.477158 | 2.819218 | 3.861278 | 4.022416 | 20.99854 | -4.96963 | 0.007009 | 0.026965 | 31.33333 |
| FGL2 | FGL2 | 14190 | AI385601\| | fibrinogen- | 5.316278 | 5.552102 | 5.114588 | 2.819218 | 3.754363 | 2.92288 | 18.98215 | -4.96323 | 1.20E-05 | 0.002593 | 31.19466 |
| PPARGC1A | PPARGC1A | 19017 | A830037N | peroxisom | 4.643852 | 4.967139 | 4.685744 | 3.04161 | 3.223848 | 3.700488 | 16.49298 | -4.1971 | 1.34E-05 | 0.002593 | 18.3423 |
| ITGB3 | ITGB3 | 16416 | CD61\|GP3. | integrin be | 5.103284 | 4.552102 | 4.740192 | 4.234255 | 3.513355 | 3.92288 | 20.31319 | -3.56071 | 0.006629 | 0.025909 | 11.8 |
| CSF1R | CSF1R | 12978 | AI323359\| | colony stin | 4.853306 | 5.521728 | 5.444736 | 2.819218 | 4.577485 | 4.022416 | 23.26057 | -3.07973 | 0.007675 | 0.028019 | 8.454545 |
| CYBB | CYBB | 13058 | C88302\|C( | cytochrom | 4.643852 | 5.289068 | 4.685744 | 3.234255 | 4.513355 | 3.28545 | 19.36376 | -2.67807 | 0.015397 | 0.042728 | 6.4 |
| CD68 | CD68 | 12514 | Lamp4\|Sca | CD68 antig | 8.704548 | 8.912583 | 8.316222 | 6.188452 | 6.638886 | 6.574957 | 188.1763 | -2.4366 | 0.000305 | 0.009526 | 5.413636 |
| CD36 | CD36 | 12491 | FAT\|GPIV\| | CD36 antig | 6.550743 | 7.755042 | 7.377622 | 5.092236 | 6.141386 | 5.473077 | 84.35204 | -2.14136 | 0.010515 | 0.034009 | 4.411765 |
| TLR2 | TLR2 | 24088 | Ly105 | toll-like rec | 5.425212 | 5.359457 | 5.114588 | 4.321718 | 4.754363 | 4.759381 | 31.03419 | -1.88139 | 0.015333 | 0.042728 | 3.684292 |
| ABCA1 | ABCA1 | 11303 | ABC-1\|Abc | ATP-bindin | 6.621132 | 6.69506 | 6.614661 | 5.278649 | 5.861278 | 4.870413 | 63.56668 | -1.81263 | 0.007072 | 0.026965 | 3.512821 |
| FNIP2 | FNIP2 | 329679 | D630023B: | folliculin in | 9.042596 | 8.938683 | 8.643516 | 7.164993 | 7.253596 | 7.30535 | 266.5241 | -1.76411 | 0.000177 | 0.007533 | 3.396634 |
| DLL1 | DLL1 | 13388 | Delta1 | delta-like 1 | 5.425212 | 5.137064 | 4.843286 | 4.693687 | 4.375851 | 4.700488 | 29.09296 | -1.69599 | 0.008451 | 0.029577 | 3.24 |
| NR1H3 | NR1H3 | 22259 | AU018371 | nuclear rec | 6.330353 | 6.596496 | 6.614661 | 5.234255 | 5.411475 | 5.54179 | 62.0276 | -1.64955 | 0.000463 | 0.010605 | 3.137358 |
| JAG1 | JAG1 | 16449 | ABE2\|Gsfa | jagged 1 | 5.773135 | 6.033229 | 6.114588 | 4.40418 | 5.223848 | 5.324979 | 44.60066 | -1.64829 | 0.00868 | 0.029935 | 3.134615 |
| MET | MET | 17295 | AI838057\| | met proto- | 9.181071 | 8.969954 | 8.927819 | 7.600578 | 7.638886 | 7.700488 | 323.2409 | -1.47273 | 6.77E-05 | 0.004535 | 2.77546 |
| H2-T23 | H2-T23 | 15040 | 37b\|37c\|H | histocompa | 6.813777 | 6.886003 | 6.508867 | 5.788844 | 6.031203 | 5.244808 | 74.1436 | -1.4021 | 0.007275 | 0.027515 | 2.642857 |
| GAS1 | GAS1 | 14451 | AW554192 | growth arr | 12.98064 | 12.98395 | 13.48739 | 11.84943 | 11.79033 | 11.72378 | 5670.464 | -1.3873 | 0.000644 | 0.010744 | 2.61588 |
| ICAM1 | ICAM1 | 15894 | CD54\|Icam | intercellula | 8.206095 | 8.200759 | 8.419933 | 7.245483 | 6.861278 | 6.998168 | 201.5908 | -1.38481 | 0.000669 | 0.010744 | 2.611382 |
| LGALS3 | LGALS3 | 16854 | GBP\|L-34\| | lectin_ gala | 8.221281 | 8.006058 | 8.067454 | 6.962176 | 6.848339 | 6.935704 | 181.879 | -1.33696 | 6.25E-05 | 0.004535 | 2.526191 |
| H2-AB1 | H2-AB1 | 14961 | AI845868\| | histocompa | 7.823761 | 7.674499 | 7.46912 | 6.501042 | 6.697779 | 6.607378 | 139.9657 | -1.24409 | 0.000729 | 0.010744 | 2.368683 |
| IRAK3 | IRAK3 | 73914 | 4833428C: | interleukin | 5.872671 | 5.426571 | 5.508867 | 5.092236 | 5.263377 | 4.815965 | 40.22297 | -1.17585 | 0.012532 | 0.038166 | 2.259259 |
| ADCY7 | ADCY7 | 11513 | AA407758 | adenylate | 4.813777 | 4.82512 | 5.307233 | 4.693687 | 4.697779 | 4.700488 | 28.63446 | -1.16684 | 0.004735 | 0.022433 | 2.245197 |
| EGR1 | EGR1 | 13653 | A530045N | early grow | 6.731315 | 6.65364 | 6.69955 | 5.906681 | 6.20367 | 5.54179 | 78.21867 | -1.15122 | 0.016322 | 0.043994 | 2.221012 |
| NCEH1 | NCEH1 | 320024 | Aadacl1\|B. | neutral chc | 6.853306 | 6.989507 | 7.223709 | 5.989143 | 5.911904 | 6.490565 | 95.42899 | -1.14066 | 0.003567 | 0.020324 | 2.204819 |
| HMOX1 | HMOX1 | 15368 | D8Wsu38e | heme oxyg | 12.29759 | 12.13142 | 11.92268 | 10.9485 | 11.11056 | 10.89852 | 3001.662 | -1.13976 | 0.000847 | 0.010744 | 2.203439 |
| PTGS2 | PTGS2 | 19225 | COX2\|Cox- | prostaglan | 8.720668 | 8.494616 | 8.150212 | 7.501042 | 7.172864 | 7.646907 | 246.8889 | -1.11849 | 0.003804 | 0.020324 | 2.17119 |
| SERPINE1 | SERPINE1 | 18787 | PAI-1\|PAI1 | serine (or c | 8.744514 | 9.116887 | 8.150212 | 7.609295 | 8.019707 | 7.400927 | 288.7326 | -1.10602 | 0.018238 | 0.047085 | 2.152513 |
| TRIB3 | TRIB3 | 228775 | Ifld2\|Nipk\| | tribbles ho | 6.501833 | 6.625351 | 6.477158 | 5.726108 | 5.984661 | 5.639087 | 71.45846 | -1.10119 | 0.003189 | 0.019427 | 2.145308 |
| FAS | FAS | 14102 | AI196731\| | Fas (TNF re | 6.451207 | 5.69506 | 6.428248 | 5.693687 | 5.339325 | 5.670114 | 58.87597 | -1.05166 | 0.018272 | 0.047085 | 2.072917 |
| PLCG2 | PLCG2 | 234779 | PLCgamma | phospholip | 7.302064 | 7.27092 | 7.279925 | 6.66052 | 6.768169 | 6.159919 | 120.0024 | -0.95099 | 0.017447 | 0.046493 | 1.933204 |
| PSEN2 | PSEN2 | 19165 | ALG-3\|Ad4 | presenilin : | 6.910639 | 7.106691 | 6.996532 | 6.574105 | 6.183206 | 6.092805 | 100.0097 | -0.95085 | 0.008849 | 0.030292 | 1.933008 |
| RELB | RELB | 19698 | shep | avian retic | 7.704548 | 7.450955 | 7.444736 | 6.757817 | 6.668633 | 6.948415 | 143.2625 | -0.89167 | 0.002199 | 0.015391 | 1.855321 |
| EGFR | EGFR | 13649 | 9030024J1 | epidermal | 7.273209 | 7.186308 | 7.279925 | 6.42408 | 6.808811 | 6.400927 | 119.0599 | -0.88708 | 0.006321 | 0.02575 | 1.849431 |
| NUMB | NUMB | 18222 | Nb | numb gene | 5.688246 | 5.967139 | 5.79266 | 5.188452 | 5.577485 | 5.400927 | 48.58654 | -0.87177 | 0.014386 | 0.041648 | 1.829901 |
| CEBPA | CEBPA | 12606 | C/ebpalph | CCAAT/enh | 6.882257 | 7.289068 | 7.508867 | 6.42408 | 6.726349 | 6.607378 | 119.9536 | -0.82933 | 0.011217 | 0.035072 | 1.77686 |
| TRAF2 | TRAF2 | 22030 | AI325259 | TNF recept | 8.175234 | 7.897877 | 7.830795 | 7.129073 | 7.311311 | 7.295434 | 194.9041 | -0.82861 | 0.003062 | 0.019149 | 1.775972 |
| STAT3 | STAT3 | 20848 | 1110034C( | signal trans | 9.978125 | 9.911119 | 9.694388 | 9.054434 | 9.008119 | 9.170808 | 708.9299 | -0.81213 | 0.001245 | 0.012 | 1.755798 |
| DAPK1 | DAPK1 | 69635 | D13Ucla1\| | death asso | 6.793599 | 7.166812 | 6.614661 | 6.538036 | 6.008119 | 6.265272 | 94.64253 | -0.80183 | 0.019504 | 0.0484 | 1.743316 |
| PCX | PCX | 18563 | Pc\|Pcb | pyruvate c | 7.236309 | 7.26176 | 7.436516 | 6.574105 | 6.726349 | 6.759381 | 127.9175 | -0.78258 | 0.000893 | 0.010744 | 1.720207 |
| LIPA | LIPA | 16889 | AA960673 | lysosomal | 7.813777 | 8.257158 | 8.150212 | 7.245483 | 7.393773 | 7.599341 | 214.2707 | -0.7597 | 0.006371 | 0.02575 | 1.693133 |
| MTMR4 | MTMR4 | 170749 | AA596759 | myotubula | 7.03617 | 6.932924 | 6.818195 | 6.164993 | 6.428963 | 6.507843 | 100.3002 | -0.75928 | 0.006424 | 0.02575 | 1.692642 |
| NOTCH1 | NOTCH1 | 18128 | 9930111A: | notch 1 | 6.198441 | 6.156964 | 6.194758 | 5.66052 | 5.808811 | 5.815965 | 62.79494 | -0.73063 | 0.000517 | 0.010744 | 1.659365 |
| H2-M3 | H2-M3 | 14991 | H-2M3\|Hn | histocompa | 5.910639 | 5.897877 | 5.843286 | 5.626573 | 5.608512 | 5.363453 | 52.28734 | -0.72723 | 0.012402 | 0.038017 | 1.655462 |
| TRAF6 | TRAF6 | 22034 | 2310003F1 | TNF recept | 6.731315 | 6.989507 | 6.79266 | 6.164993 | 6.393773 | 6.419306 | 95.79813 | -0.71204 | 0.00528 | 0.024041 | 1.638118 |
| OSBPL5 | OSBPL5 | 79196 | 1110006M | oxysterol b | 9.296697 | 9.382881 | 9.054083 | 8.596199 | 8.593082 | 8.583131 | 483.6028 | -0.69254 | 0.002422 | 0.016703 | 1.61613 |
| LGMN | LGMN | 19141 | AEP\|AI746 | legumain | 7.89178 | 8.111798 | 8.083337 | 7.591807 | 7.375851 | 7.344344 | 212.7703 | -0.68043 | 0.003644 | 0.020324 | 1.60262 |
| CFLAR | CFLAR | 12633 | 2310024N: | CASP8 and | 7.720668 | 8.101566 | 7.927819 | 7.311071 | 7.393773 | 7.315198 | 197.8618 | -0.67082 | 0.004701 | 0.022433 | 1.591981 |
| MEF2A | MEF2A | 17258 | A430079H | myocyte er | 8.171329 | 8.410083 | 8.124856 | 7.652108 | 7.74741 | 7.533378 | 245.5479 | -0.66704 | 0.004703 | 0.022433 | 1.587816 |
| CDKN1A | CDKN1A | 12575 | CAP20\|CD | cyclin-depe | 10.39205 | 10.62535 | 10.67355 | 9.899591 | 9.965315 | 9.943662 | 1217.68 | -0.64247 | 0.001963 | 0.014852 | 1.560996 |
| KIF3A | KIF3A | 16568 | Kif3\|Kifl\|K | kinesin fan | 6.167414 | 6.195958 | 6.461038 | 5.906681 | 5.936566 | 5.870413 | 68.10451 | -0.63837 | 0.003493 | 0.020225 | 1.556575 |
| ZFYVE16 | ZFYVE16 | 218441 | AI035632\| | zinc finger_ | 8.287708 | 8.166812 | 7.726772 | 7.353196 | 7.631352 | 7.591259 | 221.7591 | -0.63254 | 0.019361 | 0.048312 | 1.550296 |
| ANAPC7 | ANAPC7 | 56317 | APC7\|AW5 | anaphase p | 6.330353 | 6.342179 | 6.051394 | 5.726108 | 5.911904 | 5.998168 | 66.71863 | -0.63129 | 0.016617 | 0.044534 | 1.548947 |
| LOXL1 | LOXL1 | 16949 | Loxl | lysyl oxida: | 9.344292 | 9.572619 | 9.407369 | 8.881714 | 8.996438 | 8.685381 | 567.3003 | -0.61893 | 0.006375 | 0.02575 | 1.535731 |
| IRS1 | IRS1 | 16367 | G972R\|IRS | insulin rece | 8.451207 | 8.186308 | 8.501004 | 7.609295 | 7.768169 | 8.098519 | 274.8341 | -0.61029 | 0.013497 | 0.040007 | 1.526563 |
| ATG101 | ATG101 | 68118 | 9430023L2 | autophagy | 6.476742 | 6.27092 | 6.394696 | 6.188452 | 5.886813 | 6.022416 | 73.85753 | -0.58529 | 0.01093 | 0.034405 | |

**<u>Upregulated</u>**

| | | | | | | | | | | | | | | | |
|---|---|---|---|---|---|---|---|---|---|---|---|---|---|---|---|
| GRB10 | GRB10 | 14783 | 5730571D( | growth fac | 8.933871 | 8.81263 | 8.996532 | 9.467875 | 9.633239 | 9.414733 | 592.1477 | 0.585594 | 0.002543 | 0.016795 | 1.500657 |
| RAC1 | RAC1 | 19353 | AL023026\| | RAS-relate | 10.09815 | 10.19836 | 10.43548 | 10.82671 | 10.88444 | 10.82375 | 1493.501 | 0.598967 | 0.004203 | 0.021862 | 1.514632 |
| NRP2 | NRP2 | 18187 | 1110048P( | neuropilin | 7.929254 | 8.091261 | 8.403157 | 8.61796 | 8.679632 | 8.995108 | 350.3684 | 0.609693 | 0.015868 | 0.043268 | 1.525935 |
| PDGFRB | PDGFRB | 18596 | AI528809\| | platelet de | 6.03617 | 6.156964 | 6.029699 | 6.677199 | 6.623779 | 6.88371 | 84.5219 | 0.624255 | 0.003286 | 0.019761 | 1.541414 |
| TLN1 | TLN1 | 21894 | Tln | talin 1 | 8.441514 | 8.701849 | 8.55516 | 9.197151 | 9.233832 | 9.186988 | 473.1266 | 0.634615 | 0.001253 | 0.012 | 1.552524 |
| FN1 | FN1 | 14268 | E330027I0 | fibronectin | 14.9607 | 15.11966 | 15.043 | 15.5449 | 15.77838 | 15.73087 | 42140.49 | 0.643541 | 0.001615 | 0.013249 | 1.562158 |
| SKP1A | SKP1A | 21402 | 15kDa\|261 | S-phase kir | 8.972527 | 9.169263 | 9.15523 | 9.69778 | 9.756096 | 9.79852 | 687.3507 | 0.648401 | 0.000738 | 0.010744 | 1.56743 |
| TBL1XR1 | TBL1XR1 | 81004 | 8030499H( | transducin | 7.127668 | 7.306989 | 7.650641 | 7.975722 | 8.031203 | 8.075527 | 207.1634 | 0.654683 | 0.014534 | 0.041789 | 1.57427 |
| HSBP1 | HSBP1 | 68196 | 0610007A( | heat shock | 7.848423 | 8.038602 | 8.140123 | 8.538036 | 8.788633 | 8.72655 | 325.5483 | 0.668634 | 0.004447 | 0.02219 | 1.589567 |
| VIM | VIM | 22352 | - | vimentin | 12.10277 | 12.40775 | 12.22699 | 12.91418 | 13.06463 | 12.98608 | 6282.25 | 0.742268 | 0.001679 | 0.013345 | 1.672803 |
| CTNNB1 | CTNNB1 | 12387 | Bfc\|Catnb\| | catenin (ca | 11.08108 | 11.20853 | 11.23797 | 11.88441 | 11.97836 | 11.93091 | 3005.822 | 0.754997 | 0.000166 | 0.007533 | 1.687628 |
| AMOTL2 | AMOTL2 | 56332 | AW54973¢ | angiomotir | 7.095063 | 7.529382 | 7.477158 | 7.975722 | 8.282742 | 8.192341 | 216.5768 | 0.779159 | 0.010289 | 0.03351 | 1.71613 |
| LOXL2 | LOXL2 | 94352 | 1110004B( | lysyl oxida: | 9.900061 | 10.1333 | 10.03515 | 10.73709 | 10.93962 | 10.82375 | 1377.81 | 0.810315 | 0.000847 | 0.010744 | 1.753594 |

| | | | | | | | | | | | | | | | |
|---|---|---|---|---|---|---|---|---|---|---|---|---|---|---|---|
| PRDX1 | PRDX1 | 18477 | MSP23\|Nk | peroxiredo | 6.344292 | 6.735324 | 6.508867 | 7.245483 | 7.402651 | 7.39165 | 122.6195 | 0.814737 | 0.003761 | 0.020324 | 1.758977 |
| CALM1 | CALM1 | 12313 | AI256814\| | calmodulin | 7.309188 | 7.735324 | 7.849491 | 8.337543 | 8.553769 | 8.455375 | 263.2181 | 0.816032 | 0.010639 | 0.034176 | 1.760558 |
| F5 | F5 | 14067 | AI173222\| | coagulatio | 6.90124 | 6.944419 | 7.175129 | 7.878112 | 7.754363 | 7.863718 | 171.195 | 0.823247 | 0.000874 | 0.010744 | 1.769384 |
| MAF | MAF | 17132 | 2810401A2 | avian musc | 5.287708 | 5.552102 | 5.600015 | 6.519658 | 6.098317 | 6.437453 | 60.37484 | 0.875669 | 0.008562 | 0.029746 | 1.834859 |
| CKAP4 | CKAP4 | 216197 | 5630400A0 | cytoskeleto | 9.512675 | 9.723696 | 9.898194 | 10.62227 | 10.65198 | 10.58313 | 1148.332 | 0.908162 | 0.001321 | 0.012394 | 1.876652 |
| COL1A2 | COL1A2 | 12843 | AA960264 | collagen_ t | 10.2967 | 10.52365 | 10.51669 | 11.18772 | 11.38987 | 11.48839 | 1911.517 | 0.910019 | 0.001424 | 0.012845 | 1.87907 |
| GCNT1 | GCNT1 | 14537 | 5630400D2 | glucosamir | 6.562715 | 6.625351 | 6.867948 | 7.40418 | 7.661253 | 7.737577 | 141.3542 | 0.921025 | 0.002898 | 0.018369 | 1.893459 |
| FLNB | FLNB | 286940 | AL024016\| | filamin_ be | 8.001404 | 8.166812 | 8.302717 | 8.995807 | 9.114619 | 9.186988 | 395.6436 | 0.944931 | 0.000796 | 0.010744 | 1.925097 |
| CFHR2 | CFHR2 | 545366 | Cfhrb_4/2 | compleme | 6.135705 | 6.596496 | 6.477158 | 7.574105 | 7.273092 | 7.159919 | 116.9228 | 0.945431 | 0.005014 | 0.023137 | 1.925764 |
| CCN2 | CCN2 | 14219 | Ccn2\|Fisp1 | connective | 12.63962 | 12.94066 | 13.14565 | 13.90225 | 13.88602 | 13.91541 | 10846.45 | 0.97808 | 0.001192 | 0.012 | 1.969843 |
| THBS2 | THBS2 | 21826 | TSP2\|Thbs | thrombosp | 7.294904 | 8.132046 | 7.811853 | 8.448574 | 8.905672 | 8.957876 | 306.2334 | 1.007918 | 0.013563 | 0.040007 | 2.011006 |
| LEPR | LEPR | 16847 | LEPROT\|Le | leptin rece | 4.731315 | 5.82512 | 5.539894 | 6.538036 | 6.375851 | 6.363453 | 59.53274 | 1.013806 | 0.018993 | 0.047891 | 2.019231 |
| SCD2 | SCD2 | 20250 | Mir5114\|S | stearoyl-Co | 9.938473 | 10.07697 | 10.42928 | 11.2201 | 11.42407 | 11.11412 | 1664.071 | 1.097026 | 0.001487 | 0.012853 | 2.139133 |
| FASN | FASN | 14104 | A630082H | fatty acid s | 6.198441 | 6.054602 | 6.252091 | 7.278649 | 7.162448 | 7.28545 | 104.3495 | 1.102906 | 0.000111 | 0.006533 | 2.147869 |
| FABP4 | FABP4 | 11770 | 422/aP2\|A | fatty acid b | 6.385319 | 7.022422 | 6.508867 | 7.453424 | 8.233832 | 7.903428 | 152.3468 | 1.259583 | 0.0092 | 0.030477 | 2.394265 |
| ACTA2 | ACTA2 | 11475 | 0610041G0 | actin_ alph | 5.598048 | 5.82512 | 5.270707 | 6.66052 | 6.899413 | 6.730235 | 71.70528 | 1.278544 | 0.00589 | 0.025406 | 2.425941 |
| FZD2 | FZD2 | 57265 | AL033370\| | frizzled ho | 5.001404 | 5.521728 | 5.685744 | 6.643646 | 6.623779 | 6.54179 | 64.1339 | 1.280966 | 0.00619 | 0.02575 | 2.430017 |
| COL1A1 | COL1A1 | 12842 | Col1a-1\|Cc | collagen_ t | 9.363241 | 9.743572 | 9.647083 | 10.79363 | 10.97204 | 11.04256 | 1226.52 | 1.356357 | 0.000581 | 0.010744 | 2.560378 |
| RBX1 | RBX1 | 56438 | 1500002P1 | ring-box 1 | 6.018892 | 6.506298 | 6.643516 | 7.547138 | 7.942666 | 7.909941 | 136.6879 | 1.453118 | 0.001385 | 0.012733 | 2.737991 |
| ELOVL6 | ELOVL6 | 170439 | C77826\|FA | ELOVL fam | 4.03617 | 4.552102 | 4.570267 | 5.66052 | 5.886813 | 5.54179 | 32.92878 | 1.568184 | 0.006509 | 0.025871 | 2.965312 |
| CDKN2C | CDKN2C | 12580 | C77269\|IN | cyclin-depe | 6.947633 | 7.065172 | 6.855669 | 8.551668 | 8.664948 | 8.494904 | 217.268 | 1.65942 | 4.99E-05 | 0.004535 | 3.158895 |
| LOXL4 | LOXL4 | 67573 | 4833426I2 | lysyl oxidas | 10.39957 | 10.89862 | 10.99233 | 12.34245 | 12.44059 | 12.63884 | 3144.762 | 1.696413 | 0.000475 | 0.010605 | 3.240941 |
| ATF7IP | ATF7IP | 54343 | 2610204M | activating t | 6.213708 | 6.011534 | 6.093829 | 7.841586 | 7.893127 | 7.909941 | 127.4647 | 1.872842 | 1.66E-05 | 0.002593 | 3.662533 |
| CCNA2 | CCNA2 | 12428 | AA408589 | cyclin A2 | 5.526495 | 6.011534 | 5.939501 | 7.685466 | 7.740424 | 7.92288 | 111.7695 | 2.092746 | 0.000327 | 0.009526 | 4.265591 |
| ANGPTL4 | ANGPTL4 | 57875 | Arp4\|Bk89 | angiopoiet | 4.813777 | 5.289068 | 6.051394 | 7.289538 | 7.151955 | 7.722856 | 83.65798 | 2.110282 | 0.003942 | 0.020774 | 4.317757 |
| MKI67 | MKI67 | 17345 | D630048A | antigen ide | 5.070117 | 6.033229 | 5.867948 | 7.582984 | 7.95479 | 8.143431 | 109.5478 | 2.365321 | 0.001016 | 0.011618 | 5.152672 |
| THBS1 | THBS1 | 21825 | TSP-1\|TSP | thrombosp | 8.945348 | 9.154492 | 9.15523 | 11.36996 | 11.51745 | 11.67203 | 1262.807 | 2.450422 | 2.57E-05 | 0.00301 | 5.46576 |
| MMP13 | MMP13 | 17386 | Clg\|MMP-1 | matrix met | 9.677275 | 10.06517 | 10.29023 | 12.42932 | 12.34048 | 12.81097 | 2467.625 | 2.517348 | 0.000176 | 0.007533 | 5.725289 |
| POSTN | POSTN | 50706 | A630052E0 | periostin_ | 3.344292 | 3.289068 | 3.892195 | 6.66052 | 6.712135 | 7.446442 | 37.37777 | 5.379378 | 0.000865 | 0.010744 | 41.625 |

*Rlpk2* $^{104Asp}$ primary articular chondrocytes - MDP treated vs control
Nanostring Fibrosis Panel

**Downregulated**

| upperSymb | Symbol | GeneID | Alias | Descriptior | 20210302_ | 20210302_ | 20210302_ | 20201124_ | 20201124_ | 20201124_ | baseMean | log2FoldCh | pvalue | padj | FC | |
|---|---|---|---|---|---|---|---|---|---|---|---|---|---|---|---|---|
| CHIL1 | CHIL1 | 12654 | AW208766 | chitinase-li | 10.97263 | 10.57922 | 10.51809 | 8.670058 | 8.049781 | 9.455703 | 836.1274 | -1.895 | 0.004951 | 0.178947 | 3.719219 | downregulated |
| GAS1 | GAS1 | 14451 | AW554192 | growth arr | 11.91347 | 12.04885 | 11.65735 | 10.12411 | 10.01024 | 11.12858 | 2267.834 | -1.375648 | 0.01097 | 0.178947 | 2.594843 | |
| VEGFB | VEGFB | 22340 | VEGF-B\|Vr | vascular er | 6.06752 | 6.010978 | 5.787576 | 4.136878 | 4.772756 | 5.750647 | 42.84512 | -1.321262 | 0.012262 | 0.180963 | 2.498846 | |
| COL10A1 | COL10A1 | 12813 | Col10\|Col1 | collagen_ t | 10.79846 | 11.15162 | 10.612 | 9.622965 | 9.824184 | 9.625116 | 1236.796 | -1.189946 | 0.00128 | 0.178947 | 2.281442 | |
| ICAM1 | ICAM1 | 15894 | CD54\|Icam | intercellula | 7.24337 | 7.351207 | 6.821523 | 6.592942 | 5.958111 | 6.375138 | 105.6915 | -0.914122 | 0.022327 | 0.230675 | 1.884422 | |
| CCNA2 | CCNA2 | 12428 | AA408589\| | cyclin A2 | 6.956489 | 7.041352 | 6.821523 | 6.269328 | 6.056965 | 6.375138 | 96.12228 | -0.800522 | 0.007329 | 0.178947 | 1.741732 | |
| FAP | FAP | 14089 | SIMP | fibroblast a | 8.931018 | 9.167196 | 8.726175 | 8.031384 | 8.29834 | 8.345594 | 383.5536 | -0.73348 | 0.008761 | 0.178947 | 1.662645 | |
| CASP12 | CASP12 | 12364 | - | caspase 12 | 8.176819 | 8.393027 | 8.271775 | 7.351321 | 7.193599 | 8.072575 | 240.4933 | -0.718546 | 0.019204 | 0.219101 | 1.645522 | |
| LEPR | LEPR | 16847 | LEPROT\|Le | leptin rece | 6.312632 | 6.500784 | 6.123179 | 5.436438 | 5.625915 | 6.023665 | 64.16741 | -0.713431 | 0.01537 | 0.196958 | 1.639699 | |
| MMP3 | MMP3 | 17392 | EMS-2\|MM | matrix met | 7.195275 | 7.085746 | 6.770298 | 6.458806 | 6.131443 | 6.750647 | 106.3028 | -0.602899 | 0.028149 | 0.250993 | 1.518765 | |
| F5 | F5 | 14067 | AI173222\| | coagulatior | 7.588757 | 7.778318 | 7.482721 | 6.72184 | 6.743372 | 7.540724 | 158.6044 | -0.598408 | 0.028829 | 0.251814 | 1.514045 | |
| STAT1 | STAT1 | 20846 | 2010005J0 | signal trans | 6.040553 | 6.041352 | 5.91882 | 5.136878 | 5.028013 | 6.119881 | 52.50014 | -0.587139 | 0.015869 | 0.196958 | 1.502265 | |

**Upregulated**

| | | | | | | | | | | | | | | | | |
|---|---|---|---|---|---|---|---|---|---|---|---|---|---|---|---|---|
| RPS6KA2 | RPS6KA2 | 20112 | 90kDa\|D17 | ribosomal | 5.219523 | 5.314371 | 5.150146 | 5.191325 | 5.15841 | 5.973039 | 40.34922 | 0.598368 | 0.000446 | 0.178947 | 1.514003 | upregulated |
| COL5A1 | COL5A1 | 12831 | AI413331 | collagen_ t | 9.778199 | 10.0711 | 9.575167 | 10.27669 | 10.30841 | 10.78407 | 1122.323 | 0.666181 | 0.019671 | 0.219101 | 1.586867 | |
| COL1A2 | COL1A2 | 12843 | AA960264\| | collagen_ t | 10.04903 | 10.46901 | 9.873017 | 10.57129 | 10.79906 | 11.03759 | 1414.916 | 0.673022 | 0.020477 | 0.219101 | 1.59441 | |
| ITGA5 | ITGA5 | 16402 | Cd49e\|Fnr | integrin alp | 6.46238 | 6.564914 | 6.123179 | 7.016382 | 7.008384 | 6.780394 | 101.0743 | 0.693244 | 0.034727 | 0.290362 | 1.616915 | |
| S100A4 | S100A4 | 20198 | 18A2\|42a\| | S100 calciu | 8.625515 | 8.433669 | 8.875039 | 9.724919 | 9.411951 | 8.920572 | 511.5073 | 0.764583 | 0.03272 | 0.280081 | 1.698878 | |
| EHMT1 | EHMT1 | 77683 | 9230102N1 | euchromat | 5.06752 | 5.211277 | 4.85469 | 5.099403 | 4.988485 | 5.866124 | 36.2837 | 0.786965 | 0.014135 | 0.196958 | 1.725441 | |
| LOX | LOX | 16948 | AI893619\| | lysyl oxidas | 9.976191 | 10.52382 | 10.06059 | 11.31934 | 11.30941 | 11.02367 | 1665.997 | 1.029014 | 0.003289 | 0.178947 | 2.040629 | |
| PHYKPL | PHYKPL | 72947 | 2900006B1 | 5-phospho | 5.56056 | 5.041352 | 5.439652 | 5.645891 | 5.638739 | 6.413612 | 49.29267 | 1.057601 | 0.042429 | 0.307794 | 2.081467 | |
| ACTA2 | ACTA2 | 11475 | 0610041G0 | actin_ alph | 5.598035 | 5.882654 | 6.009968 | 6.834469 | 6.81856 | 6.413612 | 76.61472 | 1.164572 | 0.020465 | 0.219101 | 2.241667 | |
| IL1R1 | IL1R1 | 16177 | CD121a\|CD | interleukin | 5.634561 | 5.211277 | 5.202613 | 5.949507 | 6.184882 | 6.165684 | 52.88381 | 1.178803 | 0.015272 | 0.196958 | 2.263889 | |
| COL1A1 | COL1A1 | 12842 | Col1a-1\|Co | collagen_ t | 9.413295 | 9.899333 | 8.957501 | 10.45417 | 10.54988 | 11.00173 | 1057.166 | 1.237459 | 0.009085 | 0.178947 | 2.357829 | |
| FABP4 | FABP4 | 11770 | 422/aP2\|A | fatty acid b | 5.927342 | 5.813942 | 5.439652 | 6.243793 | 7.094684 | 6.920572 | 75.5834 | 1.411725 | 0.008269 | 0.178947 | 2.66055 | |
| LPL | LPL | 16956 | - | lipoprotein | 5.441915 | 5.813942 | 5.095698 | 6.23518 | 6.219439 | 6.689246 | 60.37601 | 1.421676 | 0.011289 | 0.178947 | 2.678965 | |
| FST | FST | 14313 | AL033346\| | follistatin | 5.013072 | 4.778318 | 4.565183 | 5.451388 | 5.357719 | 5.809541 | 35.81611 | 1.61471 | 0.007006 | 0.178947 | 3.0625 | |
| SERPINE1 | SERPINE1 | 18787 | PAI-1\|PAI1 | serine (or c | 6.836194 | 7.564914 | 7.614427 | 9.8223 | 9.637142 | 7.750647 | 294.9386 | 2.067774 | 0.022636 | 0.230675 | 4.192394 | |
| C3AR1 | C3AR1 | 12267 | AZ3B\|C3AF | complemer | 4.400095 | 4.626315 | 4.643186 | 5.436438 | 4.43327 | 6.558002 | 32.36175 | 2.610053 | 0.010929 | 0.178947 | 6.105263 | |
| TNN | TNN | 329278 | Tnw\|tenas | tenascin N | 4.219523 | 4.626315 | 3.717186 | 5.260866 | 5.998469 | 6.375138 | 32.7385 | 3.979822 | 0.001842 | 0.178947 | 15.77778 | |

WT bone marrow derived macrophages - MDP treated vs control
Nanostring Fibrosis Panel

**Downregulated**

| upperSymb | Symbol | GeneID | Alias | Descriptior | 20210614_ | 20210614_ | 20210614_ | 20210302_ | 20210302_ | 20210302_ | baseMean | log2FoldCh | pvalue | padj | log_10_pa | FC |
|---|---|---|---|---|---|---|---|---|---|---|---|---|---|---|---|---|
| ANGPTL4 | ANGPTL4 | 57875 | Arp4\|Bk89 | angiopoiet | 7.647892 | 8.153658 | 7.920556 | 4.48785 | 5.504988 | 4.899021 | 86.56193 | -3.77327 | 0.001148 | 0.003966 | 2.401642 | 13.67308 |
| CXCR3 | CXCR3 | 12766 | Cd183\|Cm | chemokine | 5.17396 | 4.83955 | 4.809048 | 4.400387 | 3.541514 | 3.651093 | 21.15009 | -3.75582 | 0.013761 | 0.025711 | 1.589874 | 13.5087 |
| CD244A | CD244A | NA | NA | NA | 4.682107 | 4.759379 | 4.971319 | 3.207742 | 4.126476 | 3.821018 | 19.17747 | -3.70215 | 0.008864 | 0.018154 | 1.741037 | 13.01541 |
| COL5A3 | COL5A3 | 53867 | - | collagen_ t | 4.074425 | 3.952024 | 3.809048 | 3.985349 | 3.541514 | 3.236056 | 13.60818 | -3.26833 | 1.48E-05 | 0.000282 | 3.549281 | 9.63532 |
| KLF5 | KLF5 | 12224 | 4930520J0 | Kruppel-lik | 4.004035 | 3.878024 | 3.664658 | 3.722315 | 2.804548 | 3.651093 | 12.30164 | -3.10856 | 5.12E-05 | 0.000565 | 3.247658 | 8.625206 |
| BCL2 | BCL2 | 12043 | AW986256 | B cell leuke | 4.852032 | 4.759379 | 4.370926 | 3.985349 | 4.026941 | 4.557984 | 21.48764 | -2.86082 | 0.018653 | 0.032991 | 1.481609 | 7.264303 |
| PPARD | PPARD | 19015 | NUC-1\|NU | peroxisom | 5.26707 | 5.384984 | 5.001693 | 4.307278 | 4.307049 | 4.458448 | 27.62215 | -2.64514 | 0.001278 | 0.004235 | 2.37318 | 6.255562 |
| EEF2K | EEF2K | 13631 | C86191\|eE | eukaryotic | 5.948894 | 6.089528 | 6.016643 | 4.648314 | 5.026941 | 4.821018 | 42.96896 | -2.22576 | 0.002729 | 0.007113 | 2.147937 | 4.67758 |
| CEBPA | CEBPA | 12606 | C/ebpalpha | CCAAT/enh | 9.544745 | 9.77348 | 9.632263 | 7.295205 | 7.695319 | 7.738556 | 391.6066 | -2.18466 | 0.000191 | 0.001172 | 2.931171 | 4.546192 |
| PLCG1 | PLCG1 | 18803 | AI894140\| | phospholip | 6.141539 | 6.121949 | 5.826121 | 4.859819 | 4.611903 | 5.043411 | 43.23488 | -2.14206 | 0.000114 | 0.000868 | 3.061533 | 4.413927 |
| CXCR4 | CXCR4 | 12767 | CD184\|Cm | chemokine | 10.12194 | 10.22535 | 10.2152 | 8.015097 | 8.476974 | 8.174655 | 590.1222 | -2.03551 | 0.000172 | 0.001115 | 2.952616 | 4.099681 |
| PELI2 | PELI2 | 93834 | AW047589 | pellino 2 | 6.021957 | 6.169255 | 6.287095 | 4.722315 | 5.26398 | 5.110525 | 48.36376 | -1.96074 | 0.004884 | 0.011291 | 1.947269 | 3.892617 |
| LPAR5 | LPAR5 | 381810 | GPR93\|Gm | lysophosph | 7.910926 | 7.99204 | 7.940292 | 6.128307 | 6.804548 | 6.483983 | 148.0578 | -1.72449 | 0.002212 | 0.006039 | 2.219002 | 3.304622 |
| LRP6 | LRP6 | 16974 | C030016K | low density | 7.71553 | 8.009475 | 8.134407 | 6.550134 | 6.877797 | 6.323519 | 154.1805 | -1.57121 | 0.001252 | 0.004175 | 2.379333 | 2.971545 |
| ARRB1 | ARRB1 | 109689 | 1200006I1 | arrestin_ b | 8.153783 | 8.176991 | 8.24962 | 6.629206 | 7.000946 | 6.860547 | 182.5121 | -1.54545 | 0.000337 | 0.001761 | 2.754354 | 2.918946 |
| KIF3A | KIF3A | 16568 | Kif3\|Kifl\|K | kinesin fam | 7.181953 | 7.192339 | 7.038782 | 5.648314 | 6.348869 | 5.821018 | 92.9605 | -1.51559 | 0.002418 | 0.006497 | 2.187285 | 2.859155 |
| RORA | RORA | 19883 | 9530021D1 | RAR-relate | 6.726501 | 6.685379 | 7.067782 | 5.48785 | 5.804548 | 5.936496 | 77.96525 | -1.49561 | 0.00229 | 0.006185 | 2.208629 | 2.819828 |
| CD4 | CD4 | 12504 | L3T4\|Ly-4 | CD4 antige | 4.489462 | 4.273952 | 4.171618 | 4.307278 | 4.026941 | 4.651093 | 19.97408 | -1.49303 | 0.005574 | 0.012422 | 1.905798 | 2.8148 |
| ACVRL1 | ACVRL1 | 11482 | AI115505\| | activin A re | 5.704475 | 5.878024 | 5.299373 | 4.792704 | 4.863442 | 5.236056 | 39.2788 | -1.4461 | 0.004788 | 0.011147 | 1.952843 | 2.7247 |
| GNG2 | GNG2 | 14702 | - | guanine nu | 8.165923 | 8.218811 | 8.240097 | 6.84333 | 7.230813 | 7.026133 | 196.8361 | -1.32847 | 0.000633 | 0.002724 | 2.564773 | 2.511356 |
| RPS6KA2 | RPS6KA2 | 20112 | 90kDa\|D1 | ribosomal | 5.910926 | 5.584293 | 5.702132 | 5.207742 | 4.804548 | 5.29495 | 42.73754 | -1.30171 | 0.002691 | 0.007113 | 2.147937 | 2.465201 |
| PRKAB2 | PRKAB2 | 108097 | 5730553K2 | protein kin | 5.96751 | 5.630096 | 5.461124 | 4.792704 | 5.077567 | 5.405981 | 41.90829 | -1.28757 | 0.016232 | 0.029659 | 1.527838 | 2.441159 |
| MMP8 | MMP8 | 17394 | BB138268 | matrix met | 4.930035 | 5.244806 | 4.809048 | 4.722315 | 4.611903 | 4.821018 | 28.97066 | -1.24066 | 0.001923 | 0.005557 | 2.255136 | 2.363063 |
| TBC1D4 | TBC1D4 | 210789 | 5930406J0 | TBC1 doma | 5.205669 | 5.273952 | 5.299373 | 5.100827 | 4.389511 | 5.110525 | 33.43552 | -1.23614 | 0.027847 | 0.045081 | 1.346007 | 2.355667 |
| GAB1 | GAB1 | 14388 | AA408973 | growth fac | 6.682107 | 6.769648 | 6.616402 | 5.52967 | 6.173782 | 5.899021 | 77.62441 | -1.19548 | 0.011847 | 0.022646 | 1.645004 | 2.290217 |
| PDE3B | PDE3B | 18576 | 9830102A( | phosphodi | 7.091498 | 7.137891 | 7.184914 | 6.444781 | 6.077567 | 6.142947 | 102.5322 | -1.19037 | 0.001295 | 0.004263 | 2.370323 | 2.28211 |
| SMAD3 | SMAD3 | 17127 | AU022421 | SMAD fami | 6.821659 | 6.759379 | 6.720511 | 6.015097 | 6.052476 | 5.821018 | 82.42576 | -1.14218 | 0.001801 | 0.005393 | 2.268184 | 2.207147 |
| MAP3K1 | MAP3K1 | 26401 | MAPKKK1\| | mitogen-ac | 8.75357 | 8.86611 | 8.884336 | 7.740237 | 7.870638 | 7.727909 | 316.7352 | -1.1417 | 9.33E-05 | 0.0008 | 3.097117 | 2.206402 |
| EPAS1 | EPAS1 | 13819 | HIF-2alpha | endothelia | 6.221266 | 6.330536 | 6.158197 | 5.044243 | 5.348869 | 6.142947 | 58.66154 | -1.09265 | 0.028533 | 0.046042 | 1.336845 | 2.132653 |
| SKP2 | SKP2 | 27401 | 4930500A( | S-phase kir | 6.141539 | 6.584293 | 6.184914 | 5.60984 | 5.577138 | 5.780376 | 63.10504 | -1.05616 | 0.002156 | 0.005921 | 2.227609 | 2.079383 |
| CSNK1E | CSNK1E | 27373 | AI426939\| | casein kina | 6.588998 | 6.800021 | 6.800434 | 5.685789 | 6.196866 | 6.142947 | 82.66335 | -1.04037 | 0.0104 | 0.020191 | 1.694845 | 2.056758 |
| CKAP4 | CKAP4 | 216197 | 5630400A( | cytoskeleto | 7.389466 | 7.337455 | 7.21763 | 6.422755 | 6.611903 | 6.458448 | 119.9489 | -1.03468 | 0.000658 | 0.002724 | 2.564773 | 2.048656 |
| NFAM1 | NFAM1 | 74039 | 4921501M | Nfat activa | 9.185933 | 8.981035 | 9.085615 | 7.900233 | 8.253009 | 8.412645 | 397.9413 | -0.95795 | 0.004686 | 0.010985 | 1.9592 | 1.942548 |
| ATG2B | ATG2B | 76559 | 2410024A | autophagy | 6.382547 | 6.512739 | 6.224085 | 5.757939 | 5.863442 | 5.780376 | 67.97135 | -0.93649 | 0.001913 | 0.005557 | 2.255136 | 1.913873 |
| ANAPC1 | ANAPC1 | 17222 | 2610021O( | anaphase p | 6.930035 | 6.849265 | 6.948112 | 6.400387 | 5.863442 | 6.432453 | 95.05006 | -0.93134 | 0.009577 | 0.019193 | 1.716863 | 1.907042 |
| PPARG | PPARG | 19016 | Nr1c3\|PPA | peroxisom | 6.891561 | 6.858915 | 6.783052 | 5.89224 | 6.369333 | 6.29495 | 91.45616 | -0.93127 | 0.008843 | 0.018154 | 1.741037 | 1.906949 |
| NCOR2 | NCOR2 | 20602 | N-CoR\|SM | nuclear rec | 8.108372 | 8.035238 | 8.194807 | 7.331124 | 7.585908 | 7.060484 | 210.7403 | -0.90716 | 0.009116 | 0.018568 | 1.731244 | 1.875346 |
| ANAPC7 | ANAPC7 | 56317 | APC7\|AW5 | anaphase p | 6.074425 | 6.022414 | 6.144651 | 5.354583 | 5.679018 | 5.780376 | 57.38404 | -0.89607 | 0.008044 | 0.01694 | 1.771097 | 1.860992 |
| SCD2 | SCD2 | 20250 | Mir5114\|S | stearoyl-Co | 10.91812 | 10.75486 | 10.81601 | 9.786249 | 10.15622 | 9.94801 | 1347.976 | -0.88413 | 0.001794 | 0.005393 | 2.268184 | 1.845653 |
| DEPDC5 | DEPDC5 | 277854 | AV016528 | DEP domai | 7.013024 | 6.652464 | 6.826121 | 6.181747 | 6.173782 | 6.265803 | 91.69833 | -0.88146 | 0.001484 | 0.004639 | 2.3336 | 1.842238 |
| KRAS | KRAS | 16653 | AI929937\| | v-Ki-ras2 Ki | 7.057147 | 6.933877 | 6.940292 | 6.258368 | 6.369333 | 6.379014 | 100.869 | -0.87736 | 0.000228 | 0.001366 | 2.864682 | 1.837012 |
| FNIP2 | FNIP2 | 329679 | D630023B | folliculin in | 9.57844 | 9.600202 | 9.668213 | 8.904213 | 8.892011 | 8.695488 | 597.624 | -0.82434 | 0.000506 | 0.002396 | 2.620577 | 1.770722 |
| ERN1 | ERN1 | 78943 | 9030414B1 | endoplasm | 7.570851 | 7.824853 | 7.734143 | 7.072812 | 7.089951 | 6.936496 | 165.5953 | -0.81793 | 0.001588 | 0.004902 | 2.309618 | 1.762871 |
| ARHGEF6 | ARHGEF6 | 73341 | 1600028C( | Rac/Cdc42 | 8.780139 | 8.774755 | 8.806899 | 7.947281 | 8.046134 | 8.134909 | 341.3291 | -0.81146 | 0.000111 | 0.000868 | 3.061533 | 1.75499 |
| PARP1 | PARP1 | 11545 | 5830444G | poly (ADP-r | 8.621687 | 8.66627 | 8.659904 | 7.835015 | 7.899066 | 8.017416 | 311.5298 | -0.80515 | 9.92E-05 | 0.000821 | 3.085399 | 1.74733 |
| ATG101 | ATG101 | 68118 | 9430023L2 | autophagy | 7.082987 | 7.056361 | 6.826121 | 6.331124 | 6.611903 | 6.29495 | 104.0097 | -0.80174 | 0.009958 | 0.019638 | 1.706892 | 1.743207 |
| ALDH9A1 | ALDH9A1 | 56752 | AA139417 | aldehyde d | 9.555551 | 9.603076 | 9.549872 | 8.771074 | 8.97113 | 8.711789 | 585.5907 | -0.791 | 0.000842 | 0.00322 | 2.492195 | 1.730278 |
| HMOX1 | HMOX1 | 15368 | D8Wsu38e | heme oxyg | 10.42694 | 10.48185 | 10.46385 | 9.612275 | 9.81391 | 9.648273 | 1078.28 | -0.7873 | 0.000303 | 0.001657 | 2.780743 | 1.725847 |
| VEGFB | VEGFB | 22340 | VEGF-B\|Vr | vascular en | 7.004035 | 6.849265 | 7.074942 | 6.331124 | 6.679018 | 6.29495 | 104.3694 | -0.76294 | 0.0162 | 0.029659 | 1.527838 | 1.69694 |
| SKI | SKI | 108077 | 4930534J0 | superkiller | 8.213489 | 8.298981 | 8.016643 | 7.128307 | 7.662529 | 7.695488 | 228.4772 | -0.75912 | 0.009881 | 0.019565 | 1.708515 | 1.69246 |
| TBL1XR1 | TBL1XR1 | 81004 | 8030499H( | transducin | 8.416817 | 8.398281 | 8.399724 | 7.638792 | 7.76648 | 7.810965 | 269.071 | -0.74852 | 0.000144 | 0.001013 | 2.994438 | 1.680068 |
| ARG1 | ARG1 | 11846 | AI\|AI2565 | arginase_ l | 5.871932 | 6.005136 | 5.971319 | 5.60984 | 5.804548 | 5.458448 | 55.21049 | -0.74282 | 0.029728 | 0.047508 | 1.323236 | 1.673443 |
| APC | APC | 11789 | AI047805\| | adenomato | 8.19386 | 8.056361 | 8.074942 | 7.354583 | 7.550503 | 7.496583 | 220.9851 | -0.7416 | 0.000791 | 0.003048 | 2.515952 | 1.672027 |
| PRKAG1 | PRKAG1 | 19082 | AA571379 | protein kin | 7.091498 | 7.337455 | 6.93243 | 6.466476 | 6.645851 | 6.60529 | 115.0805 | -0.7408 | 0.006335 | 0.013871 | 1.857892 | 1.671104 |
| CDKN2C | CDKN2C | 12580 | C77269\|IN | cyclin-depe | 9.238613 | 9.220683 | 9.266917 | 8.560258 | 8.577138 | 8.521458 | 476.8895 | -0.73549 | 1.23E-05 | 0.000277 | 3.557665 | 1.664962 |
| CASP6 | CASP6 | 12368 | CASP-6\|M | caspase 6 | 8.368608 | 8.450275 | 8.305473 | 7.477203 | 7.920026 | 7.840918 | 266.9484 | -0.71134 | 0.009782 | 0.019521 | 1.709488 | 1.637329 |
| FLI1 | FLI1 | 14247 | EWSR2\|Fli- | Friend leuk | 9.44532 | 9.467724 | 9.43493 | 8.624389 | 8.940685 | 8.78552 | 555.0323 | -0.70531 | 0.001906 | 0.005557 | 2.255136 | 1.6305 |
| LATS2 | LATS2 | 50523 | 4932411G( | large tumo | 7.758923 | 7.759379 | 7.715938 | 7.194803 | 7.219586 | 7.094037 | 175.7171 | -0.69955 | 0.000403 | 0.001981 | 2.703155 | 1.623995 |
| GCNT1 | GCNT1 | 14537 | 5630400D | glucosamin | 8.777504 | 8.652464 | 8.702132 | 8.015097 | 8.312343 | 7.945714 | 337.9991 | -0.68488 | 0.006703 | 0.014484 | 1.839117 | 1.607566 |
| CCNA2 | CCNA2 | 12428 | AA408589 | cyclin A2 | 9.60692 | 9.574076 | 9.536968 | 8.792704 | 9.004221 | 8.973022 | 608.0243 | -0.68467 | 0.000656 | 0.002724 | 2.564773 | 1.607333 |
| HADH | HADH | 15107 | AA409008 | hydroxyacy | 7.944202 | 7.978824 | 7.876162 | 7.194803 | 7.476974 | 7.419278 | 200.6273 | -0.67644 | 0.002636 | 0.007007 | 2.154462 | 1.598186 |
| PCCB | PCCB | 66904 | 1300012P( | propionyl C | 7.630475 | 7.554909 | 7.566477 | 7.015097 | 7.138449 | 7.008645 | 159.6766 | -0.66383 | 0.000714 | 0.002862 | 2.543403 | 1.584282 |
| PRKACB | PRKACB | 18749 | CbPKA\|Pka | protein kin | 8.831854 | 8.814972 | 8.720511 | 8.264573 | 8.32811 | 7.98201 | 359.6222 | -0.65807 | 0.006555 | 0.014227 | 1.846874 | 1.577975 |
| LDLRAP1 | LDLRAP1 | 100017 | AA691260 | low density | 9.572372 | 9.512739 | 9.486851 | 8.744683 | 8.984457 | 8.977523 | 593.5 | -0.65739 | 0.001507 | 0.00468 | 2.329743 | 1.577224 |
| EHMT1 | EHMT1 | 77683 | 9230102N | euchromat | 6.368608 | 6.411456 | 6.184914 | 6.207742 | 5.774175 | 5.936496 | 70.87632 | -0.64665 | 0.02068 | 0.035806 | 1.446043 | 1.565526 |
| INPP5D | INPP5D | 16331 | SHIP\|SHIP- | inositol pol | 7.642109 | 7.733384 | 7.967477 | 7.141854 | 7.18537 | 7.445509 | 183.4552 | -0.63561 | 0.009204 | 0.018595 | 1.730598 | 1.553595 |
| MKI67 | MKI67 | 17345 | D630048A | antigen ide | 10.31055 | 10.41555 | 10.4047 | 9.634007 | 9.90258 | 9.754381 | 1075.13 | -0.63284 | 0.001887 | 0.005557 | 2.255136 | 1.550615 |
| MAF | MAF | 17132 | 2810401A | avian musc | 9.050614 | 9.026701 | 9.005445 | 8.371931 | 8.590273 | 8.399286 | 427.775 | -0.62278 | 0.001342 | 0.004358 | 2.360722 | 1.539835 |
| ADIPOR1 | ADIPOR1 | 72674 | 2810031L1 | adiponecti | 8.47327 | 8.462986 | 8.388274 | 7.685789 | 8.167953 | 7.850766 | 288.3159 | -0.61348 | 0.017998 | 0.032293 | 1.490891 | 1.529942 |
| CD84 | CD84 | 12523 | A130013D | CD84 antig | 12.12142 | 12.11894 | 12.09004 | 11.43793 | 11.57108 | 11.50907 | 3585.803 | -0.60985 | 0.000111 | 0.000868 | 3.061533 | 1.526101 |
| SLC25A13 | SLC25A13 | 50799 | AI785475\| | solute carri | 6.862016 | 6.789968 | 6.81761 | 6.377667 | 6.348869 | 6.533736 | 98.47218 | -0.60963 | 0.000175 | 0.001115 | 2.952616 | 1.525871 |
| F8 | F8 | 14069 | Cf-8\|Cf8\|F | coagulatio | 5.108372 | 5.022414 | 5.347467 | 5.100827 | 5.126476 | 5.174655 | 35.42515 | -0.60054 | 0.018977 | 0.033209 | 1.478741 | 1.516289 |
| UBE4B | UBE4B | 63958 | 4930551I1 | ubiquitinat | 7.665101 | 7.938435 | 7.872057 | 7.128307 | 7.550503 | 7.323519 | 191.2948 | -0.60022 | 0.021582 | 0.036647 | 1.435965 | 1.51595 |
| ERP29 | ERP29 | 67397 | 1200015M | endoplasm | 11.65974 | 11.63291 | 11.64043 | 11.00401 | 11.0938 | 11.07 | 2610.571 | -0.59624 | 3.04E-05 | 0.000438 | 3.358206 | 1.511766 |
| PROS1 | PROS1 | 19128 | AW214361 | protein S (a | 7.841979 | 7.764523 | 7.688193 | 7.168571 | 7.274868 | 7.405981 | 184.0583 | -0.59148 | 0.00186 | 0.005534 | 2.256947 | 1.506788 |
| AKR1B8 | AKR1B8 | 14187 | FR-1\|Fgfrp | aldo-keto r | 7.56475 | 7.456645 | 7.482819 | 7.181747 | 7.1621 | 6.800841 | 154.8596 | -0.58956 | 0.024009 | 0.039909 | 1.398934 | 1.504793 |

**Upregulated**

| | | | | | | | | | | | | | | | |
|---|---|---|---|---|---|---|---|---|---|---|---|---|---|---|---|
| TXN1 | TXN1 | 22166 | ADF\|AW55 | thioredoxin | 12.28248 | 12.26332 | 12.2516 | 12.92982 | 12.82321 | 12.85872 | 6072.99 | 0.60352 | 5.02E-05 | 0.000565 | 3.247658 | 1.519419 |
| STAT3 | STAT3 | 20848 | 1110034C( | signal trans | 8.389466 | 8.371563 | 8.501538 | 9.185022 | 8.984457 | 9.017416 | 428.0327 | 0.623924 | 0.001219 | 0.004104 | 2.386776 | 1.541061 |
| CASP8 | CASP8 | 12370 | CASP-8\|FLI | caspase 8 | 9.195835 | 9.376611 | 9.334102 | 10.09387 | 9.802669 | 9.922556 | 787.3936 | 0.628063 | 0.003407 | 0.008466 | 2.072335 | 1.545489 |
| PLCB3 | PLCB3 | 18797 | mKIAA409 | phospholip | 7.403206 | 7.137891 | 7.144651 | 7.784091 | 7.961053 | 7.990943 | 190.0593 | 0.645549 | 0.0048 | 0.011147 | 1.952843 | 1.564334 |
| NCF4 | NCF4 | 17972 | AI451400\| | neutrophil | 10.05823 | 10.00731 | 10.02591 | 10.73242 | 10.74609 | 10.82477 | 1350.353 | 0.732562 | 2.09E-05 | 0.000318 | 3.496915 | 1.661587 |
| TLR6 | TLR6 | 21899 | - | toll-like rec | 7.56475 | 7.774755 | 7.596639 | 8.411614 | 8.572733 | 8.236056 | 260.6719 | 0.735373 | 0.004517 | 0.010794 | 1.966829 | 1.664828 |
| F11R | F11R | 16456 | 9130004G | F11 recept | 6.76957 | 6.685379 | 6.635899 | 7.498419 | 7.40941 | 7.533736 | 136.12 | 0.737279 | 0.000125 | 0.000913 | 3.03975 | 1.667029 |
| MOB1B | MOB1B | 68473 | 1110003E( | MOB kinas | 7.09996 | 7.022414 | 7.046087 | 7.89224 | 7.727381 | 7.870262 | 174.0137 | 0.737605 | 0.000168 | 0.001115 | 2.952616 | 1.667406 |
| MAPK9 | MAPK9 | 26420 | AI851083\| | mitogen-ac | 7.698916 | 7.685379 | 7.540851 | 8.545045 | 8.301735 | 8.385803 | 260.7075 | 0.744926 | 0.00114 | 0.003966 | 2.401642 | 1.675889 |
| GPR65 | GPR65 | 14744 | Dig1\|Gpcr | G-protein c | 9.762925 | 9.777301 | 9.811193 | 10.54122 | 10.4992 | 10.58331 | 1146.105 | 0.752035 | 9.92E-06 | 0.00026 | 3.585766 | 1.684167 |
| PTPRC | PTPRC | 19264 | B220\|CD4 | protein tyr | 10.79845 | 10.74514 | 10.79881 | 11.55331 | 11.53133 | 11.52761 | 2286.792 | 0.753894 | 2.90E-06 | 0.000144 | 3.841277 | 1.686338 |
| PIK3R5 | PIK3R5 | 320207 | AV230647 | phosphoin | 7.618745 | 7.430996 | 7.556281 | 8.28303 | 8.369333 | 8.385803 | 245.6905 | 0.786884 | 0.000303 | 0.001657 | 2.780743 | 1.725344 |
| RELA | RELA | 19697 | p65 | v-rel reticu | 5.985888 | 6.398281 | 6.211146 | 7.181747 | 6.933831 | 7.077358 | 99.13861 | 0.808735 | 0.004989 | 0.011427 | 1.942086 | 1.751675 |
| CHMP4B | CHMP4B | 75608 | 2010012F( | charged m | 9.844499 | 9.639925 | 9.735273 | 10.68 | 10.47697 | 10.54894 | 1139.565 | 0.823915 | 0.00062 | 0.002724 | 2.564773 | 1.770203 |
| PTAFR | PTAFR | 19204 | PAFR | platelet-ac | 10.68773 | 10.67517 | 10.71994 | 11.60006 | 11.41803 | 11.54439 | 2206.517 | 0.824058 | 0.000118 | 0.000875 | 3.057911 | 1.770379 |
| EP400 | EP400 | 75560 | 1700020J0 | E1A bindin | 6.891561 | 6.978824 | 7.21763 | 7.977816 | 7.892011 | 7.790645 | 175.8353 | 0.825473 | 0.002445 | 0.006534 | 2.184832 | 1.772116 |
| H2-M3 | H2-M3 | 14991 | H-2M3\|Hn | histocomp | 7.896426 | 7.701559 | 7.734143 | 8.826652 | 8.384493 | 8.695488 | 295.3865 | 0.840293 | 0.004566 | 0.010807 | 1.966302 | 1.790414 |
| CD14 | CD14 | 12475 | - | CD14 antig | 10.72445 | 10.60952 | 10.67824 | 11.58588 | 11.50325 | 11.49187 | 2193.272 | 0.853862 | 4.59E-05 | 0.000543 | 3.264857 | 1.807333 |
| MAP2K1 | MAP2K1 | 26395 | MAPKK1\|M | mitogen-ac | 9.195835 | 9.151697 | 9.11717 | 10.12149 | 9.989424 | 10.04984 | 778.3318 | 0.892415 | 3.63E-05 | 0.000481 | 3.317694 | 1.856281 |
| TLR1 | TLR1 | 21897 | - | toll-like rec | 9.338532 | 9.052161 | 9.222474 | 10.19643 | 10.06977 | 10.07526 | 808.5002 | 0.903578 | 0.000668 | 0.002745 | 2.561498 | 1.8707 |
| ADAM17 | ADAM17 | 11491 | CD156b\|Ta | a disintegri | 10.89095 | 10.8432 | 10.86278 | 11.79056 | 11.86028 | 11.80211 | 2595.23 | 0.950315 | 3.47E-06 | 0.000157 | 3.804981 | 1.932295 |
| H2-K1 | H2-K1 | 14972 | H-2K\|H-2K | histocomp | 12.52485 | 12.42998 | 12.51789 | 13.57063 | 13.42416 | 13.42794 | 8093.642 | 0.982832 | 6.72E-05 | 0.000669 | 3.174592 | 1.976342 |
| H2-T23 | H2-T23 | 15040 | 37b\|37c\|H | histocomp | 11.0091 | 10.78301 | 10.9984 | 12.03565 | 11.93726 | 11.84892 | 2769.466 | 1.009062 | 0.000392 | 0.001967 | 2.706111 | 2.012602 |
| FCGR4 | FCGR4 | 246256 | 4833442P2 | Fc recepto | 10.59649 | 10.41064 | 10.62983 | 11.78787 | 11.41496 | 11.54212 | 2140.379 | 1.034351 | 0.001331 | 0.004353 | 2.361193 | 2.048192 |
| MMP9 | MMP9 | 17395 | AW743869 | matrix met | 8.766916 | 8.615938 | 8.699818 | 9.511523 | 9.881364 | 9.813485 | 594.2148 | 1.035133 | 0.001138 | 0.003966 | 2.401642 | 2.049302 |
| LYN | LYN | 17096 | AA407514 | Yamaguchi | 10.0712 | 10.05006 | 9.992269 | 11.15944 | 11.05327 | 11.06366 | 1514.875 | 1.052035 | 1.44E-05 | 0.000282 | 3.549281 | 2.073453 |
| H2-AA | H2-AA | 14960 | Aalpha\|H- | histocomp | 7.594997 | 7.323583 | 7.411084 | 8.560258 | 8.527925 | 8.509074 | 253.8478 | 1.076989 | 0.000242 | 0.001416 | 2.848994 | 2.109629 |
| CASP7 | CASP7 | 12369 | AI314680\| | caspase 7 | 8.04843 | 7.829769 | 8.038782 | 9.168571 | 8.96442 | 9.043411 | 365.9655 | 1.078118 | 0.000352 | 0.001804 | 2.743759 | 2.11128 |
| MMP13 | MMP13 | 17386 | Clg\|MMP- | matrix met | 8.51183 | 8.244806 | 8.425159 | 9.720059 | 9.42171 | 9.29495 | 489.9256 | 1.085841 | 0.001134 | 0.003966 | 2.401642 | 2.122613 |
| CCL2 | CCL2 | 20296 | AI323594\| | chemokine | 12.16466 | 11.8553 | 11.91408 | 13.20855 | 13.01988 | 13.08364 | 5959.677 | 1.125569 | 0.000515 | 0.002414 | 2.61735 | 2.181876 |
| NFKB1 | NFKB1 | 18033 | NF-KB1\|NF | nuclear fac | 8.621687 | 8.551937 | 8.579121 | 9.792704 | 9.727381 | 9.690012 | 572.2389 | 1.14805 | 7.61E-06 | 0.000233 | 3.632238 | 2.216141 |
| PSMB8 | PSMB8 | 16913 | Lmp-7\|Lm | proteasom | 10.76093 | 10.54448 | 10.76158 | 11.92051 | 11.80643 | 11.79192 | 2459.771 | 1.149639 | 0.000161 | 0.001112 | 2.953765 | 2.218583 |
| ITGA5 | ITGA5 | 16402 | Cd49e\|Fnr | integrin alp | 7.157841 | 7.222553 | 7.268479 | 8.411614 | 8.301735 | 8.438996 | 222.8923 | 1.157355 | 1.96E-05 | 0.000318 | 3.496915 | 2.230481 |
| IL10RA | IL10RA | 16154 | AW553859 | interleukin | 8.77222 | 8.782381 | 8.902559 | 10.07987 | 10.01563 | 9.887108 | 678.6984 | 1.171642 | 8.89E-05 | 0.0008 | 3.097117 | 2.25268 |
| H2-AB1 | H2-AB1 | 14961 | AI845868\| | histocomp | 6.670792 | 7.005136 | 6.978973 | 8.155274 | 7.974473 | 8.150939 | 179.6773 | 1.197858 | 0.00064 | 0.002724 | 2.564773 | 2.293989 |
| CYBB | CYBB | 13058 | C88302\|C( | cytochrom | 12.3069 | 12.21247 | 12.3408 | 13.53754 | 13.46038 | 13.4903 | 7597.97 | 1.209099 | 1.10E-05 | 0.00026 | 3.585766 | 2.311932 |
| PHLPP1 | PHLPP1 | 98432 | AI836256\| | PH domain | 6.682107 | 6.800021 | 6.851358 | 8.18829 | 7.884922 | 7.990943 | 168.851 | 1.233459 | 0.000307 | 0.001657 | 2.780743 | 2.351301 |
| THBS1 | THBS1 | 21825 | TSP-1\|TSP | thrombosp | 5.81139 | 6.005136 | 5.826121 | 7.31925 | 6.98777 | 7.158888 | 91.65188 | 1.25846 | 0.000434 | 0.002096 | 2.678538 | 2.392402 |
| NRP2 | NRP2 | 18187 | 1110048P( | neuropilin | 10.3069 | 10.10988 | 10.24089 | 11.55521 | 11.39326 | 11.5176 | 1850.83 | 1.268796 | 7.56E-05 | 0.000709 | 3.149478 | 2.409603 |
| BST2 | BST2 | 69550 | 2310015I1 | bone marr | 11.08778 | 10.73927 | 11.06913 | 12.3564 | 12.14403 | 12.25147 | 3121.477 | 1.284916 | 0.000574 | 0.002568 | 2.590373 | 2.436678 |
| HC | HC | 15139 | C5\|C5a\|He | hemolytic | 6.410028 | 6.630096 | 6.171618 | 7.60984 | 7.419258 | 8.051973 | 132.4039 | 1.29389 | 0.003214 | 0.008067 | 2.093305 | 2.451883 |
| TRAF2 | TRAF2 | 22030 | AI325259 | TNF recept | 7.318732 | 7.309576 | 7.24962 | 8.690406 | 8.532469 | 8.678999 | 249.5699 | 1.338757 | 1.44E-05 | 0.000282 | 3.549281 | 2.529333 |
| AXL | AXL | 26362 | AI323647\| | AXL recept | 8.795849 | 8.706912 | 8.772081 | 10.23959 | 9.986115 | 10.08364 | 690.117 | 1.343932 | 7.07E-05 | 0.000676 | 3.170103 | 2.538421 |
| LOX | LOX | 16948 | AI893619\| | lysyl oxida | 4.76957 | 4.584293 | 4.224085 | 6.072812 | 5.711439 | 6.008645 | 37.49104 | 1.412348 | 0.004374 | 0.010552 | 1.976657 | 2.661701 |
| HCK | HCK | 15162 | AI849071\| | hemopoiet | 9.24244 | 8.978824 | 9.139538 | 10.61834 | 10.52337 | 10.58183 | 921.2128 | 1.454796 | 6.14E-05 | 0.000651 | 3.186368 | 2.741179 |
| ITGA4 | ITGA4 | 16401 | CD49D\|Itg | integrin alp | 10.06473 | 10.07408 | 10.11543 | 11.60006 | 11.5353 | 11.49972 | 1801.38 | 1.460503 | 1.71E-06 | 9.47E-05 | 4.023698 | 2.752043 |
| GNB4 | GNB4 | 14696 | 6720453A | guanine nu | 5.074425 | 4.584293 | 4.370926 | 6.307278 | 5.834296 | 6.236056 | 42.25975 | 1.4746 | 0.009199 | 0.018595 | 1.730598 | 2.779066 |
| OAS1A/G | OAS1A/G | NA | NA | NA | 6.647892 | 6.153658 | 6.236909 | 7.89224 | 7.750967 | 7.821018 | 135.6533 | 1.481972 | 0.000999 | 0.0036 | 2.443721 | 2.793304 |
| CXCL16 | CXCL16 | 66102 | 0910001K | chemokine | 10.37907 | 10.33573 | 10.3789 | 11.87764 | 11.84985 | 11.83845 | 2210.169 | 1.491071 | 1.58E-07 | 3.92E-05 | 4.406889 | 2.810976 |
| NR1H3 | NR1H3 | 22259 | AU018371 | nuclear rec | 7.443659 | 7.411456 | 7.455649 | 9.047845 | 9.042953 | 9.085722 | 303.9901 | 1.628642 | 9.24E-08 | 3.92E-05 | 4.406889 | 3.092219 |
| RELB | RELB | 19698 | shep | avian retic | 8.052795 | 8.009475 | 7.978973 | 9.617133 | 9.666669 | 9.690012 | 456.9342 | 1.64906 | 7.00E-07 | 7.58E-05 | 4.120548 | 3.136292 |
| FLNB | FLNB | 286940 | AL024016\| | filamin_ be | 6.236696 | 6.199952 | 6.335593 | 8.155274 | 7.719432 | 8.017416 | 138.2109 | 1.728565 | 0.000237 | 0.001401 | 2.853519 | 3.313981 |
| IL21R | IL21R | 60504 | NILR | interleukin | 7.495888 | 7.030976 | 7.24962 | 9.058598 | 8.9441 | 9.013037 | 280.5349 | 1.758933 | 0.000267 | 0.001527 | 2.816021 | 3.384477 |
| MYD88 | MYD88 | 17874 | - | myeloid dif | 8.88178 | 8.720209 | 8.740911 | 10.71554 | 10.5096 | 10.59067 | 827.7887 | 1.829092 | 2.10E-05 | 0.000318 | 3.496915 | 3.553134 |
| CDKN1A | CDKN1A | 12575 | CAP20\|CDI | cyclin-depe | 9.760258 | 9.653851 | 9.753237 | 11.59206 | 11.58043 | 11.58109 | 1610.719 | 1.864735 | 7.62E-07 | 7.58E-05 | 4.120548 | 3.642009 |
| AIM2 | AIM2 | 383619 | Gm1313\|If | absent in n | 7.4303 | 7.281148 | 7.674118 | 9.539938 | 9.202579 | 9.28045 | 338.1272 | 1.89285 | 0.000282 | 0.001586 | 2.799646 | 3.71368 |
| XAF1 | XAF1 | 327959 | Fbox39 | XIAP associ | 8.667949 | 8.259453 | 8.540851 | 10.48652 | 10.30572 | 10.37048 | 693.8581 | 1.905158 | 0.000145 | 0.001013 | 2.994438 | 3.745499 |
| CFLAR | CFLAR | 12633 | 2310024N | CASP8 and | 9.453597 | 9.342623 | 9.492209 | 11.4441 | 11.23918 | 11.37048 | 1342.182 | 1.925429 | 1.38E-05 | 0.000282 | 3.549281 | 3.798499 |
| CD86 | CD86 | 12524 | B7\|B7-2\|B | CD86 antig | 5.852032 | 5.411456 | 5.75658 | 7.775426 | 7.338527 | 7.616878 | 98.71175 | 1.959074 | 0.000689 | 0.002792 | 2.554053 | 3.888123 |
| SRC | SRC | 20779 | AW259666 | Rous sarco | 5.726501 | 6.169255 | 6.299373 | 8.141854 | 7.834296 | 7.999822 | 130.5553 | 1.968415 | 0.000691 | 0.002792 | 2.554053 | 3.913379 |
| CCL4 | CCL4 | 20303 | AT744.1\|A | chemokine | 9.366856 | 8.965487 | 9.115435 | 11.33776 | 11.03497 | 11.03696 | 1130.629 | 1.98934 | 9.91E-05 | 0.000821 | 3.085399 | 3.970554 |
| IFI35 | IFI35 | 70110 | 2010008K | interferon- | 9.177962 | 8.814972 | 9.194807 | 11.11892 | 10.96946 | 11.08259 | 1067.326 | 1.999849 | 0.000116 | 0.000875 | 3.058156 | 3.99958 |
| MET | MET | 17295 | AI838057\| | met proto- | 4.489462 | 3.630096 | 3.809048 | 5.757939 | 5.429039 | 6.323519 | 29.99224 | 2.090198 | 0.008362 | 0.017317 | 1.761525 | 4.258065 |
| PELI1 | PELI1 | 67245 | 2810468L( | pellino 1 | 8.232854 | 8.207525 | 8.326623 | 10.52838 | 10.21818 | 10.26764 | 628.9786 | 2.092567 | 3.63E-05 | 0.000481 | 3.317694 | 4.265064 |
| BIRC3 | BIRC3 | 11796 | AW10767( | baculoviral | 7.396353 | 7.536987 | 7.359245 | 9.697303 | 9.726582 | 9.593608 | 370.7618 | 2.227646 | 3.91E-06 | 0.000162 | 3.790387 | 4.683692 |
| TLR2 | TLR2 | 24088 | Ly105 | toll-like rec | 7.423574 | 7.4693 | 7.347467 | 9.66013 | 9.5661 | 9.642615 | 366.6343 | 2.230664 | 9.73E-07 | 8.06E-05 | 4.093475 | 4.6935 |
| STAT1 | STAT1 | 20846 | 2010005J0 | signal tran | 10.2783 | 9.724174 | 10.18077 | 12.4417 | 12.2561 | 12.2945 | 2345.898 | 2.255872 | 9.14E-05 | 0.0008 | 3.097117 | 4.77623 |
| FLT1 | FLT1 | 14254 | AI323757\| | FMS-like ty | 5.004035 | 4.987648 | 4.940292 | 7.181747 | 6.974473 | 7.205682 | 66.21011 | 2.266705 | 5.58E-06 | 0.000185 | 3.733045 | 4.812228 |
| OAS1B | OAS1B | 23961 | Flv\|L1\|Mn | 2'-5' oligoa | 3.489462 | 4.022414 | 4.370926 | 5.792704 | 5.920026 | 6.142947 | 31.04767 | 2.282662 | 0.00523 | 0.011785 | 1.928662 | 4.865748 |
| JAK2 | JAK2 | 16452 | Fd17 | Janus kinas | 7.687732 | 7.769648 | 7.955889 | 10.34729 | 10.04454 | 10.0196 | 501.7359 | 2.350441 | 6.31E-05 | 0.000653 | 3.185161 | 5.0998 |
| P2RY13 | P2RY13 | 74191 | 2010001L( | purinergic | 3.588998 | 3.630096 | 4.274711 | 6.422755 | 5.804548 | 5.738556 | 30.06356 | 2.464281 | 0.004465 | 0.010721 | 1.969752 | 5.518519 |
| MEFV | MEFV | 54483 | FMF\|TRIM | Mediterrar | 3.141539 | 3.717559 | 3.504193 | 5.60984 | 5.307049 | 5.458448 | 21.9544 | 2.474749 | 0.001747 | 0.005296 | 2.276086 | 5.558706 |
| DAXX | DAXX | 13163 | - | Fas death | 7.149713 | 6.819921 | 6.971319 | 9.539938 | 9.431474 | 9.425881 | 298.801 | 2.521596 | 2.11E-05 | 0.000318 | 3.496915 | 5.742169 |
| CASP4 | CASP4 | 12363 | CASP-11\|C | caspase 4_ | 8.615798 | 8.384984 | 8.509487 | 11.16443 | 10.90521 | 11.00534 | 869.6003 | 2.534072 | 1.57E-05 | 0.000289 | 3.53892 | 5.792041 |
| RAPGEF2 | RAPGEF2 | 76089 | 5830453M | Rap guanin | 7.682107 | 7.207525 | 7.556281 | 10.18011 | 10.00095 | 9.966243 | 435.2 | 2.593993 | 9.28E-05 | 0.0008 | 3.097117 | 6.037675 |
| FAS | FAS | 14102 | AI196731\| | Fas (TNF re | 4.540088 | 4.630096 | 4.224085 | 7.114632 | 6.523367 | 7.110525 | 51.64174 | 2.691534 | 0.000641 | 0.002724 | 2.564773 | 6.46 |
| MARCO | MARCO | 17167 | AI323439\| | macrophag | 5.588998 | 5.67449 | 5.859674 | 8.301254 | 8.374404 | 8.265803 | 128.9592 | 2.702901 | 1.05E-05 | 0.00026 | 3.585766 | 6.511097 |
| IRF1 | IRF1 | 16362 | AU020929 | interferon | 9.311464 | 9.033108 | 9.24962 | 12.3418 | 12.11061 | 12.19316 | 1671.152 | 3.026756 | 1.01E-05 | 0.00026 | 3.585766 | 8.149749 |

| | | | | | | | | | | | | | | | |
|---|---|---|---|---|---|---|---|---|---|---|---|---|---|---|---|
| HCAR2 | HCAR2 | 80885 | Gpr109a\|G | hydroxycar | 4.540088 | 4.089528 | 4.41673 | 7.0003 | 6.98777 | 7.190252 | 52.13251 | 3.027183 | 0.000172 | 0.001115 | 2.952616 | 8.152162 |
| C3 | C3 | 12266 | AI255234\| | complemen | 8.913328 | 8.741232 | 9.055166 | 12.28989 | 11.98736 | 11.99483 | 1445.114 | 3.199879 | 1.95E-05 | 0.000318 | 3.496915 | 9.188818 |
| NLRP3 | NLRP3 | 216799 | AGTAVPRL | NLR family | 6.057147 | 6.005136 | 6.089155 | 9.331124 | 8.906086 | 9.298552 | 195.976 | 3.237401 | 3.89E-05 | 0.000495 | 3.305101 | 9.430939 |
| MMP14 | MMP14 | 17387 | AI325305\| | matrix met | 4.325963 | 4.952024 | 5.11717 | 7.947281 | 7.789441 | 7.695488 | 79.04276 | 3.252776 | 0.000475 | 0.002268 | 2.644369 | 9.531981 |
| KDR | KDR | 16542 | 6130401C0 | kinase inse | 5.004035 | 4.987648 | 5.323621 | 8.482536 | 8.046134 | 8.051973 | 100.3798 | 3.274733 | 0.000113 | 0.000868 | 3.061533 | 9.678161 |
| ICAM1 | ICAM1 | 15894 | CD54\|Icam | intercellula | 7.934773 | 7.978824 | 7.892464 | 11.32816 | 11.19113 | 11.14994 | 764.9472 | 3.312597 | 6.38E-07 | 7.58E-05 | 4.120548 | 9.93553 |
| FGL2 | FGL2 | 14190 | AI385601\| | fibrinogen- | 6.450292 | 5.83955 | 6.738658 | 9.979703 | 9.438754 | 9.602378 | 257.4632 | 3.379341 | 0.000178 | 0.001115 | 2.952616 | 10.40598 |
| ISG15 | ISG15 | 1E+08 | G1p2\|IGI15 | ISG15 ubiq | 10.04186 | 9.304289 | 9.963625 | 13.47352 | 13.14236 | 13.28386 | 2967.266 | 3.507581 | 6.16E-05 | 0.000651 | 3.186368 | 11.37331 |
| ISG20 | ISG20 | 57444 | 1600023I0 | interferon- | 7.495888 | 7.056361 | 7.601605 | 11.28532 | 10.76937 | 10.92954 | 583.9421 | 3.645853 | 4.18E-05 | 0.000507 | 3.294812 | 12.51731 |
| TNF | TNF | 21926 | DIF\|TNF-a\| | tumor necr | 7.642109 | 7.371563 | 7.692855 | 11.32371 | 11.2224 | 11.2219 | 681.4308 | 3.722822 | 4.59E-06 | 0.000169 | 3.772106 | 13.20326 |
| KLRK1 | KLRK1 | 27007 | D6H12S24 | killer cell le | 4.074425 | 4.022414 | 3.586655 | 7.508911 | 6.848943 | 6.879912 | 44.84504 | 3.77259 | 0.000548 | 0.002476 | 2.606302 | 13.66667 |
| SERPINE1 | SERPINE1 | 18787 | PAI-1\|PAI1 | serine (or c | 2.004035 | 1.952024 | 3.323621 | 5.400387 | 4.920026 | 5.405981 | 14.26439 | 3.787643 | 0.00684 | 0.014716 | 1.83221 | 13.81002 |
| VCAM1 | VCAM1 | 22329 | CD106\|Vca | vascular ce | 3.382547 | 4.67449 | 3.940292 | 7.455669 | 7.389511 | 7.39256 | 52.19519 | 3.981157 | 0.002112 | 0.005865 | 2.231723 | 15.79239 |
| CXCL1 | CXCL1 | 14825 | Fsp\|Gro1\| | chemokine | 2.852032 | 2.952024 | 3.41673 | 5.648314 | 5.892011 | 5.738556 | 21.35662 | 3.996529 | 0.001996 | 0.005674 | 2.246126 | 15.96155 |
| OASL1 | OASL1 | 231655 | 7530414C1 | 2'-5' oligoa | 5.436995 | 4.915499 | 6.103231 | 9.67185 | 9.26398 | 9.372193 | 176.1456 | 4.033511 | 0.000167 | 0.001115 | 2.952616 | 16.376 |
| GBP3 | GBP3 | 55932 | AW228655 | guanylate b | 5.891561 | 5.67449 | 5.842995 | 9.985349 | 9.668735 | 9.775215 | 223.8504 | 4.139373 | 4.76E-06 | 0.000169 | 3.772106 | 17.62282 |
| TNFSF10 | TNFSF10 | 22035 | A330042I2 | tumor necr | 5.463467 | 5.022414 | 6.03144 | 9.923949 | 9.588092 | 9.74649 | 197.9934 | 4.353323 | 4.89E-05 | 0.000565 | 3.248042 | 20.44 |
| IL1B | IL1B | 16176 | IL-1beta\|Il- | interleukin | 3.76957 | 2.800021 | 3.809048 | 6.740237 | 6.504988 | 6.800841 | 33.6092 | 4.413597 | 0.012475 | 0.023664 | 1.625912 | 21.31205 |
| CXCL2 | CXCL2 | 20310 | CINC-2a\|G | chemokine | 4.540088 | 4.089528 | 4.274711 | 8.360389 | 8.213939 | 8.34458 | 79.00503 | 4.442472 | 3.09E-05 | 0.000438 | 3.358206 | 21.7429 |
| IL27 | IL27 | 246779 | IL-27\|IL-27 | interleukin | 2.004035 | -0.3699 | 1.738658 | 5.48785 | 5.504988 | 5.458448 | 9.876588 | 4.873803 | 1.40E-06 | 8.67E-05 | 4.061794 | 29.31978 |
| FPR1 | FPR1 | 14293 | FPR\|LXA4R | formyl pep | 3.141539 | 3.089528 | 2.738658 | 6.377667 | 6.348869 | 6.236056 | 25.2006 | 4.977136 | 0.000545 | 0.002476 | 2.606302 | 31.49685 |
| LIPG | LIPG | 16891 | 3110013K0 | lipase_ enc | 2.004035 | 1.630096 | 2.876162 | 6.258368 | 5.348869 | 5.899021 | 16.03062 | 5.033423 | 0.00321 | 0.008067 | 2.093305 | 32.75 |
| GBP5 | GBP5 | 229898 | 5330409J0 | guanylate b | 7.021957 | 6.500461 | 6.859674 | 12.13721 | 11.68872 | 11.80844 | 646.3076 | 5.148095 | 7.98E-06 | 0.000233 | 3.632238 | 35.45938 |
| CXCL10 | CXCL10 | 15945 | C7\|CRG-2\| | chemokine | 8.597987 | 8.169255 | 8.659904 | 15.07364 | 14.77902 | 14.83217 | 3293.326 | 6.431054 | 1.35E-06 | 8.67E-05 | 4.061794 | 86.28598 |
| CXCL11 | CXCL11 | 56066 | Cxc11\|H17 | chemokine | 3.141539 | 3.215059 | 3.224085 | 8.801266 | 7.848943 | 8.213336 | 53.47174 | 6.534497 | 0.000284 | 0.001586 | 2.799646 | 92.7 |
| PTGS2 | PTGS2 | 19225 | COX2\|Cox- | prostagland | 2.004035 | 3.089528 | 3.323621 | 8.128307 | 7.568314 | 8.068946 | 41.17772 | 6.627273 | 0.000332 | 0.001754 | 2.756075 | 98.85714 |

*Ripk2* $^{104Asp}$ bone marrow derived macrophages - MDP treated vs control

Nanostring Fibrosis Panel

**Downregulated**

| upperSymb | Symbol | GeneID | Alias | Descriptior | 20210614_ | 20210614_ | 20210614_ | 20210302_ | 20210302_ | 20210302_ | baseMean | log2FoldCh | pvalue | padj | log_10_pa | FC |
|---|---|---|---|---|---|---|---|---|---|---|---|---|---|---|---|---|
| EEF2K | EEF2K | 13631 | C86191\|eE | eukaryotic | 5.957208 | 6.031251 | 5.875693 | 4.187986 | 3.513182 | 4.137655 | 30.92059 | -5.7827 | 5.38E-08 | 8.86E-06 | 5.05273 | 55.05103 |
| ANGPTL4 | ANGPTL4 | 57875 | Arp4\|Bk89 | angiopoiet | 7.328839 | 7.659282 | 7.781365 | 3.187986 | 3.683107 | 4.722618 | 52.97352 | -5.56224 | 0.000266 | 0.001037 | 2.984244 | 47.25 |
| PTCH1 | PTCH1 | 19206 | A230106A: | patched hc | 4.112183 | 3.986857 | 3.379267 | 4.062455 | 3.098145 | 3.722618 | 13.24082 | -3.19649 | 5.02E-05 | 0.000317 | 3.498291 | 9.167284 |
| CXCR4 | CXCR4 | 12767 | CD184\|Cm | chemokine | 10.15496 | 10.01015 | 10.06877 | 8.165266 | 8.028882 | 7.575061 | 512.1822 | -2.22269 | 9.37E-05 | 0.000492 | 3.307769 | 4.66763 |
| CEBPA | CEBPA | 12606 | C/ebpalpha | CCAAT/enh | 9.657648 | 9.526015 | 9.648727 | 7.850951 | 7.52441 | 7.671634 | 400.7515 | -2.03124 | 3.37E-05 | 0.000238 | 3.623171 | 4.087548 |
| GAB1 | GAB1 | 14388 | AA408973 | growth fac | 6.423385 | 6.446288 | 6.442276 | 5.062455 | 5.420073 | 5.196549 | 56.95844 | -1.94184 | 0.002615 | 0.006396 | 2.194117 | 3.841945 |
| PELI2 | PELI2 | 93834 | AW047589 | pellino 2 | 6.276159 | 6.272259 | 6.404802 | 5.357911 | 5.157039 | 5.137655 | 54.47934 | -1.78355 | 0.000137 | 0.000661 | 3.179477 | 3.442721 |
| PLCG1 | PLCG1 | 18803 | AI894140\| | phospholip | 5.624082 | 5.794212 | 5.379267 | 5.062455 | 4.420073 | 5.076255 | 37.42829 | -1.68344 | 0.004624 | 0.009976 | 2.001055 | 3.211933 |
| LPAR5 | LPAR5 | 381810 | GPR93\|Gr | lysophosph | 7.930943 | 7.768216 | 7.8293 | 6.533761 | 6.490462 | 6.280613 | 140.9346 | -1.66329 | 0.000139 | 0.000661 | 3.179477 | 3.167381 |
| KIF3A | KIF3A | 16568 | Kif3\|Kifl\|K | kinesin fan | 7.015507 | 7.183254 | 6.902829 | 5.888426 | 5.9055 | 5.838095 | 87.76669 | -1.54555 | 0.000134 | 0.000659 | 3.18128 | 2.919153 |
| LRP6 | LRP6 | 16974 | C030016K1 | low density | 7.861121 | 7.807036 | 7.80073 | 6.625391 | 6.346072 | 6.618782 | 144.66 | -1.52238 | 4.21E-05 | 0.000285 | 3.544956 | 2.872639 |
| ARRB1 | ARRB1 | 109689 | 1200006I1 | arrestin_ b | 8.135372 | 8.222073 | 8.091548 | 6.831842 | 6.816963 | 7.137655 | 186.0108 | -1.39179 | 0.000321 | 0.001193 | 2.923504 | 2.624033 |
| RORA | RORA | 19883 | 9530021D: | RAR-relate | 6.964626 | 6.909689 | 6.741837 | 5.924952 | 5.76111 | 6.167403 | 85.1304 | -1.27271 | 0.000816 | 0.002443 | 2.612136 | 2.416154 |
| SKP2 | SKP2 | 27401 | 4930500A( | S-phase kir | 6.043796 | 5.986857 | 6.286157 | 5.357911 | 5.371163 | 5.459583 | 53.85137 | -1.26327 | 0.001368 | 0.003652 | 2.437465 | 2.400399 |
| MAP3K1 | MAP3K1 | 26401 | MAPKKK1\| | mitogen-ac | 8.86707 | 8.931325 | 8.913986 | 7.732307 | 7.816963 | 7.88361 | 327.9981 | -1.18136 | 3.13E-05 | 0.000224 | 3.649627 | 2.267902 |
| GNG2 | GNG2 | 14702 | - | guanine nu | 8.177474 | 8.126408 | 7.968517 | 6.812477 | 7.113095 | 7.320877 | 192.2014 | -1.16178 | 0.003481 | 0.008111 | 2.090911 | 2.237332 |
| KRAS | KRAS | 16653 | AI929937\| | v-Ki-ras2 Ki | 6.507042 | 6.561765 | 6.638001 | 5.888426 | 5.600645 | 5.910245 | 72.72372 | -1.15358 | 0.000271 | 0.001037 | 2.984244 | 2.224656 |
| CKAP4 | CKAP4 | 216197 | 5630400A( | cytoskeleto | 7.311492 | 7.26608 | 7.265399 | 6.384383 | 6.395826 | 6.385583 | 114.1505 | -1.14068 | 1.21E-05 | 0.000128 | 3.894093 | 2.204856 |
| SKI | SKI | 108077 | 4930534J0 | superkiller | 7.957208 | 7.889734 | 7.959929 | 7.172879 | 6.988916 | 6.995636 | 180.2744 | -1.031 | 7.29E-05 | 0.000419 | 3.378008 | 2.043439 |
| TBL1XR1 | TBL1XR1 | 81004 | 8030499H( | transducin | 8.264186 | 8.218878 | 8.286157 | 7.423203 | 7.383547 | 7.398183 | 227.3901 | -0.96897 | 2.51E-06 | 4.97E-05 | 4.303691 | 1.957444 |
| SCD1 | SCD1 | 20249 | AA589638 | stearoyl-Cc | 5.264186 | 5.332631 | 5.126501 | 5.24688 | 4.76111 | 5.012124 | 34.86978 | -0.96143 | 0.001891 | 0.00484 | 2.31519 | 1.94724 |
| SCD2 | SCD2 | 20250 | Mir5114\|S | stearoyl-Cc | 10.93567 | 10.98873 | 10.91454 | 9.927204 | 10.14618 | 10.08991 | 1448.529 | -0.90958 | 0.000227 | 0.000944 | 3.024935 | 1.878496 |
| EPAS1 | EPAS1 | 13819 | HIF-2alpha | endothelia | 6.706024 | 6.909689 | 7.071758 | 6.275449 | 6.213622 | 6.280613 | 95.41819 | -0.90025 | 0.001923 | 0.004897 | 2.310113 | 1.866393 |
| PRKDC | PRKDC | 19090 | AI326420\| | protein kin | 5.714847 | 5.531177 | 5.638001 | 5.303463 | 5.26807 | 5.360048 | 44.30101 | -0.87789 | 0.001356 | 0.003641 | 2.438729 | 1.837685 |
| ALDH9A1 | ALDH9A1 | 56752 | AA139417 | aldehyde d | 9.36992 | 9.3488 | 9.345051 | 8.521887 | 8.66293 | 8.489331 | 496.7306 | -0.84641 | 0.000167 | 0.000764 | 3.116859 | 1.798026 |
| NCOR2 | NCOR2 | 20602 | N-CoR\|SM | nuclear rec | 8.105488 | 8.212466 | 8.156874 | 7.533761 | 7.281367 | 7.495207 | 222.4793 | -0.82922 | 0.00042 | 0.001461 | 2.835255 | 1.77673 |
| NFAM1 | NFAM1 | 74039 | 4921501M | Nfat activa | 9.002954 | 9.083133 | 9.037478 | 8.303463 | 8.320537 | 8.260052 | 406.7324 | -0.80559 | 1.93E-05 | 0.000174 | 3.760189 | 1.747863 |
| CASP6 | CASP6 | 12368 | CASP-6\|Mc | caspase 6 | 8.171077 | 8.143361 | 8.118806 | 7.461005 | 7.642465 | 7.280613 | 223.3075 | -0.79227 | 0.003263 | 0.007688 | 2.11416 | 1.731797 |
| CDKN2C | CDKN2C | 12580 | C77269\|IN | cyclin-depe | 8.638066 | 8.586769 | 8.49969 | 7.897644 | 7.964394 | 7.810081 | 300.8236 | -0.76293 | 0.000337 | 0.001234 | 2.908663 | 1.696933 |
| FNIP2 | FNIP2 | 329679 | D630023B: | folliculin in | 9.455327 | 9.48803 | 9.50855 | 8.727145 | 8.703007 | 8.874621 | 558.771 | -0.75744 | 0.000177 | 0.000788 | 3.103489 | 1.690487 |
| ARHGEF6 | ARHGEF6 | 73341 | 1600028C( | Rac/Cdc42 | 8.524714 | 8.533751 | 8.502649 | 7.960576 | 7.807803 | 7.791097 | 291.3814 | -0.74673 | 0.000135 | 0.000659 | 3.18128 | 1.67798 |
| INPP5D | INPP5D | 16331 | SHIP\|SHIP- | inositol pol | 7.732335 | 7.811285 | 7.746837 | 7.029288 | 7.26807 | 7.239193 | 177.4376 | -0.71603 | 0.002349 | 0.005861 | 2.23206 | 1.642655 |
| ACAA2 | ACAA2 | 52538 | 0610011L0 | acetyl-Coe | 7.923349 | 7.964137 | 7.786231 | 7.275449 | 7.26807 | 7.347109 | 193.2142 | -0.71299 | 0.000222 | 0.000937 | 3.028307 | 1.639201 |
| HMOX1 | HMOX1 | 15368 | D8Wsu38e | heme oxyg | 10.16143 | 10.16428 | 10.17924 | 9.603024 | 9.470284 | 9.419971 | 912.0931 | -0.69529 | 0.00022 | 0.000937 | 3.028307 | 1.619207 |
| CCNA2 | CCNA2 | 12428 | AA408589 | cyclin A2 | 8.828971 | 8.865418 | 8.788658 | 8.310383 | 8.135234 | 8.160023 | 365.767 | -0.68711 | 0.000232 | 0.000955 | 3.02005 | 1.610055 |
| PRKACB | PRKACB | 18749 | CbPKA\|Pka | protein kin | 8.676967 | 8.730569 | 8.698616 | 8.110818 | 8.067771 | 8.122548 | 338.0787 | -0.6675 | 4.02E-06 | 6.21E-05 | 4.206767 | 1.588314 |
| LDLRAP1 | LDLRAP1 | 100017 | AA691260 | low density | 9.394842 | 9.473356 | 9.463695 | 8.924952 | 8.888222 | 8.692244 | 564.0002 | -0.64815 | 0.00128 | 0.003455 | 2.46155 | 1.56716 |
| LATS2 | LATS2 | 50523 | 4932411G( | large tumo | 7.692686 | 7.772581 | 7.599818 | 7.357911 | 7.067771 | 7.107282 | 172.8058 | -0.63694 | 0.002958 | 0.007094 | 2.149128 | 1.55503 |
| PCCB | PCCB | 66904 | 1300012P( | propionyl ( | 7.379663 | 7.596651 | 7.392091 | 6.906804 | 6.956126 | 7.044546 | 148.328 | -0.63549 | 0.002035 | 0.005155 | 2.287781 | 1.553467 |
| EGR1 | EGR1 | 13653 | A530045N: | early growt | 6.190185 | 6.272259 | 6.326799 | 5.995341 | 5.798585 | 6.044546 | 68.81347 | -0.63019 | 0.001557 | 0.004091 | 2.38812 | 1.547773 |
| ERN1 | ERN1 | 78943 | 9030414B1 | endoplasm | 7.566749 | 7.645068 | 7.638001 | 7.062455 | 7.157039 | 7.167403 | 165.7409 | -0.61957 | 0.000555 | 0.001791 | 2.746886 | 1.536421 |
| PARP1 | PARP1 | 11545 | 5830444G: | poly (ADP-r | 8.417991 | 8.559241 | 8.648727 | 8.118723 | 7.9055 | 7.987321 | 309.3116 | -0.60695 | 0.002665 | 0.006486 | 2.188005 | 1.523032 |
| FLI1 | FLI1 | 14247 | EWSR2\|Fli- | Friend leuk | 9.336006 | 9.351721 | 9.249631 | 8.822192 | 8.642465 | 8.814788 | 524.9855 | -0.59316 | 0.000678 | 0.002069 | 2.684262 | 1.508551 |

**Upregulated**

| | | | | | | | | | | | | | | | | |
|---|---|---|---|---|---|---|---|---|---|---|---|---|---|---|---|---|
| CUL1 | CUL1 | 26965 | - | cullin 1 | 8.86707 | 8.917594 | 8.987654 | 9.50389 | 9.541088 | 9.535686 | 598.6202 | 0.587737 | 0.000139 | 0.000661 | 3.179477 | 1.502888 |
| SOS1 | SOS1 | 20662 | 4430401P( | son of seve | 6.732335 | 6.917594 | 6.92064 | 7.591708 | 7.44392 | 7.541377 | 146.1456 | 0.609876 | 0.001729 | 0.004449 | 2.351762 | 1.526128 |
| SPOP | SPOP | 20747 | AI315626\| | speckle-typ | 7.071541 | 7.278411 | 7.031343 | 7.906804 | 7.789307 | 7.671634 | 175.8466 | 0.611465 | 0.004105 | 0.009095 | 2.041213 | 1.52781 |
| CYP27A1 | CYP27A1 | 104086 | 1300013A( | cytochrom | 7.566749 | 7.601566 | 7.781365 | 8.397439 | 8.254649 | 8.239193 | 251.3395 | 0.61312 | 0.002399 | 0.005927 | 2.227197 | 1.529563 |
| NCF1 | NCF1 | 17969 | NCF-47K\|N | neutrophil | 10.04292 | 10.04575 | 10.07971 | 10.69578 | 10.56982 | 10.77549 | 1321.76 | 0.617697 | 0.000522 | 0.001709 | 2.767177 | 1.534424 |
| H2-D1 | H2-D1 | 14964 | H-2D\|H2-D | histocompa | 13.4441 | 13.44774 | 13.48564 | 14.10995 | 14.09321 | 14.0765 | 14029.62 | 0.633447 | 2.76E-06 | 5.09E-05 | 4.292858 | 1.551267 |
| PTPN6 | PTPN6 | 15170 | 70Z-SHP\|H | protein tyr | 10.10213 | 10.13321 | 10.07375 | 10.74001 | 10.72264 | 10.79943 | 1378.161 | 0.644806 | 2.10E-05 | 0.000176 | 3.754323 | 1.563529 |
| CSF3R | CSF3R | 12986 | Cd114\|Csfg | colony stim | 7.512113 | 7.451743 | 7.627194 | 8.225075 | 8.157039 | 8.280613 | 234.8555 | 0.65553 | 0.00064 | 0.002014 | 2.695873 | 1.575194 |
| PLCB3 | PLCB3 | 18797 | mKIAA409: | phospholip | 7.328839 | 6.97175 | 7.222963 | 7.888426 | 7.844099 | 8.003904 | 186.5386 | 0.695178 | 0.004533 | 0.009822 | 2.007779 | 1.619084 |
| GNPTAB | GNPTAB | 432486 | EG432486\| | N-acetylglu | 8.382435 | 8.296712 | 8.226548 | 9.012415 | 8.984858 | 9.068393 | 405.0323 | 0.700059 | 0.000121 | 0.000616 | 3.210255 | 1.624571 |
| MOB1B | MOB1B | 68473 | 1110003E( | MOB kinas | 7.233812 | 7.176681 | 6.998177 | 7.879148 | 7.879505 | 7.910245 | 182.6487 | 0.709128 | 0.00037 | 0.001317 | 2.88055 | 1.634816 |
| IL1RAP | IL1RAP | 16180 | 6430709H( | interleukin | 5.896454 | 6.170078 | 5.929464 | 6.942874 | 6.642465 | 6.819479 | 84.45645 | 0.711684 | 0.003906 | 0.008772 | 2.056921 | 1.637715 |
| EP400 | EP400 | 75560 | 1700020J0 | E1A bindin | 7.190185 | 7.074319 | 7.251392 | 8.029288 | 7.816963 | 8.020298 | 189.1964 | 0.744868 | 0.001057 | 0.002935 | 2.532459 | 1.675821 |
| CD14 | CD14 | 12475 | - | CD14 antig | 10.54155 | 10.75228 | 10.71274 | 11.42559 | 11.3897 | 11.43915 | 2110.696 | 0.745771 | 0.000368 | 0.001317 | 2.88055 | 1.67687 |
| RAP1B | RAP1B | 215449 | 2810443E1 | RAS relatec | 12.48529 | 12.59634 | 12.54597 | 13.31748 | 13.31428 | 13.37068 | 7849.259 | 0.790708 | 2.84E-05 | 0.000213 | 3.671459 | 1.729923 |
| RELA | RELA | 19697 | p65 | v-rel reticu | 6.22766 | 6.272259 | 6.286157 | 7.317269 | 6.922573 | 7.122548 | 103.3512 | 0.792228 | 0.002285 | 0.005758 | 2.239711 | 1.731747 |
| CD180 | CD180 | 17079 | F630107B1 | CD180 anti | 12.00205 | 12.07498 | 12.04866 | 12.89045 | 12.78931 | 12.88193 | 5587.147 | 0.810768 | 3.05E-05 | 0.000221 | 3.655003 | 1.754145 |
| CAPN5 | CAPN5 | 12337 | nCL-3 | calpain 5 | 6.32308 | 6.561765 | 6.353271 | 7.371208 | 7.395826 | 7.091852 | 115.3201 | 0.811516 | 0.004411 | 0.0096 | 2.017716 | 1.755055 |
| GPR65 | GPR65 | 14744 | Dig1\|Gpcr: | G-protein ( | 9.704917 | 9.719307 | 9.613571 | 10.48719 | 10.40497 | 10.60662 | 1089.484 | 0.813953 | 0.000249 | 0.000999 | 3.000434 | 1.758022 |
| TCF7L2 | TCF7L2 | 21416 | TCF4B\|TCF | transcriptic | 6.098762 | 6.183254 | 6.126501 | 7.110818 | 6.853032 | 7.122548 | 95.83535 | 0.823181 | 0.000651 | 0.002036 | 2.69122 | 1.769303 |
| CASP8 | CASP8 | 12370 | CASP-8\|FLI | caspase 8 | 9.169473 | 9.150087 | 9.128418 | 10.00177 | 9.941543 | 10.01622 | 758.9797 | 0.828249 | 3.51E-06 | 5.78E-05 | 4.238037 | 1.775529 |
| H2-DMA | H2-DMA | 14998 | H-2Ma\|H2 | histocompa | 7.334575 | 7.332631 | 7.514426 | 8.351216 | 8.120513 | 8.300886 | 226.8678 | 0.835483 | 0.000961 | 0.002743 | 2.561736 | 1.784454 |
| STAT3 | STAT3 | 20848 | 1110034C( | signal trans | 8.34598 | 8.393416 | 8.385693 | 9.199213 | 9.160641 | 9.379241 | 449.0391 | 0.856923 | 0.000251 | 0.001002 | 2.999283 | 1.811172 |
| NCF4 | NCF4 | 17972 | AI451400\| | neutrophil | 10.12961 | 10.01568 | 10.11688 | 10.99749 | 10.89581 | 10.95464 | 1466.692 | 0.857539 | 5.02E-05 | 0.000317 | 3.498291 | 1.811944 |
| H2-AA | H2-AA | 14960 | Aalpha\|H-: | histocompa | 8.371316 | 8.393416 | 8.439191 | 9.282504 | 9.29126 | 9.266938 | 458.4977 | 0.864939 | 5.51E-06 | 7.17E-05 | 4.144774 | 1.821262 |
| TXN1 | TXN1 | 22166 | ADF\|AW55 | thioredoxir | 12.22399 | 12.23498 | 12.1674 | 13.12487 | 13.01974 | 13.11684 | 6418.282 | 0.877372 | 2.41E-05 | 0.000195 | 3.710091 | 1.837026 |
| H2-AB1 | H2-AB1 | 14961 | AI845868\| | histocompa | 7.723618 | 7.789911 | 7.938234 | 8.695781 | 8.712855 | 8.786312 | 309.6408 | 0.895085 | 0.000316 | 0.001191 | 2.923938 | 1.85972 |
| PTPRC | PTPRC | 19264 | B220\|CD4! | protein tyr | 10.79349 | 10.83704 | 10.76851 | 11.72391 | 11.63212 | 11.7456 | 2435.687 | 0.898354 | 2.19E-05 | 0.000181 | 3.743334 | 1.863938 |
| PTAFR | PTAFR | 19204 | PAFR | platelet-ac | 10.86261 | 10.87458 | 10.88536 | 11.84143 | 11.69992 | 11.80003 | 2569.588 | 0.903942 | 2.78E-05 | 0.000213 | 3.671459 | 1.871172 |
| PTK2 | PTK2 | 14083 | FAK\|FRNK | PTK2 prote | 5.766686 | 6.060106 | 5.911762 | 6.995341 | 6.835111 | 6.978957 | 85.90442 | 0.96148 | 0.000896 | 0.002602 | 2.584619 | 1.947306 |
| MAPK9 | MAPK9 | 26420 | AI851083\| | mitogen-ac | 7.52722 | 7.551641 | 7.761736 | 8.716766 | 8.46155 | 8.661217 | 276.9257 | 0.981685 | 0.000925 | 0.002671 | 2.573264 | 1.974771 |

| | | | | | | | | | | | | | | | |
|---|---|---|---|---|---|---|---|---|---|---|---|---|---|---|---|
| H2-M3 | H2-M3 | 14991 | H-2M3\|Hm | histocompa | 7.745313 | 7.728324 | 7.442276 | 8.711548 | 8.626925 | 8.666435 | 284.7338 | 1.011023 | 0.000437 | 0.001489 | 2.827238 | 2.01534 |
| MMP13 | MMP13 | 17386 | Clg\|MMP-1 | matrix met | 8.218382 | 8.387736 | 8.21937 | 9.381101 | 9.234282 | 9.280613 | 441.6972 | 1.012054 | 0.000127 | 0.000638 | 3.195474 | 2.01678 |
| MECP2 | MECP2 | 17257 | 1500041B( | methyl Cp( | 5.642698 | 5.811285 | 5.721659 | 6.792848 | 6.779969 | 6.856473 | 77.03748 | 1.020299 | 0.000157 | 0.000726 | 3.138953 | 2.028339 |
| CHMP4B | CHMP4B | 75608 | 2010012F( | charged m | 9.614683 | 9.770401 | 9.765437 | 10.81734 | 10.72385 | 10.75849 | 1210.763 | 1.045902 | 5.86E-05 | 0.000353 | 3.45251 | 2.064657 |
| MAP2K1 | MAP2K1 | 26395 | MAPKK1\|M | mitogen-ac | 9.14193 | 9.100599 | 9.177391 | 10.23602 | 10.12973 | 10.22865 | 814.095 | 1.052507 | 1.33E-05 | 0.000136 | 3.865049 | 2.074131 |
| ADAM17 | ADAM17 | 11491 | CD156b\|Ta | a disintegri | 10.9281 | 10.96509 | 10.92836 | 12.0645 | 11.92522 | 12.087 | 2862.472 | 1.083532 | 3.05E-05 | 0.000221 | 3.655003 | 2.119218 |
| H2-K1 | H2-K1 | 14972 | H-2K\|H-2K | histocompa | 12.5704 | 12.53776 | 12.5698 | 13.70369 | 13.57168 | 13.6666 | 8800.202 | 1.087511 | 1.16E-05 | 0.000124 | 3.905103 | 2.125072 |
| FCGR4 | FCGR4 | 246256 | 4833442P2 | Fc receptor | 10.65478 | 10.73393 | 10.74808 | 11.89937 | 11.74979 | 11.82649 | 2467.341 | 1.111315 | 2.85E-05 | 0.000213 | 3.671459 | 2.160424 |
| LYN | LYN | 17096 | AA407514 | Yamaguchi | 9.973844 | 9.999016 | 9.986597 | 11.12069 | 11.13157 | 11.11779 | 1504.344 | 1.134215 | 2.77E-08 | 8.86E-06 | 5.05273 | 2.194991 |
| PIK3R5 | PIK3R5 | 320207 | AV230647 | phosphoino | 7.450053 | 7.384888 | 7.149341 | 8.551391 | 8.478967 | 8.410674 | 239.5559 | 1.134661 | 0.000268 | 0.001037 | 2.984244 | 2.195669 |
| CASP7 | CASP7 | 12369 | AI314680\| | caspase 7 | 7.960922 | 7.994351 | 8.043587 | 9.225075 | 9.013028 | 9.314244 | 385.8424 | 1.175701 | 0.000224 | 0.000937 | 3.028307 | 2.259027 |
| TLR6 | TLR6 | 21899 | - | toll-like rec | 7.481417 | 7.440813 | 7.346699 | 8.497841 | 8.568465 | 8.781511 | 259.4761 | 1.17852 | 0.000239 | 0.000978 | 3.009862 | 2.263445 |
| TRAF2 | TRAF2 | 22030 | AI325259 | TNF recept | 7.282108 | 7.53632 | 7.442276 | 8.608648 | 8.746802 | 8.50106 | 259.4901 | 1.184852 | 0.000463 | 0.001537 | 2.813412 | 2.273401 |
| MMP9 | MMP9 | 17395 | AW743869 | matrix met | 8.609961 | 8.70339 | 8.67782 | 9.853322 | 9.793953 | 9.925558 | 613.3928 | 1.188352 | 1.52E-05 | 0.00015 | 3.822881 | 2.278922 |
| H2-T23 | H2-T23 | 15040 | 37b\|37c\|H | histocompa | 11.10968 | 11.06501 | 11.20172 | 12.36665 | 12.26598 | 12.34751 | 3387.705 | 1.200303 | 1.93E-05 | 0.000174 | 3.760189 | 2.297879 |
| TLR1 | TLR1 | 21897 | - | toll-like rec | 9.393469 | 9.447654 | 9.469757 | 10.63093 | 10.6099 | 10.68069 | 1051.864 | 1.200414 | 3.02E-06 | 5.26E-05 | 4.278667 | 2.298055 |
| NFKB1 | NFKB1 | 18033 | NF-KB1\|NF | nuclear fac | 8.701591 | 8.766029 | 8.746837 | 9.984567 | 10.05621 | 9.991485 | 663.7305 | 1.268399 | 2.78E-06 | 5.09E-05 | 4.292858 | 2.408941 |
| F11R | F11R | 16456 | 9130004G2 | F11 recepto | 5.029721 | 5.074319 | 4.964229 | 6.485667 | 6.241103 | 6.385583 | 51.86791 | 1.302734 | 2.57E-05 | 0.000201 | 3.695777 | 2.466959 |
| ITGA5 | ITGA5 | 16402 | Cd49e\|Fnr | integrin alp | 7.029721 | 6.986857 | 7.299832 | 8.429572 | 8.513182 | 8.410674 | 219.5349 | 1.337309 | 0.000355 | 0.001279 | 2.89314 | 2.526795 |
| CCL2 | CCL2 | 20296 | AI323594\| | chemokine | 11.69241 | 11.88066 | 11.80852 | 13.08078 | 13.13249 | 13.25876 | 5695.485 | 1.363151 | 5.80E-05 | 0.000353 | 3.45251 | 2.572465 |
| CYBB | CYBB | 13058 | C88302\|CG | cytochrom | 12.36415 | 12.45582 | 12.44208 | 13.81947 | 13.73522 | 13.8323 | 8829.854 | 1.374789 | 5.08E-06 | 6.96E-05 | 4.15708 | 2.5933 |
| BST2 | BST2 | 69550 | 2310015I1 | bone marr | 11.12796 | 11.18816 | 11.20808 | 12.60689 | 12.50364 | 12.5688 | 3735.945 | 1.38462 | 3.64E-06 | 5.80E-05 | 4.236805 | 2.611032 |
| NRP2 | NRP2 | 18187 | 1110048P( | neuropilin | 10.23151 | 10.2036 | 10.15311 | 11.5803 | 11.58598 | 11.61001 | 1903.035 | 1.395136 | 4.90E-07 | 2.20E-05 | 4.657582 | 2.630134 |
| AXL | AXL | 26362 | AI323647\| | AXL recept | 8.872995 | 8.903731 | 8.913986 | 10.43116 | 10.14981 | 10.30591 | 774.0403 | 1.396873 | 6.88E-05 | 0.0004 | 3.39782 | 2.633301 |
| PHLPP1 | PHLPP1 | 98432 | AI836256\| | PH domain | 6.741 | 6.948789 | 6.810317 | 8.324123 | 8.199683 | 8.203744 | 185.8433 | 1.40124 | 5.01E-05 | 0.000317 | 3.498291 | 2.641286 |
| IL10RA | IL10RA | 16154 | AW553859 | interleukin | 8.884772 | 8.838606 | 8.812703 | 10.30693 | 10.27141 | 10.31092 | 760.5442 | 1.44953 | 2.85E-07 | 1.58E-05 | 4.801323 | 2.731191 |
| PSMB8 | PSMB8 | 16913 | Lmp-7\|Lmp | proteasom | 10.53969 | 10.58242 | 10.65738 | 12.11181 | 11.96182 | 12.08407 | 2561.665 | 1.459076 | 1.46E-05 | 0.000147 | 3.83173 | 2.749321 |
| HC | HC | 15139 | C5\|C5a\|He | hemolytic c | 6.029721 | 6.284537 | 6.659375 | 8.062455 | 7.642465 | 7.865576 | 136.3044 | 1.513943 | 0.001242 | 0.003372 | 2.47213 | 2.855895 |
| CXCL16 | CXCL16 | 66102 | 0910001K2 | chemokine | 10.56736 | 10.52537 | 10.40401 | 12.07216 | 11.98841 | 12.01059 | 2454.673 | 1.524573 | 9.38E-06 | 0.000103 | 3.987426 | 2.877015 |
| FLNB | FLNB | 286940 | AL024016\| | filamin_ be | 6.32308 | 6.074319 | 6.392091 | 7.802696 | 7.807803 | 7.838095 | 131.5694 | 1.553769 | 0.000186 | 0.000811 | 3.091034 | 2.935831 |
| HCK | HCK | 15162 | AI849071\| | hemopoiet | 9.146829 | 9.133213 | 9.141767 | 10.66372 | 10.65784 | 10.77428 | 968.5 | 1.557874 | 2.13E-06 | 4.79E-05 | 4.31946 | 2.944197 |
| THBS1 | THBS1 | 21825 | TSP-1\|TSP1 | thrombosp | 5.642698 | 5.741744 | 5.701195 | 7.261235 | 7.241103 | 7.575061 | 92.23053 | 1.674559 | 0.000187 | 0.000811 | 3.091034 | 3.192217 |
| ITGA4 | ITGA4 | 16401 | CD49D\|Itg | integrin alp | 9.999347 | 10.11786 | 9.992925 | 11.75847 | 11.66039 | 11.74004 | 1882.158 | 1.683453 | 4.61E-06 | 6.70E-05 | 4.173992 | 3.211957 |
| LOX | LOX | 16948 | AI893619\| | lysyl oxidas | 4.566749 | 4.04575 | 4.571912 | 6.126586 | 5.972614 | 6.044546 | 37.30661 | 1.707078 | 0.004064 | 0.009044 | 2.043638 | 3.264989 |
| OAS1A/G | OAS1A/G | NA | NA | NA | 6.288033 | 6.284537 | 6.571912 | 8.225075 | 8.052341 | 8.084074 | 152.3236 | 1.753033 | 0.000169 | 0.000764 | 3.116859 | 3.370663 |
| NR1H3 | NR1H3 | 22259 | AU018371 | nuclear rec | 7.334575 | 7.296712 | 7.265399 | 9.045967 | 9.105639 | 9.072329 | 291.3825 | 1.781857 | 2.57E-07 | 1.58E-05 | 4.801323 | 3.438685 |
| RELB | RELB | 19698 | shep | avian reticu | 8.158195 | 8.001807 | 8.193964 | 10.1305 | 9.951974 | 10.09185 | 544.221 | 1.946749 | 1.88E-05 | 0.000174 | 3.760189 | 3.855047 |
| IL21R | IL21R | 60504 | NILR | interleukin | 7.239938 | 7.290637 | 7.244337 | 9.243269 | 9.153427 | 9.317564 | 304.0565 | 1.992393 | 2.12E-06 | 4.79E-05 | 4.31946 | 3.978963 |
| MET | MET | 17295 | AI838057\| | met proto- | 4.444758 | 4.209249 | 3.379267 | 5.995341 | 5.939447 | 5.80062 | 31.15619 | 1.999164 | 0.00395 | 0.008829 | 2.054078 | 3.997684 |
| XAF1 | XAF1 | 327959 | Fbox39 | XIAP associ | 8.54712 | 8.556712 | 8.537696 | 10.58602 | 10.42757 | 10.62683 | 748.0502 | 2.004928 | 4.75E-06 | 6.71E-05 | 4.173276 | 4.013688 |
| MYD88 | MYD88 | 17874 | - | myeloid dif | 8.787748 | 8.887723 | 8.493753 | 10.72585 | 10.70424 | 10.76215 | 847.4066 | 2.011927 | 6.77E-05 | 0.000398 | 3.399808 | 4.033204 |
| AIM2 | AIM2 | 383619 | Gm1313\|If | absent in n | 7.697145 | 7.750622 | 7.411116 | 9.671802 | 9.592667 | 9.666435 | 396.6248 | 2.033703 | 3.86E-05 | 0.000268 | 3.571188 | 4.094544 |
| CFLAR | CFLAR | 12633 | 2310024N1 | CASP8 and | 9.36992 | 9.476036 | 9.432999 | 11.49557 | 11.42682 | 11.52136 | 1402.498 | 2.058116 | 1.05E-06 | 3.97E-05 | 4.40087 | 4.164422 |
| CD86 | CD86 | 12524 | B7\|B7-2\|B | CD86 antig | 5.86509 | 6.19631 | 6.366328 | 8.282504 | 8.120513 | 8.114935 | 142.7763 | 2.067339 | 0.000383 | 0.001353 | 2.868796 | 4.19113 |
| CDKN1A | CDKN1A | 12575 | CAP20\|CDI | cyclin-depe | 9.607594 | 9.695366 | 9.578939 | 11.82521 | 11.68373 | 11.78031 | 1657.959 | 2.138867 | 2.35E-06 | 4.91E-05 | 4.308478 | 4.404161 |
| PELI1 | PELI1 | 67245 | 2810468L0 | pellino 1 | 8.32308 | 8.353179 | 8.414262 | 10.58173 | 10.40193 | 10.58336 | 695.9903 | 2.166867 | 5.49E-06 | 7.17E-05 | 4.144774 | 4.490471 |
| IFI35 | IFI35 | 70110 | 2010008K1 | interferon- | 9.076684 | 9.031251 | 9.077723 | 11.26034 | 11.15162 | 11.31673 | 1138.085 | 2.18597 | 1.73E-06 | 4.50E-05 | 4.346865 | 4.550325 |
| SRC | SRC | 20779 | AW259666 | Rous sarco | 5.605223 | 5.630713 | 6.272351 | 8.054235 | 7.988916 | 8.099587 | 122.9423 | 2.275532 | 0.001016 | 0.002855 | 2.544404 | 4.841761 |
| STAT1 | STAT1 | 20846 | 2010005J0 | signal trans | 10.11718 | 10.13576 | 10.22027 | 12.47877 | 12.4197 | 12.45356 | 2528.746 | 2.29561 | 4.10E-07 | 2.03E-05 | 4.692979 | 4.909614 |
| TLR2 | TLR2 | 24088 | Ly105 | toll-like rec | 7.522202 | 7.494651 | 7.560596 | 9.867466 | 9.814679 | 9.793484 | 408.8742 | 2.316193 | 2.88E-07 | 1.58E-05 | 4.801323 | 4.980162 |
| BIRC3 | BIRC3 | 11796 | AW10767C | baculoviral | 7.63342 | 7.561765 | 7.366328 | 9.879148 | 9.737184 | 9.987321 | 414.203 | 2.364155 | 2.09E-05 | 0.000176 | 3.754323 | 5.148509 |
| CCL4 | CCL4 | 20303 | AT744.1\|A | chemokine | 8.532221 | 8.774759 | 8.372812 | 10.83424 | 10.90335 | 11.02335 | 855.2031 | 2.368509 | 5.08E-05 | 0.000317 | 3.498291 | 5.164072 |
| DAXX | DAXX | 13163 | - | Fas death c | 6.833029 | 7.241098 | 7.171825 | 9.554308 | 9.395826 | 9.621471 | 315.8128 | 2.469741 | 8.65E-05 | 0.00048 | 3.318493 | 5.539445 |
| GNB4 | GNB4 | 14696 | 6720453A2 | guanine nu | 4.444758 | 4.649822 | 4.031343 | 6.647418 | 6.742001 | 6.910245 | 47.53542 | 2.572793 | 0.000155 | 0.00072 | 3.142465 | 5.9496 |
| JAK2 | JAK2 | 16452 | Fd17 | Janus kinas | 7.804379 | 7.798499 | 7.680436 | 10.35289 | 10.25802 | 10.44448 | 532.4307 | 2.608311 | 2.04E-06 | 4.79E-05 | 4.31946 | 6.097893 |
| RAPGEF2 | RAPGEF2 | 76089 | 5830453M | Rap guanin | 7.576464 | 7.710233 | 7.50855 | 10.20294 | 10.16064 | 10.28401 | 480.0828 | 2.637598 | 2.39E-06 | 4.91E-05 | 4.308478 | 6.222949 |
| OAS1B | OAS1B | 23961 | Flv\|L1\|Mn | 2'-5' oligoa | 4.264186 | 3.401894 | 3.741837 | 6.157612 | 6.320537 | 6.137655 | 32.08781 | 2.783085 | 0.003642 | 0.008408 | 2.075307 | 6.883229 |
| CASP4 | CASP4 | 12363 | CASP-11\|C | caspase 4_ | 8.343137 | 8.457177 | 8.204907 | 11.12854 | 11.06489 | 11.15632 | 846.7717 | 2.794604 | 3.09E-06 | 5.26E-05 | 4.278667 | 6.938402 |
| NLRP3 | NLRP3 | 216799 | AGTAVPRL | NLR family_ | 6.624082 | 6.510419 | 6.616306 | 9.394186 | 9.244501 | 9.489331 | 252.4413 | 2.839745 | 4.51E-06 | 6.70E-05 | 4.173992 | 7.158937 |
| FAS | FAS | 14102 | AI196731\| | Fas (TNF re | 4.264186 | 4.828159 | 4.701195 | 7.275449 | 6.972614 | 7.410674 | 60.07583 | 2.840528 | 0.000768 | 0.002313 | 2.635762 | 7.162822 |
| MMP14 | MMP14 | 17387 | AI325305\| | matrix met | 4.679223 | 5.156782 | 5.126501 | 7.647418 | 7.826065 | 8.060487 | 85.39498 | 2.997772 | 0.000147 | 0.00069 | 3.160997 | 7.987654 |
| C3 | C3 | 12266 | AI255234\| | complemer | 9.409863 | 9.810894 | 9.477299 | 12.51816 | 12.33973 | 12.44106 | 1952.806 | 3.010304 | 8.05E-07 | 3.31E-05 | 4.479731 | 8.05734 |
| HCAR2 | HCAR2 | 80885 | Gpr109a\|G | hydroxycar | 4.264186 | 4.446288 | 4.526108 | 7.24688 | 7.113095 | 7.280613 | 56.21416 | 3.067575 | 9.07E-05 | 0.000482 | 3.316928 | 8.383632 |
| MARCO | MARCO | 17167 | AI323439\| | macrophag | 4.942258 | 5.102334 | 4.701195 | 7.591708 | 7.861911 | 8.012124 | 82.62969 | 3.07109 | 5.23E-05 | 0.000323 | 3.490648 | 8.404078 |
| IRF1 | IRF1 | 16362 | AU020929 | interferon | 9.304202 | 9.304269 | 9.336783 | 12.47454 | 12.38701 | 12.49043 | 1888.302 | 3.1435 | 8.67E-08 | 1.01E-05 | 4.994806 | 8.83665 |
| IL1B | IL1B | 16176 | IL-1beta\|Il- | interleukin | 3.605223 | 3.308785 | 3.156874 | 5.772949 | 6.127892 | 6.076255 | 25.53958 | 3.240201 | 0.000245 | 0.000992 | 3.003303 | 9.449259 |
| CXCL2 | CXCL2 | 20310 | CINC-2a\|G | chemokine | 5.112183 | 5.446288 | 5.126501 | 8.24688 | 8.352386 | 8.459583 | 110.7096 | 3.271218 | 2.03E-05 | 0.000176 | 3.754323 | 9.654611 |
| FLT1 | FLT1 | 14254 | AI323757\| | FMS-like ty | 3.52722 | 4.156782 | 3.819839 | 6.924952 | 6.53555 | 6.575061 | 38.22825 | 3.329839 | 0.001385 | 0.003678 | 2.4344 | 10.05499 |
| ISG15 | ISG15 | 1E+08 | G1p2\|IGI15 | ISG15 ubiq | 9.930943 | 9.818692 | 10.05774 | 13.5161 | 13.40097 | 13.44429 | 3314.991 | 3.523851 | 1.44E-06 | 4.20E-05 | 4.377266 | 11.50231 |
| ICAM1 | ICAM1 | 15894 | CD54\|Icam | intercellula | 7.758175 | 7.785598 | 7.977053 | 11.39174 | 11.33336 | 11.36808 | 777.3036 | 3.549613 | 1.43E-06 | 4.20E-05 | 4.377266 | 11.70954 |
| FGL2 | FGL2 | 14190 | AI385601\| | fibrinogen- | 6.22766 | 6.467983 | 6.126501 | 9.839038 | 9.619091 | 9.807721 | 258.6156 | 3.555032 | 7.23E-06 | 8.31E-05 | 4.08037 | 11.75361 |
| FPR1 | FPR1 | 14293 | FPR\|LXA4R | formyl pep | 3.357295 | 3.649822 | 3.031343 | 6.24688 | 6.157039 | 6.225118 | 27.43444 | 3.559336 | 0.000128 | 0.000638 | 3.195474 | 11.78873 |
| P2RY13 | P2RY13 | 74191 | 2010001L0 | purinergic | 2.816727 | 3.649822 | 3.031343 | 6.062455 | 5.722636 | 6.385583 | 24.44432 | 3.594181 | 0.001417 | 0.003745 | 2.426603 | 12.07692 |
| TNF | TNF | 21926 | DIF\|TNF-a\| | tumor necr | 7.692686 | 7.933276 | 7.857314 | 11.38602 | 11.4004 | 11.44599 | 786.4888 | 3.608726 | 1.30E-06 | 4.20E-05 | 4.377266 | 12.1993 |
| ISG20 | ISG20 | 57444 | 1600023I0 | interferon- | 7.412578 | 7.390579 | 7.478802 | 11.28777 | 11.19618 | 11.25573 | 646.6942 | 3.854848 | 1.02E-07 | 1.01E-05 | 4.994806 | 14.46854 |
| GBP3 | GBP3 | 55932 | AW228655 | guanylate l | 6.190185 | 5.956483 | 6.031343 | 10.16144 | 9.972614 | 9.983145 | 264.8814 | 4.075861 | 1.62E-06 | 4.45E-05 | 4.351259 | 16.86384 |
| OASL1 | OASL1 | 231655 | 7530414C1 | 2'-5' oligoa | 5.566749 | 5.332631 | 5.616306 | 9.521887 | 9.377369 | 9.586116 | 181.0414 | 4.139134 | 6.82E-06 | 8.19E-05 | 4.08677 | 17.6199 |
| KDR | KDR | 16542 | 6130401C( | kinase inse | 4.679223 | 4.611347 | 3.819839 | 8.24688 | 8.307598 | 8.379241 | 81.04708 | 4.266613 | 8.39E-05 | 0.000476 | 3.322009 | 19.24768 |
| TNFSF10 | TNFSF10 | 22035 | A330042I2 | tumor necr | 5.766686 | 5.668681 | 5.721659 | 10.00604 | 9.746802 | 10.00596 | 225.8631 | 4.328718 | 1.38E-06 | 4.20E-05 | 4.377266 | 20.09434 |
| IL27 | IL27 | 246779 | IL-27\|IL-27 | interleukin | -0.6427 | 2.401894 | 0.571912 | 5.55722 | 5.600645 | 5.910245 | 9.403526 | 5.16451 | 6.96E-06 | 8.19E-05 | 4.08677 | 35.86513 |
| LIPG | LIPG | 16891 | 3110013K( | lipase_ enc | 2.357295 | 1.986857 | 2.571912 | 6.094877 | 6.157039 | 6.107282 | 18.53967 | 5.258285 | 0.000455 | 0.001519 | 2.81848 | 38.2738 |

| | | | | | | | | | | | | | | | |
|---|---|---|---|---|---|---|---|---|---|---|---|---|---|---|---|
| GBP5 | GBP5 | 229898 | 5330409J0 | guanylate I | 6.766686 | 6.941053 | 6.848036 | 12.14509 | 12.02494 | 12.08456 | 708.3867 | 5.294912 | 1.47E-07 | 1.21E-05 | 4.916226 | 39.25794 |
| VCAM1 | VCAM1 | 22329 | CD106\|Vca | vascular ce | 2.52722 | 3.209249 | 3.031343 | 7.772949 | 7.254649 | 7.629508 | 37.72598 | 5.863498 | 0.000318 | 0.001191 | 2.923938 | 58.22222 |
| PTGS2 | PTGS2 | 19225 | COX2\|Cox- | prostaglan | 3.057735 | 3.209249 | 3.379267 | 8.282504 | 8.157039 | 8.14515 | 52.17031 | 6.317146 | 0.000436 | 0.001489 | 2.827238 | 79.73528 |
| CXCL10 | CXCL10 | 15945 | C7\|CRG-2\| | chemokine | 8.529723 | 8.407519 | 8.566265 | 14.98875 | 14.90765 | 15.01308 | 3409.863 | 6.4891 | 4.44E-08 | 8.86E-06 | 5.05273 | 89.82843 |
| CXCL11 | CXCL11 | 56066 | Cxc11\|H17 | chemokine | 3.444758 | 3.209249 | 1.571912 | 8.330944 | 8.149807 | 8.24618 | 45.009 | 6.888056 | 0.000451 | 0.001519 | 2.81848 | 118.4435 |

ACL Rupture - *Ripk2* $^{104Asp}$ vs WT

RNA-Seq

**Dowregulated**

| Row.name: | baseMean | log2FoldCh | lfcSE | stat | pvalue | padj | Ensembl | Gene |
|---|---|---|---|---|---|---|---|---|
| ENSMUSG( | 111.5059 | -1.12529 | 0.245516 | -4.58335 | 4.58E-06 | 0.001251 | ENSMUSG( | Serpina3n |
| ENSMUSG( | 179.4747 | -0.98895 | 0.244711 | -4.04132 | 5.32E-05 | 0.007512 | ENSMUSG( | Ppp1r3c |
| ENSMUSG( | 3285.164 | -0.94718 | 0.1434 | -6.60517 | 3.97E-11 | 7.14E-08 | ENSMUSG( | Fmod |
| ENSMUSG( | 116.8309 | -0.91811 | 0.216059 | -4.24936 | 2.14E-05 | 0.003852 | ENSMUSG( | Medag |
| ENSMUSG( | 87.385 | -0.89437 | 0.258261 | -3.46306 | 0.000534 | 0.042519 | ENSMUSG( | Nmrk2 |
| ENSMUSG( | 260.0972 | -0.89094 | 0.189765 | -4.69497 | 2.67E-06 | 0.000835 | ENSMUSG( | Gpnmb |
| ENSMUSG( | 244.5888 | -0.86006 | 0.209717 | -4.10104 | 4.11E-05 | 0.006233 | ENSMUSG( | Clec3a |
| ENSMUSG( | 108.2651 | -0.85947 | 0.175196 | -4.90576 | 9.31E-07 | 0.000355 | ENSMUSG( | Mmp3 |
| ENSMUSG( | 27536.29 | -0.8301 | 0.125095 | -6.63571 | 3.23E-11 | 6.77E-08 | ENSMUSG( | Col2a1 |
| ENSMUSG( | 9923.499 | -0.7791 | 0.114304 | -6.81607 | 9.36E-12 | 2.35E-08 | ENSMUSG( | Neat1 |
| ENSMUSG( | 2537.306 | -0.76904 | 0.162114 | -4.74379 | 2.10E-06 | 0.000713 | ENSMUSG( | Fndc1 |
| ENSMUSG( | 79.22738 | -0.75827 | 0.21589 | -3.51231 | 0.000444 | 0.039324 | ENSMUSG( | Zfp579 |
| ENSMUSG( | 224.2379 | -0.75717 | 0.146703 | -5.16121 | 2.45E-07 | 0.000114 | ENSMUSG( | Ccdc3 |
| ENSMUSG( | 4698.716 | -0.74426 | 0.116949 | -6.36402 | 1.97E-10 | 2.75E-07 | ENSMUSG( | Comp |
| ENSMUSG( | 4582.26 | -0.73941 | 0.114301 | -6.46896 | 9.87E-11 | 1.55E-07 | ENSMUSG( | Acan |
| ENSMUSG( | 6906.089 | -0.73853 | 0.094999 | -7.77412 | 7.60E-15 | 3.19E-11 | ENSMUSG( | Prg4 |
| ENSMUSG( | 2271.252 | -0.73779 | 0.149419 | -4.93776 | 7.90E-07 | 0.000311 | ENSMUSG( | Tnn |
| ENSMUSG( | 511.3479 | -0.7176 | 0.13295 | -5.39753 | 6.76E-08 | 4.05E-05 | ENSMUSG( | Cemip |
| ENSMUSG( | 457.0384 | -0.71095 | 0.201812 | -3.52282 | 0.000427 | 0.03864 | ENSMUSG( | Asb2 |
| ENSMUSG( | 211931.1 | -0.70064 | 0.165303 | -4.23851 | 2.25E-05 | 0.003986 | ENSMUSG( | Gm26917 |
| ENSMUSG( | 651.8082 | -0.69782 | 0.149328 | -4.67306 | 2.97E-06 | 0.000889 | ENSMUSG( | Cdon |
| ENSMUSG( | 4858.488 | -0.67572 | 0.165998 | -4.07069 | 4.69E-05 | 0.006937 | ENSMUSG( | Thbs4 |
| ENSMUSG( | 490.9114 | -0.67559 | 0.194921 | -3.46599 | 0.000528 | 0.042519 | ENSMUSG( | Cilp2 |
| ENSMUSG( | 556.0406 | -0.67328 | 0.152581 | -4.41263 | 1.02E-05 | 0.002292 | ENSMUSG( | Fbn2 |
| ENSMUSG( | 10156.61 | -0.66283 | 0.124287 | -5.33301 | 9.66E-08 | 5.06E-05 | ENSMUSG( | Col12a1 |
| ENSMUSG( | 45279.38 | -0.66016 | 0.119676 | -5.51619 | 3.46E-08 | 2.29E-05 | ENSMUSG( | Fn1 |
| ENSMUSG( | 210.6626 | -0.65335 | 0.146421 | -4.46213 | 8.11E-06 | 0.001926 | ENSMUSG( | Sertad4 |
| ENSMUSG( | 440.0441 | -0.65281 | 0.16562 | -3.94161 | 8.09E-05 | 0.010284 | ENSMUSG( | Cpxm2 |
| ENSMUSG( | 503.9215 | -0.65251 | 0.184413 | -3.53829 | 0.000403 | 0.037317 | ENSMUSG( | Amd1 |
| ENSMUSG( | 1695.754 | -0.65128 | 0.121404 | -5.36455 | 8.12E-08 | 4.44E-05 | ENSMUSG( | Cilp |
| ENSMUSG( | 318.5267 | -0.63065 | 0.147373 | -4.27929 | 1.87E-05 | 0.003468 | ENSMUSG( | P4ha2 |
| ENSMUSG( | 2112.295 | -0.6273 | 0.169141 | -3.70872 | 0.000208 | 0.022589 | ENSMUSG( | Abi3bp |
| ENSMUSG( | 299.196 | -0.62723 | 0.141032 | -4.44744 | 8.69E-06 | 0.002024 | ENSMUSG( | Nt5e |
| ENSMUSG( | 294.246 | -0.62602 | 0.11625 | -5.38511 | 7.24E-08 | 4.14E-05 | ENSMUSG( | Slc39a14 |
| ENSMUSG( | 267.8773 | -0.6252 | 0.135352 | -4.61909 | 3.85E-06 | 0.001128 | ENSMUSG( | Dtx3 |
| ENSMUSG( | 8408.278 | -0.62477 | 0.139975 | -4.46345 | 8.07E-06 | 0.001926 | ENSMUSG( | Col6a2 |
| ENSMUSG( | 742.6938 | -0.62065 | 0.125285 | -4.9539 | 7.27E-07 | 0.000295 | ENSMUSG( | Thbs3 |
| ENSMUSG( | 575.3479 | -0.61557 | 0.098299 | -6.26229 | 3.79E-10 | 4.34E-07 | ENSMUSG( | Crispld2 |
| ENSMUSG( | 242.1636 | -0.60697 | 0.149917 | -4.04871 | 5.15E-05 | 0.007446 | ENSMUSG( | Trim3 |
| ENSMUSG( | 101.7729 | -0.60509 | 0.174656 | -3.46444 | 0.000531 | 0.042519 | ENSMUSG( | Gpr157 |
| ENSMUSG( | 365.9861 | -0.60192 | 0.139837 | -4.30448 | 1.67E-05 | 0.003309 | ENSMUSG( | Zfp462 |
| ENSMUSG( | 294.9438 | -0.60028 | 0.121086 | -4.95748 | 7.14E-07 | 0.000295 | ENSMUSG( | Ntn1 |

| | | | | | | | | |
|---|---|---|---|---|---|---|---|---|
| ENSMUSG( | 149.6015 | -0.59994 | 0.163995 | -3.65826 | 0.000254 | 0.02576 | ENSMUSG( | Arhgap20 |
| ENSMUSG( | 681.5599 | -0.59905 | 0.14605 | -4.10168 | 4.10E-05 | 0.006233 | ENSMUSG( | Itga11 |
| ENSMUSG( | 709.3708 | -0.59707 | 0.163026 | -3.66239 | 0.00025 | 0.02576 | ENSMUSG( | Ssc5d |
| ENSMUSG( | 966.3514 | -0.59393 | 0.132666 | -4.47686 | 7.57E-06 | 0.001906 | ENSMUSG( | Col9a3 |
| ENSMUSG( | 1548.377 | -0.59146 | 0.153489 | -3.85345 | 0.000116 | 0.013953 | ENSMUSG( | Col15a1 |
| ENSMUSG( | 1057.942 | -0.58311 | 0.132099 | -4.41417 | 1.01E-05 | 0.002292 | ENSMUSG( | Col9a2 |
| ENSMUSG( | 2470.191 | -0.58149 | 0.123405 | -4.71206 | 2.45E-06 | 0.000791 | ENSMUSG( | Col16a1 |
| ENSMUSG( | 3654.373 | -0.58103 | 0.162905 | -3.56665 | 0.000362 | 0.034719 | ENSMUSG( | Thbs2 |
| ENSMUSG( | 10971.58 | -0.57695 | 0.150371 | -3.83687 | 0.000125 | 0.01465 | ENSMUSG( | Lars2 |
| ENSMUSG( | 455.4365 | -0.57034 | 0.132735 | -4.29684 | 1.73E-05 | 0.003353 | ENSMUSG( | Loxl3 |
| ENSMUSG( | 10807.11 | -0.56851 | 0.131104 | -4.33634 | 1.45E-05 | 0.002939 | ENSMUSG( | Col6a3 |
| ENSMUSG( | 1599.809 | -0.56749 | 0.147749 | -3.8409 | 0.000123 | 0.014547 | ENSMUSG( | Vcan |
| ENSMUSG( | 499.6776 | -0.56669 | 0.164737 | -3.43997 | 0.000582 | 0.044632 | ENSMUSG( | Gm15564 |
| ENSMUSG( | 10576.24 | -0.5653 | 0.141074 | -4.00707 | 6.15E-05 | 0.008405 | ENSMUSG( | Col5a1 |
| ENSMUSG( | 53820.32 | -0.56042 | 0.140173 | -3.99807 | 6.39E-05 | 0.008546 | ENSMUSG( | Col3a1 |
| ENSMUSG( | 1420.559 | -0.56037 | 0.112911 | -4.96297 | 6.94E-07 | 0.000295 | ENSMUSG( | Col27a1 |
| ENSMUSG( | 220.6469 | -0.55513 | 0.142332 | -3.90027 | 9.61E-05 | 0.011967 | ENSMUSG( | Pkdcc |
| ENSMUSG( | 6899.825 | -0.55178 | 0.134728 | -4.09551 | 4.21E-05 | 0.006308 | ENSMUSG( | Col6a1 |
| ENSMUSG( | 489.6378 | -0.54697 | 0.114621 | -4.77197 | 1.82E-06 | 0.000637 | ENSMUSG( | Fbln7 |
| ENSMUSG( | 1779.154 | -0.54693 | 0.125502 | -4.35797 | 1.31E-05 | 0.002707 | ENSMUSG( | Prelp |
| ENSMUSG( | 456.3058 | -0.54684 | 0.145353 | -3.76215 | 0.000168 | 0.018619 | ENSMUSG( | Matn2 |
| ENSMUSG( | 5755.171 | -0.54567 | 0.137568 | -3.96653 | 7.29E-05 | 0.009556 | ENSMUSG( | Hspg2 |
| ENSMUSG( | 177.676 | -0.54406 | 0.160473 | -3.39038 | 0.000698 | 0.049556 | ENSMUSG( | Ddit4l |
| ENSMUSG( | 1250.241 | -0.54132 | 0.086628 | -6.24879 | 4.14E-10 | 4.34E-07 | ENSMUSG( | Sash1 |
| ENSMUSG( | 1105.907 | -0.53987 | 0.140848 | -3.83303 | 0.000127 | 0.014742 | ENSMUSG( | Peg3 |
| ENSMUSG( | 192.8663 | -0.53771 | 0.156762 | -3.43014 | 0.000603 | 0.045874 | ENSMUSG( | Meltf |
| ENSMUSG( | 193.2008 | -0.53395 | 0.156007 | -3.42262 | 0.00062 | 0.046188 | ENSMUSG( | Trpv4 |
| ENSMUSG( | 305.5551 | -0.53006 | 0.150715 | -3.51698 | 0.000436 | 0.03894 | ENSMUSG( | Ldlr |
| ENSMUSG( | 921.5418 | -0.5259 | 0.129555 | -4.05931 | 4.92E-05 | 0.007199 | ENSMUSG( | Chad |
| ENSMUSG( | 1027.814 | -0.5244 | 0.149506 | -3.50758 | 0.000452 | 0.039502 | ENSMUSG( | Sned1 |
| ENSMUSG( | 455.9261 | -0.51688 | 0.131116 | -3.94216 | 8.08E-05 | 0.010284 | ENSMUSG( | Boc |
| ENSMUSG( | 3627.976 | -0.51484 | 0.145563 | -3.5369 | 0.000405 | 0.037317 | ENSMUSG( | Mir6236 |
| ENSMUSG( | 151090.8 | -0.51443 | 0.120076 | -4.28415 | 1.83E-05 | 0.003459 | ENSMUSG( | Gm42418 |
| ENSMUSG( | 750.9654 | -0.51419 | 0.139062 | -3.69758 | 0.000218 | 0.023401 | ENSMUSG( | Sod3 |
| ENSMUSG( | 262.1706 | -0.51322 | 0.141626 | -3.62375 | 0.00029 | 0.028872 | ENSMUSG( | Fgfr3 |
| ENSMUSG( | 267.0767 | -0.51236 | 0.148604 | -3.44783 | 0.000565 | 0.043879 | ENSMUSG( | Errfi1 |
| ENSMUSG( | 363.0789 | -0.50833 | 0.149631 | -3.3972 | 0.000681 | 0.048658 | ENSMUSG( | Spaca6 |
| ENSMUSG( | 1012.931 | -0.50616 | 0.119895 | -4.22166 | 2.43E-05 | 0.004179 | ENSMUSG( | 2010111I01Rik |
| ENSMUSG( | 823.4786 | -0.5044 | 0.126341 | -3.99236 | 6.54E-05 | 0.008662 | ENSMUSG( | Cspg4 |
| ENSMUSG( | 391.5484 | -0.49397 | 0.137322 | -3.59716 | 0.000322 | 0.031616 | ENSMUSG( | Inhba |
| ENSMUSG( | 418.1114 | -0.48569 | 0.13235 | -3.66971 | 0.000243 | 0.025244 | ENSMUSG( | Hspb8 |
| ENSMUSG( | 445.1206 | -0.47155 | 0.137082 | -3.43991 | 0.000582 | 0.044632 | ENSMUSG( | Ecm2 |
| ENSMUSG( | 2506.029 | -0.46446 | 0.122838 | -3.78104 | 0.000156 | 0.017859 | ENSMUSG( | Pkd1 |
| ENSMUSG( | 458.7442 | -0.46083 | 0.131592 | -3.50198 | 0.000462 | 0.040063 | ENSMUSG( | Scarf2 |
| ENSMUSG( | 762.3518 | -0.45887 | 0.135553 | -3.38521 | 0.000711 | 0.049981 | ENSMUSG( | Ptgfrn |
| ENSMUSG( | 726.3196 | -0.45777 | 0.13465 | -3.39971 | 0.000675 | 0.048488 | ENSMUSG( | Htra1 |
| ENSMUSG( | 449.246 | -0.45633 | 0.130777 | -3.48937 | 0.000484 | 0.040603 | ENSMUSG( | Creb3l1 |

| | | | | | | | | |
|---|---|---|---|---|---|---|---|---|
| ENSMUSG( | 285.3232 | -0.45038 | 0.126255 | -3.56724 | 0.000361 | 0.034719 | ENSMUSG( | Snx11 |
| ENSMUSG( | 586.9766 | -0.4379 | 0.127698 | -3.42919 | 0.000605 | 0.045874 | ENSMUSG( | Auts2 |
| ENSMUSG( | 9450.766 | -0.41761 | 0.101594 | -4.11065 | 3.95E-05 | 0.006127 | ENSMUSG( | Lrp1 |
| ENSMUSG( | 663.1662 | -0.40583 | 0.102486 | -3.9598 | 7.50E-05 | 0.009727 | ENSMUSG( | Thra |
| ENSMUSG( | 1461.161 | -0.38037 | 0.088807 | -4.28316 | 1.84E-05 | 0.003459 | ENSMUSG( | Piezo1 |
| ENSMUSG( | 912.763 | -0.37761 | 0.085644 | -4.40902 | 1.04E-05 | 0.002292 | ENSMUSG( | Xylt1 |
| ENSMUSG( | 396.0801 | -0.37563 | 0.109857 | -3.41928 | 0.000628 | 0.046188 | ENSMUSG( | Fam160b2 |
| ENSMUSG( | 2159.388 | -0.37342 | 0.096118 | -3.88502 | 0.000102 | 0.012619 | ENSMUSG( | Sh3pxd2a |
| ENSMUSG( | 1689.244 | -0.37286 | 0.085306 | -4.37085 | 1.24E-05 | 0.002651 | ENSMUSG( | Nfat5 |
| ENSMUSG( | 614.2891 | -0.36712 | 0.09726 | -3.77461 | 0.00016 | 0.018161 | ENSMUSG( | Scd2 |
| ENSMUSG( | 1584.445 | -0.3626 | 0.081136 | -4.46907 | 7.86E-06 | 0.001926 | ENSMUSG( | Wwp2 |
| ENSMUSG( | 621.7867 | -0.36198 | 0.101625 | -3.56188 | 0.000368 | 0.035089 | ENSMUSG( | Sdc4 |
| ENSMUSG( | 937.809 | -0.36092 | 0.09767 | -3.69529 | 0.00022 | 0.023414 | ENSMUSG( | Sh3pxd2b |
| ENSMUSG( | 1413.993 | -0.36017 | 0.087241 | -4.12848 | 3.65E-05 | 0.005815 | ENSMUSG( | Leng8 |
| ENSMUSG( | 1016.019 | -0.35144 | 0.103245 | -3.40393 | 0.000664 | 0.048298 | ENSMUSG( | Nav1 |
| ENSMUSG( | 1138.427 | -0.34899 | 0.099896 | -3.49352 | 0.000477 | 0.040516 | ENSMUSG( | Map1b |
| ENSMUSG( | 125075.1 | -0.33656 | 0.080478 | -4.18195 | 2.89E-05 | 0.004847 | ENSMUSG( | Malat1 |
| ENSMUSG( | 1919.493 | -0.32856 | 0.089199 | -3.68339 | 0.00023 | 0.024328 | ENSMUSG( | Snrnp70 |
| ENSMUSG( | 811.0221 | -0.32678 | 0.095402 | -3.4253 | 0.000614 | 0.046188 | ENSMUSG( | Syne3 |
| ENSMUSG( | 5427.738 | -0.32335 | 0.090311 | -3.58036 | 0.000343 | 0.033458 | ENSMUSG( | Kmt2d |
| ENSMUSG( | 665.6435 | -0.30693 | 0.083521 | -3.6749 | 0.000238 | 0.024942 | ENSMUSG( | Ankrd52 |
| ENSMUSG( | 780.5651 | -0.30446 | 0.087532 | -3.47833 | 0.000505 | 0.042031 | ENSMUSG( | Gigyf1 |
| ENSMUSG( | 1548.492 | -0.30192 | 0.087098 | -3.46638 | 0.000528 | 0.042519 | ENSMUSG( | Igf2r |
| ENSMUSG( | 1119.772 | -0.28993 | 0.08003 | -3.62275 | 0.000291 | 0.028872 | ENSMUSG( | C4b |
| ENSMUSG( | 1097.453 | -0.28766 | 0.083126 | -3.46047 | 0.000539 | 0.04266 | ENSMUSG( | Abcc5 |
| ENSMUSG( | 2912.271 | -0.26657 | 0.062473 | -4.26695 | 1.98E-05 | 0.003613 | ENSMUSG( | Nisch |
| ENSMUSG( | 2232.521 | -0.24755 | 0.069606 | -3.55648 | 0.000376 | 0.035548 | ENSMUSG( | Kmt2a |
| ENSMUSG( | 2591.547 | -0.2412 | 0.069033 | -3.49389 | 0.000476 | 0.040516 | ENSMUSG( | Tnrc18 |

**Upregulated**

| Row.name: | baseMean | log2FoldCh | lfcSE | stat | pvalue | padj | Ensembl | Gene |
|---|---|---|---|---|---|---|---|---|
| ENSMUSG( | 72.71543 | 0.939409 | 0.214978 | 4.369798 | 1.24E-05 | 0.002651 | ENSMUSG( | Nyap2 |
| ENSMUSG( | 260.0787 | 0.419431 | 0.120745 | 3.473694 | 0.000513 | 0.042483 | ENSMUSG( | Scarna6 |
| ENSMUSG( | 389.2302 | 0.705188 | 0.125855 | 5.603191 | 2.10E-08 | 1.47E-05 | ENSMUSG( | Ifi209 |
| ENSMUSG( | 103.3895 | 0.723733 | 0.20954 | 3.453922 | 0.000552 | 0.043167 | ENSMUSG( | Kcnk2 |
| ENSMUSG( | 1524.153 | 0.692066 | 0.133285 | 5.19239 | 2.08E-07 | 0.0001 | ENSMUSG( | Ube2l6 |
| ENSMUSG( | 2211.358 | 0.918002 | 0.173442 | 5.29284 | 1.20E-07 | 6.06E-05 | ENSMUSG( | Prg2 |
| ENSMUSG( | 125.1019 | 4.273046 | 0.939768 | 4.546918 | 5.44E-06 | 0.001443 | ENSMUSG( | Gm25189 |
| ENSMUSG( | 2120.733 | 0.334472 | 0.095855 | 3.48936 | 0.000484 | 0.040603 | ENSMUSG( | B2m |
| ENSMUSG( | 90.71621 | 1.210964 | 0.21585 | 5.61022 | 2.02E-08 | 1.47E-05 | ENSMUSG( | Angpt4 |
| ENSMUSG( | 6138.315 | 0.504201 | 0.115643 | 4.359996 | 1.30E-05 | 0.002707 | ENSMUSG( | Alas2 |
| ENSMUSG( | 2763.624 | 0.688754 | 0.172121 | 4.001557 | 6.29E-05 | 0.008511 | ENSMUSG( | Ctsk |
| ENSMUSG( | 163.8932 | 0.678857 | 0.185468 | 3.660232 | 0.000252 | 0.02576 | ENSMUSG( | Gm20628 |
| ENSMUSG( | 304.8098 | 0.553017 | 0.162645 | 3.400148 | 0.000673 | 0.048488 | ENSMUSG( | Bank1 |
| ENSMUSG( | 141.3237 | 0.589415 | 0.172329 | 3.420283 | 0.000626 | 0.046188 | ENSMUSG( | Clca3a1 |
| ENSMUSG( | 169.8427 | 0.707365 | 0.170466 | 4.149591 | 3.33E-05 | 0.005371 | ENSMUSG( | Isg15 |
| ENSMUSG( | 2293.436 | 0.427376 | 0.120868 | 3.535878 | 0.000406 | 0.037317 | ENSMUSG( | Bpgm |

| | | | | | | | | |
|---|---|---|---|---|---|---|---|---|
| ENSMUSG( | 1721.544 | 0.478387 | 0.123708 | 3.86706 | 0.00011 | 0.013453 | ENSMUSG( | Snca |
| ENSMUSG( | 64.63797 | 1.654299 | 0.343514 | 4.815806 | 1.47E-06 | 0.000527 | ENSMUSG( | Igkv1-117 |
| ENSMUSG( | 188.0048 | 2.161734 | 0.593068 | 3.645 | 0.000267 | 0.026908 | ENSMUSG( | A2m |
| ENSMUSG( | 158.7232 | 0.507453 | 0.146356 | 3.467243 | 0.000526 | 0.042519 | ENSMUSG( | BC035044 |
| ENSMUSG( | 60.86661 | 1.19458 | 0.317561 | 3.761736 | 0.000169 | 0.018619 | ENSMUSG( | Pde6h |
| ENSMUSG( | 54.45016 | 0.843126 | 0.243138 | 3.467693 | 0.000525 | 0.042519 | ENSMUSG( | Siglecf |
| ENSMUSG( | 10604.1 | 0.656452 | 0.084128 | 7.803046 | 6.04E-15 | 3.19E-11 | ENSMUSG( | Hbb-bs |
| ENSMUSG( | 98.18261 | 1.26311 | 0.306531 | 4.120662 | 3.78E-05 | 0.00594 | ENSMUSG( | Shank2 |
| ENSMUSG( | 211.4813 | 1.345268 | 0.22536 | 5.969427 | 2.38E-09 | 2.30E-06 | ENSMUSG( | Ear1 |
| ENSMUSG( | 246.8385 | 1.219158 | 0.210404 | 5.794367 | 6.86E-09 | 6.16E-06 | ENSMUSG( | Ear2 |
| ENSMUSG( | 277.982 | 1.32449 | 0.209641 | 6.317903 | 2.65E-10 | 3.34E-07 | ENSMUSG( | Ear6 |
| ENSMUSG( | 1759.898 | 0.405591 | 0.100704 | 4.027535 | 5.64E-05 | 0.007878 | ENSMUSG( | Bnip3l |
| ENSMUSG( | 210.3388 | 0.646299 | 0.18196 | 3.551874 | 0.000382 | 0.035906 | ENSMUSG( | Trpc6 |
| ENSMUSG( | 2101.389 | 0.795857 | 0.168209 | 4.731366 | 2.23E-06 | 0.000738 | ENSMUSG( | Acp5 |
| ENSMUSG( | 142.7842 | 0.747502 | 0.198549 | 3.76482 | 0.000167 | 0.018619 | ENSMUSG( | Pknox2 |
| ENSMUSG( | 187.1311 | 1.936541 | 0.426131 | 4.544471 | 5.51E-06 | 0.001443 | ENSMUSG( | Aldh1a2 |
| ENSMUSG( | 53495.76 | 0.313191 | 0.089031 | 3.517759 | 0.000435 | 0.03894 | ENSMUSG( | Rn7sk |
| ENSMUSG( | 28724.31 | 0.775014 | 0.152602 | 5.078651 | 3.80E-07 | 0.000171 | ENSMUSG( | Hba-a1 |
| ENSMUSG( | 12924.96 | 0.521662 | 0.11368 | 4.588871 | 4.46E-06 | 0.001246 | ENSMUSG( | Hba-a2 |
| ENSMUSG( | 157.1411 | 1.032587 | 0.183775 | 5.618773 | 1.92E-08 | 1.47E-05 | ENSMUSG( | Alox15 |
| ENSMUSG( | 287.9708 | 0.733063 | 0.156279 | 4.69072 | 2.72E-06 | 0.000835 | ENSMUSG( | Cd79b |
| ENSMUSG( | 1272.39 | 0.514759 | 0.128104 | 4.018303 | 5.86E-05 | 0.008103 | ENSMUSG( | Hist1h2ai |
| ENSMUSG( | 2034.238 | 0.529787 | 0.130993 | 4.044399 | 5.25E-05 | 0.007498 | ENSMUSG( | Hist1h4k |
| ENSMUSG( | 876.7077 | 0.544005 | 0.131055 | 4.150959 | 3.31E-05 | 0.005371 | ENSMUSG( | Hist1h3i |
| ENSMUSG( | 268.4439 | 2.406167 | 0.189298 | 12.71103 | 5.14E-37 | 6.46E-33 | ENSMUSG( | Hist1h4m |
| ENSMUSG( | 585.6021 | 0.543535 | 0.154068 | 3.527896 | 0.000419 | 0.038182 | ENSMUSG( | Hist1h2ag |
| ENSMUSG( | 4456.294 | 0.392724 | 0.111867 | 3.510633 | 0.000447 | 0.039324 | ENSMUSG( | Hist1h4d |
| ENSMUSG( | 1276.107 | 0.550065 | 0.132074 | 4.164808 | 3.12E-05 | 0.005158 | ENSMUSG( | Hist1h2ac |
| ENSMUSG( | 1211.849 | 0.545946 | 0.160123 | 3.409531 | 0.000651 | 0.047591 | ENSMUSG( | Hist1h3c |
| ENSMUSG( | 3140.158 | 0.439427 | 0.112454 | 3.907612 | 9.32E-05 | 0.011725 | ENSMUSG( | Hist1h2ab |
| ENSMUSG( | 2539.944 | 0.500074 | 0.13225 | 3.781276 | 0.000156 | 0.017859 | ENSMUSG( | Hist1h3b |
| ENSMUSG( | 213.7854 | 0.688603 | 0.142865 | 4.819949 | 1.44E-06 | 0.000527 | ENSMUSG( | Esm1 |
| ENSMUSG( | 91.46839 | 1.448029 | 0.342655 | 4.225905 | 2.38E-05 | 0.004158 | ENSMUSG( | Greb1 |
| ENSMUSG( | 1572.997 | 0.448156 | 0.130948 | 3.422393 | 0.000621 | 0.046188 | ENSMUSG( | Rsad2 |
| ENSMUSG( | 228.9894 | 0.478431 | 0.138448 | 3.455681 | 0.000549 | 0.043154 | ENSMUSG( | Ndufb1-ps |
| ENSMUSG( | 676.8793 | 0.525276 | 0.135986 | 3.862707 | 0.000112 | 0.013563 | ENSMUSG( | Ckb |
| ENSMUSG( | 326.2041 | 2.198523 | 0.403643 | 5.446697 | 5.13E-08 | 3.23E-05 | ENSMUSG( | Ighg2c |
| ENSMUSG( | 1010.89 | 3.323031 | 0.457894 | 7.257201 | 3.95E-13 | 1.24E-09 | ENSMUSG( | Ighg2b |
| ENSMUSG( | 167.0487 | 3.292582 | 0.580072 | 5.676158 | 1.38E-08 | 1.16E-05 | ENSMUSG( | Ighg1 |
| ENSMUSG( | 87.8676 | 2.263823 | 0.540685 | 4.186953 | 2.83E-05 | 0.004806 | ENSMUSG( | Ighg3 |
| ENSMUSG( | 195.775 | 0.570848 | 0.168437 | 3.389093 | 0.000701 | 0.049556 | ENSMUSG( | Il7r |
| ENSMUSG( | 143.9671 | 0.943755 | 0.270029 | 3.495015 | 0.000474 | 0.040516 | ENSMUSG( | Iglc2 |
| ENSMUSG( | 84.80843 | 1.044707 | 0.227297 | 4.596216 | 4.30E-06 | 0.00123 | ENSMUSG( | Kng1 |
| ENSMUSG( | 2110.797 | 0.389113 | 0.104164 | 3.735573 | 0.000187 | 0.020486 | ENSMUSG( | Gda |
| ENSMUSG( | 1217.622 | 0.387443 | 0.086254 | 4.491883 | 7.06E-06 | 0.001812 | ENSMUSG( | Btaf1 |
| ENSMUSG( | 596.1891 | 0.515983 | 0.119907 | 4.303211 | 1.68E-05 | 0.003309 | ENSMUSG( | Ide |
